# Low-frequency somatic mutations are heritable in tropical trees *Dicorynia guianensis* and *Sextonia rubra*

**DOI:** 10.1101/2023.06.05.543657

**Authors:** Sylvain Schmitt, Patrick Heuret, Valérie Troispoux, Mélanie Beraud, Jocelyn Cazal, Émilie Chancerel, Charlotte Cravero, Erwan Guichoux, Olivier Lepais, João Loureiro, William Marande, Olivier Martin, Gregoire Vincent, Jérôme Chave, Christophe Plomion, Thibault Leroy, Myriam Heuertz, Niklas Tysklind

**Author notes:** Sylvain Schmitt **Email:**. T.L., M.H., and N.T. contributed equally to this work.

## Abstract

Somatic mutations potentially play a role in plant evolution, but common expectations pertaining to plant somatic mutations remain insufficiently tested. Unlike in most animals, the plant germline is assumed to be set aside late in development, leading to the expectation that plants accumulate somatic mutations along growth. Therefore, several predictions were made on the fate of somatic mutations: mutations have generally low frequency in plant tissues; mutations at high frequency have a higher chance of intergenerational transmission; branching topology of the tree dictates mutation distribution; and, exposure to UV radiation increases mutagenesis. To provide new insights into mutation accumulation and transmission in plants, we produced two high-quality reference genomes and a unique dataset of 60 high-coverage whole-genome sequences of two tropical tree species, *Dicorynia guianensis* (Fabaceae) and *Sextonia rubra* (Lauraceae). We identified 15,066 *de novo* somatic mutations in *D. guianensis* and 3,208 in *S. rubra*, surprisingly almost all found at low frequency. We demonstrate that: 1) low-frequency mutations can be transmitted to the next generation; 2) mutation phylogenies deviate from the branching topology of the tree; and 3) mutation rates and mutation spectra are not demonstrably affected by differences in UV exposure. Altogether, our results suggest far more complex links between plant growth, ageing, UV exposure, and mutation rates than commonly thought.

**Significance Statement:** The origin and fate of new mutations have received less attention in plants than in animals. Similarly to animals, plant mutations are expected to accumulate with growth and time, and under exposure to UV light. However, contrary to animals, plant reproductive organs form late in an individual’s development, allowing the transmission to the progeny of mutations accumulated along growth. Here, we resequenced DNA from different branches differentially exposed to sunlight of two tropical tree species. We showed that new mutations are generally rare in plant tissues and do not mimic branching patterns but can nevertheless be transmitted to the progeny. Our findings provide a new perspective on heritable plant mutation and its pivotal role as the engine of evolution.

---

The Weismann theory (1) states that hereditary traits are transmitted exclusively from the germline. The theory is valid in most animals (2) where germline cells are set aside early in development (1). In plants, germline segregation is generally assumed to occur late in development (3-4 but see 5), which leads to several predictions on the fate of somatic mutations occurring in plant tissues: mutations have generally low frequency in plant tissues (6); mutations at high frequency have a higher chance of intergenerational transmission; branching topology of the tree dictates mutation distribution (7); and, exposure to UV radiation increases mutagenesis (8). At present, all these hypotheses, albeit crucial for plant science, have been poorly tested empirically.

To identify a large set of de novo plant somatic mutations, we resequenced 60 samples in total for two tropical tree species, *Dicorynia guianensis* (Amshoff) and *Sextonia rubra* (Mez) van der Werff (Sup. Note A), corresponding to 3 leaves per branch for a total of up to 10 branches per tree, in addition to cambium tissues from the base of the trunk for comparison (Sup. Note B). The branches were selected as growing in either low or high light exposure, getting the benefits of the maximum contrast of forests located near the equator (5°N). Ultraviolet light (UV) exposure was assessed directly at the sampling points and additionally estimated with a canopy transmittance model inferred using terrestrial and drone lidar scans for the *D. guianensis* tree (Sup. Note C). Given that the quality of the reference genomes is known to be a key aspect to ensure accurate mutation detection, we used a combination of high-fidelity reads and optical maps to generate near chromosome-level assemblies for two wild tropical tree species, *D. guianensis* and *S. rubra*. The two genome assemblies differ in size (550 and 991Mb) and in their genomic content for Guanine Cytosine (GC), transposable elements, and genes, with highly heterogeneous patterns along chromosomes in *D. guianensis* vs. relatively homogeneous ones in *S. rubra* (Fig. 1, Sup. Note D). These two new high-quality annotated genomes were used as a reference to detect somatic mutations.

**Figure 1.**
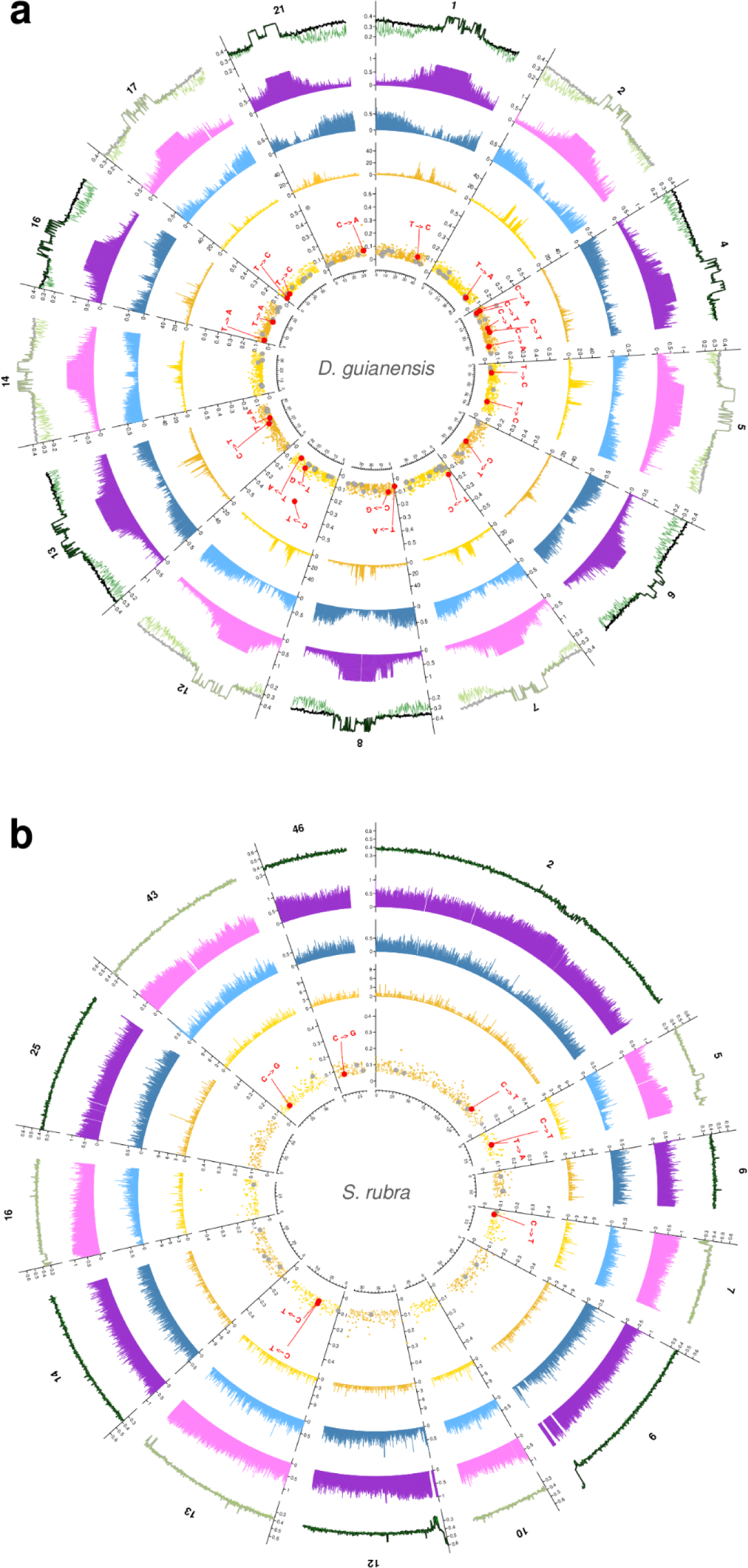
Crown mutations and transmitted mutations in the genomic landscape of the *Dicorynia guianensis* and the *Sextonia rubra* trees’ assembled pseudo-chromosomes. The genomic landscape is similarly portrayed for the two tropical trees: the *Dicorynia guianensis* tree (**a**), and the *Sextonia rubra* tree (**b**). The first (most external) track represents the percentage of Guanine Cytosine (GC) in the whole genome with the black line and in the transposable elements with the green line. The second (least external) track represents the percentage of transposable elements (TE) with purple bars. The third track (middle) represents the percentage of genes with blue bars. The fourth (least internal) track represents the number of somatic mutations detected in the tree crown with yellow bars. The number of somatic mutations correlates with genomic landscapes in *D. guianensis*, the species exhibiting a higher genomic heterogeneity in terms of percentage of genes and TEs (Poisson regression, percentage of TEs b=−0.37(0.04), p<1.10-16, percentage of genes b=-2.31(0.15), p<1.10-16), whereas this is not always significant in *S. rubra* (Poisson regression, percentage of TEs b=−0.62(0.10), p<1.10-9, percentage of genes b=−0.31(0.18), p=0.746). The fifth (innermost) track represents the allelic fraction of the somatic mutations detected in the crown in yellow, the mutations tested for transmission in grey, and the mutations found transmitted to the embryos in red. The inner labels indicate the type of mutations for somatic mutations transmitted to embryos. All measurements are calculated in non-overlapping windows of 100 kb. A ruler is drawn on each pseudo-chromosome, with tick marks every 2 Mb. The genome heterozygosities estimated with K-mer distributions were high for both species, at 0.9% for *D. guianensis* and at 0.7% for *S. rubra*.

Using a mutation detection methodology initially developed for human cancer mutations (9) and later adapted to plants (6), we identified 15,066 unique somatic mutations in *D. guianensis* and 3,208 in *S. rubra*. Only a few were restricted to a single branch (5-9%, Fig 2a-b, Sup. Note E), whereas most mutations were shared by at least two branches whose nearest shared branching point was the base of the crown (43-72%), thus originating below the base of the crown. We further tested the correspondence between the topology of the physical tree and the phylogenies obtained from the somatic mutations and found no correspondence (Fig. 2c-d, Sup. Note F). These results challenge the expectation in plants that the distribution of mutations corresponds to the branching topology of the tree following the growth of the shoot apical meristems (7). We also found no difference in the number of mutations, the type of mutations (nucleotide changes) or the mutation spectra (mutation context with 5’ and 3’ amino acids) between the branches exposed to high vs. low light conditions (Fig. 2e-f, Sup. Note G), which suggests a shielding from UVs in the bud layers (10).

**Figure 2.**
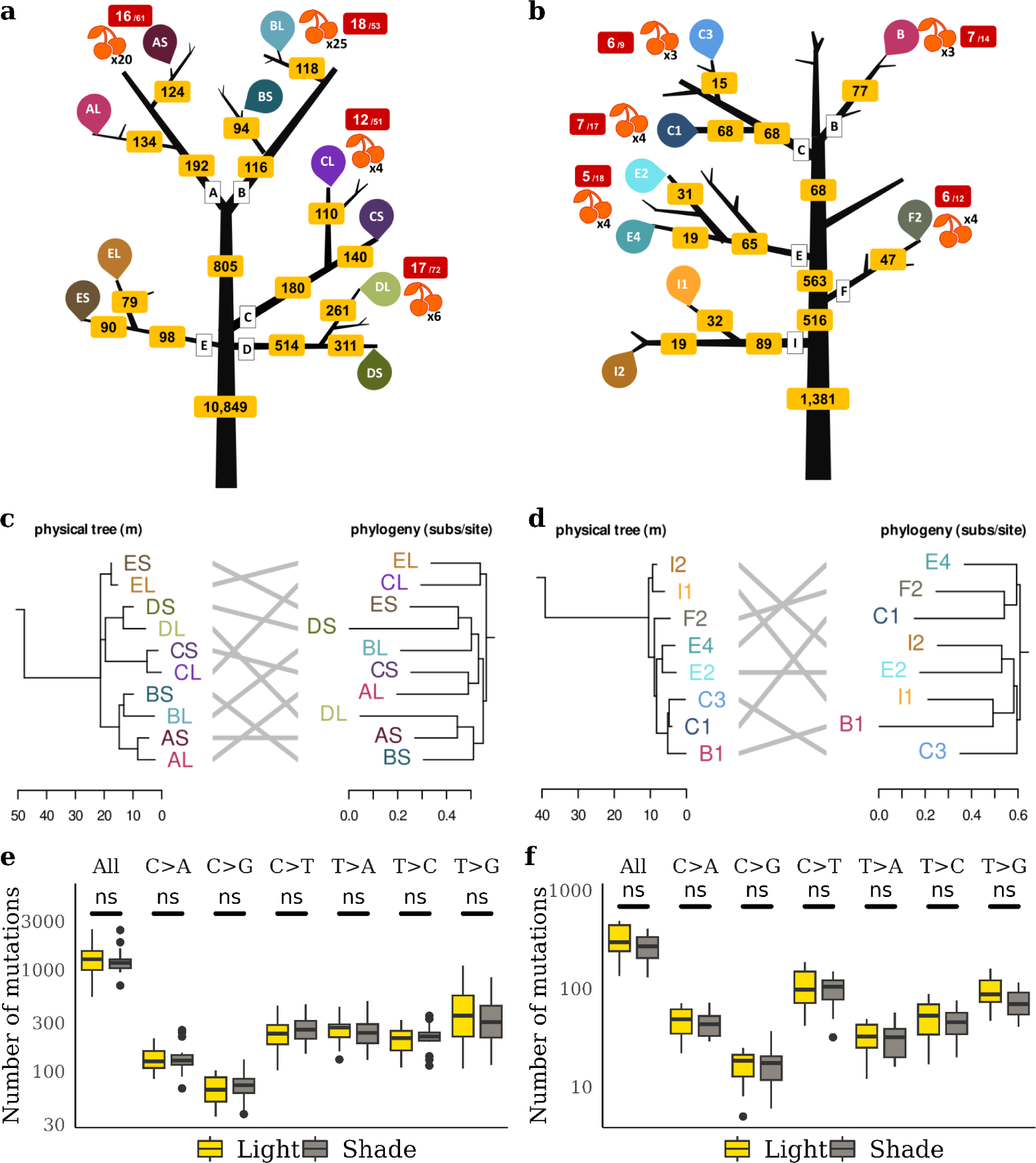
Distributions of somatic mutations through branching topology of the tree, phylogenies, and with light. The distributions of somatic mutations through physical trees, phylogenies, and with light are similarly shown for the two tropical trees: the *Dicorynia guianensis* tree (**a**,**c**,**e**), and the *Sextonia rubra* tree (**b**,**d**,**f**). (**a-b**) The branching topology of the tree is shown in black with the branch names in white boxes. The number of somatic mutations through the crown is indicated in the yellow boxes before the original branching event. The sampling points of three leaves in the light-exposed branches (“L” in letter codes, light colours) and in the shaded branches (“S”, dark colours) are indicated with unique letter codes and coloured drop symbols. Fruit sampling points are represented by red fruits, with the number of fruits sampled indicated in black. The red boxes with white labels indicate the transmission of mutations to fruit embryos out of the total number of mutations tested. (**c-d**) A side-by-side comparison of the physical tree (left, branch length in metres) and the maximum likelihood phylogeny of mutations (right, branch length in substitutions per site). The letters on the ends of the branches indicate the sampling points shown in (**a-b**). (**e-f**) Different mutagens may cause specific mutation types, *i.e.*, changing from base X to base Y (X>Y). The effect of light exposure on the accumulation of somatic mutations as a function of mutation type (X>Y) is represented in yellow and grey boxes. The yellow boxes represent the number of mutations accumulated in all leaves of light-exposed branches and the grey boxes in all leaves of shaded branches. Boxplots show the median (centre line), upper and lower quartiles (box limits), 1.5x interquartile range (whiskers), and outliers (points). The “ns” labels indicate non-significant differences in Student’s T-tests (two-sided). Mutation types include all mutations and all types of transitions and transversions. The y-axis was logarithmically scaled to facilitate reading of low values.

As compared to previous reports about somatic mutations in plants (4,10,11), we have detected far more mutations (ten to hundred times more). This discrepancy is likely associated with the methodology (6), since the vast majority of identified somatic mutations had a low allelic fraction, *i.e.* the fraction of genomic reads with the mutation, which indicates the frequency of mutated cells in the analysed sample (Fig. 3a-b, Sup. Note H). The higher total number of mutations detected in *D. guianensis* can be explained by an enrichment in low fraction mutations in the *D. guianensis* tree detected through deeper sequencing (Supplementary Fig. E1), because increasing the number of reads of a genomic region increases the chances of finding a mutation present in only a few cells of a sample. We generalised the result of the predominance of low fraction mutations in two pedunculate oaks (4,10), and a dataset from one tortuous beech *Fagus sylvatica* L. using the same methodology (Fig. 3c). We then considered mutations at a high allelic fraction (>0.25), a category of mutations for which methodological differences are expected to have a limited impact. The two tropical trees had 3 and 6 somatic mutations with allelic fraction>0.25, as compared to 56-421 somatic mutations for the reanalysed oaks and beech trees from temperate regions (Fig. 3c). Future large-scale investigations are needed to properly test whether there is a difference in mutation rates between temperate and tropical trees, after accounting for the phylogenetic signal, differences in tree age, among other null hypotheses. Overall, our results suggest that low-frequency mutations account for the vast majority of within-individual somatic diversity in plants (for all species, >90% with f<0.25).

**Figure 3.**
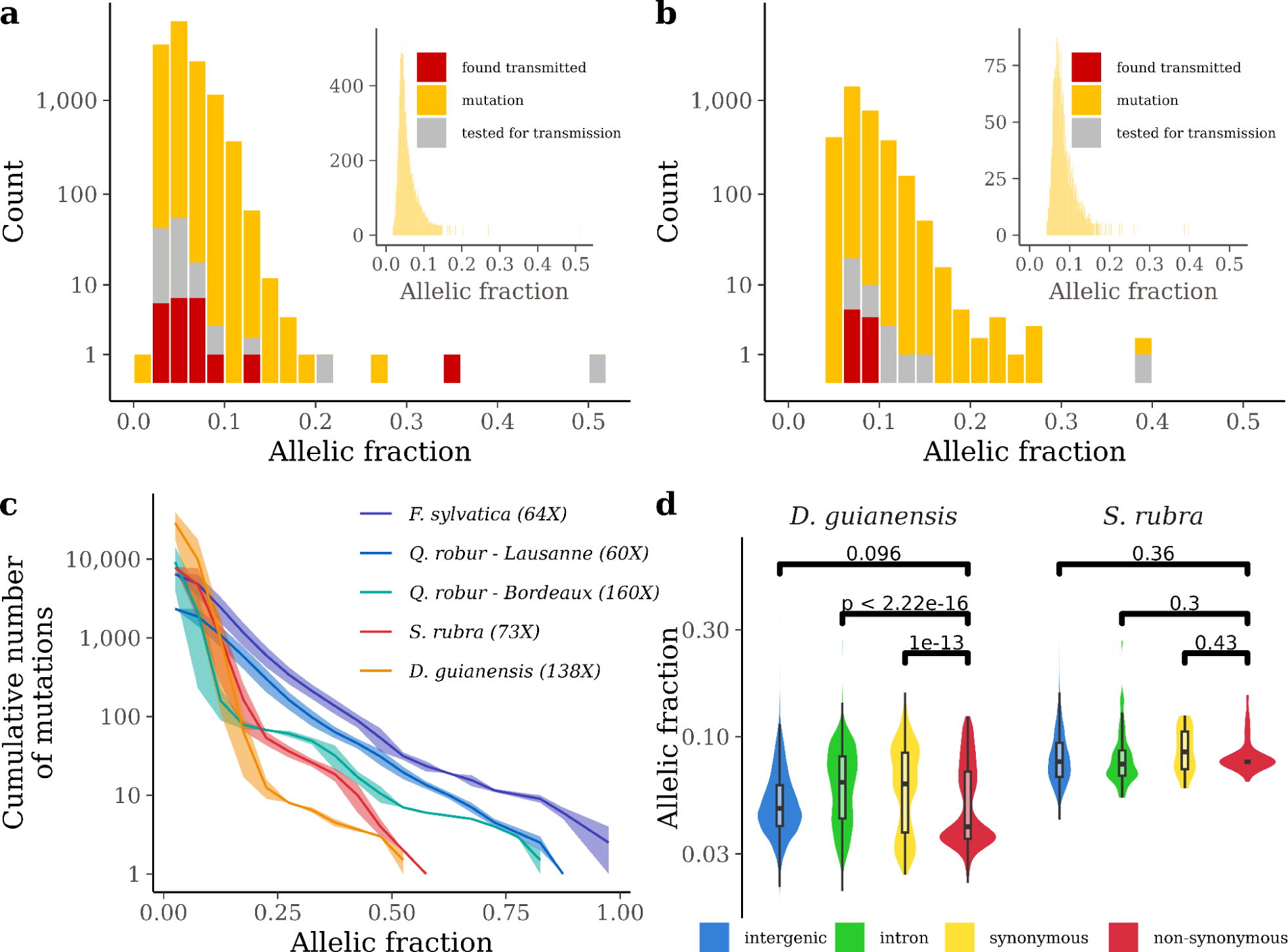
Allelic fractions of somatic mutations among trees and among genomic elements. Histogram of allelic fractions of mutations detected in the crown of the two tropical trees: the *Dicorynia guianensis* tree (**a**), and the *Sextonia rubra* tree (**b**). The main histograms show the allelic fractions of the somatic mutations using a bin of 0.02 and a log-transformed count with the mutations detected in the crown in yellow, the mutations tested for transmission in grey, and the mutations found transmitted to the embryos in red. The histograms in insets show the allelic fractions of the somatic mutations using a bin of 0.001 and a natural count. (**c**) Cumulative number of somatic mutations per branch with decreasing allelic fraction for five trees reanalysed with the same pipeline. The five trees include the two tropical trees studied, i.e. the *Dicorynia guianensis* tree in orange and the *Sextonia rubra* tree in red, and three temperate trees, two pedunculate oaks *Quercus robur* L. from Bordeaux in green and Lausanne in blue and a tortuous phenotype of common beech *Fagus sylvatica* L. in purple. All trees were analysed with the same pipeline (see methods) but were sequenced with a different depth indicated in brackets. The line represents the median value while the area represents the minimum and maximum values on the 2 to 10 branches per tree. (**d**) Comparisons of allelic fractions for non-synonymous mutations in red with synonymous mutations in yellow, intronic mutations in green and intergenic mutations in blue for the two tropical trees: the *Dicorynia guianensis* tree (left panel), and the *Sextonia rubra* tree (right panel). Boxplots show the median (centre line), upper and lower quartiles (box limits), 1.5x interquartile range (whiskers), and outliers (points). The p-value above the bars indicates the significance of the Student’s T-test (two-sided) for the pairs of groups.

The origin of the somatic mutations’ spatial distribution in the physical tree lies in the functioning of the shoot apical meristems. Shoot apical meristems divide either symmetrically into two stem cells or asymmetrically into one stem cell and one differentiated cell (12), resulting in the three-dimensional spatial distribution of stem cells and the somatic mutations they carry during tree growth. In dicots, the layered structure of shoot apical meristems limits cell movement through the prevalence of anticlinal cell divisions, which favours the retention of mutated cell clones, *e.g.* in the form of stable periclinal chimaeras (13). This mechanism could lead to sectoral chimerism through somatic mutations, which may explain both the discrepancy between the physical tree and phylogeny (Fig. 2c-d, 12) and the prevalence of numerous low-frequency somatic mutations (Fig. 3a-b).

Somatic mutations are often viewed as a source of within-tree adaptive variation (14). To test this hypothesis, we investigated whether non-synonymous somatic mutations exhibit differences in allelic fraction as compared to synonymous ones or to non-coding regions. Higher, or lower fractions would be evidence for positive, or negative selection, respectively. For both species, we detected that the average allelic fraction at non-synonymous sites was lower than those at synonymous sites (Fig. 3d). This difference is highly significant in *D. guianensis* (Student’s t-test, p-value<10^-13^) but not significant in *S. rubra* (p=0.43), likely because a limited number of mutations was detected (31 synonymous and 9 non-synonymous mutations). All together, these results are consistent with the intra-organismal purifying selection of non-synonymous mutations, as also observed in seagrass (14), supporting that far more *de novo* mutations are detrimental than beneficial. Until now, low-frequency somatic mutations have been neglected because they were assumed not to be transmitted, and therefore to have no evolutionary future. We explored the transmission of somatic mutations to the next generation through their redetection in the embryos of developing fruits. We used amplicon resequencing for 160 candidate mutations highly shared between sampled leaves and branches, including low-frequency mutations. Using stringent quality filters (Sup. Note I), we demonstrated the transmission of 23 out of 160 tested mutations to embryos in *D. guianensis* and 9 out of 36 in *S. rubra* (Fig. 1). The transmitted mutations were found in several branches of the *D. guianensis* tree but in only one branch of the *S. rubra* tree (Fig. 2). Surprisingly, almost all the mutations for which we found empirical support for their transmission were at low frequency within the plant. Consistently, we observed that the distributions of the allelic fraction of the transmitted mutations (red bars in Fig. 3a-b) were similar to the distributions of the allelic fraction of all mutations in the crown of the trees (yellow bars in Fig. 3a-b, two-sided Student’s t-Test t=1.41 [−0.40, 0.07], df=22, p=0.17 for *D. guianensis* and t = −0.34 [−0.12, 0.09], df=8, p=0.07 for *S. rubra*), resulting in all transmitted mutations having low allelic fractions. By using only mutations with high empirical variant scores, we validated the robustness of our results, namely the abundance of low-frequency mutations, their transmission, the lack of correspondence between mutation phylogenies and the topology of the tree and the absence of a spectrum associated with UV (Sup. Note J). Hence, we found that low-frequency somatic mutations are heritable and thus contribute to increased within-species diversity, which challenges current tacit assumptions that only high-frequency mutations would matter for evolution. Despite their low frequency and scarcity across the genome, low-frequency somatic mutations could substantially contribute to standing genetic variation, which is the engine of evolution (Sup. Note K). We therefore call for a new view on somatic mutations in plants with renewed assumptions: (i) the distribution of somatic mutations does not necessarily correspond to the branching topology of the tree, (ii) most somatic mutations are low-frequency mutations, and (iii) low-frequency mutations can be transmitted to embryos in trees. Our results are consistent with far more complex links between growth, ageing and mutation rates than commonly thought in plants, along the lines of recent empirical evidence in animals (2,15).

## Materials and Methods

### Choice of species and individuals

The study was conducted in the Amazon forest, in the coastal forests of French Guiana. A database of 710 tree species containing available information on the presence of tree rings, maximum diameter at breast height, architectural type, reproductive phenology and ecological and economic importance was constructed. A set of 15 candidate species was selected, and their genome size was estimated by flow cytometric analyses. On this basis, we chose to work on *Dicorynia guianensis* (Amshoff) and *Sextonia rubra* (Mez) van der Werff, which are common in French Guiana, and are ecologically and economically important species. We selected large-stature trees above 40 metres without signs of dieback or senescence to maximise the potential for mutations with an increased number of cell divisions. The architecture of the trees was studied with binoculars and by climbing to select trees where in each bough we could sample pairs of branches with contrasting light exposure. We finally selected a *D. guianensis* tree in the Saint George area (4°01’N, 51°59’W), which has an annual rainfall of 3,665 mm and a mean air temperature of 27°C, and a *S. rubra* tree near the Paracou research station (5°18’N, 52°53’W), which has an average annual rainfall of 3,041 mm and a mean air temperature of 25.7°C (Sup. Note A).

### Sampling and tree structure

On 13 October 2020, we sampled the *S. rubra* tree: three cambium tissues at the base of the trunk, about 1.3 m above the ground and equidistant around the perimeter of the trunk and three leaves from the same twig per branch for a total of eight branches were sampled. The branches were selected in three pairs, each from a different bough, plus two independent branches from two other boughs. Fruits were sampled from 5 different branches where leaves had been collected. On 22 April 2021, we sampled the *D. guianensis* tree: three cambium tissues at the base of the trunk, about 1.3 m above the ground and equidistant from the perimeter of the trunk and three leaves from a single twig per branch for a total of ten branches were sampled. The branches were selected in five pairs, each from a different bough. Fruits were sampled from 5 different branches where leaves had been collected. On 13 October 2020 and 15, 16 and 22 November 2021, we described the structure of both trees: branching patterns were recorded and all branch lengths, as well as basal and terminal diameters, were measured for branches with a basal diameter greater than 10 cm, in addition to the trunk. On 21, 22 and 23 March 2022, 30 wood cores were collected with a drill in branches from throughout the crown and in the trunk of *D. guianensis*. In both species, leaf, cambium, and fruit samples were frozen in liquid nitrogen and stored at −80 °C until DNA and RNA extraction (Sup. Note B).

### Characterisation of light conditions

A linear photosynthetically active radiation (PAR) ceptometer (AccuPar, Decagon Devices, Pullman, WA, USA) was used at each sampling position on both trees during sampling to measure direct incident light in the 400-700 nm wavelength range around noon in comparison to open incident light measured on the nearest road. A ground (TLS) and drone (DLS) lidar (light detection and ranging) campaign (TLS, Faro Focus3D 120; DLS, Yellowscan Vx20-100) was conducted on 3 May 2021 to map the transmittance of the *D. guianensis* tree canopy. TLS scans were performed horizontally from 0 to 360° and vertically from −60 to 90°, resulting in 174.8 million points per scan for 10 scans in a forest gap near the tree and 4 scans from the nearby road. DLS scans were taken at 35 m above the focal tree in 2 perpendicular flights with flight lines spaced 10 m apart in a circular area 150 m in diameter above the focal tree, resulting in 46.6 million points. Prior to the lidar acquisition, reflective strips were placed on the sample points by tree climbers to detect the sample points in the lidar cloud. AMAPvox software was used to calculate an annual illuminance index from the aerial laser scanning. The plant area density (PAD, m2/m3) was calculated for the focal tree in context (with a diameter of 30 m around the tree) using 1 m3 voxels. An estimate of the annual proportion of solar radiation above the canopy received at the sample point was then simulated considering a brightness index of 0.5 and a latitude of 5 degrees. The uncertainty in transmittance due to uncertainties in the location of the sampling point was further assessed by randomly sampling 10 positions around the sampling points to 0.5 metres and revealed small variations in transmittance. The estimates were in agreement with the light/shade classification of branches identified by the tree climbers (Sup. Note C).

### Genome assemblies and annotations

High molecular weight (HMW) DNA was extracted from 0.7 g of three leaves of both individuals using CTAB and isopropanol precipitation before RNAase treatment and bead purification (Doyle and Doyle 1987). High-fidelity (HiFi) genomic reads were produced with two sequencing runs on the PacBio Sequel II system on 2 (*D. guianensis*) to 4 (*S. rubra*) SMRTCells for each run. We obtained 1,898,004 corrected reads for *D. guianensis* (N50=21,233; DP=58.7X), which we assembled into 562 contigs (N50=37.76Mb; L50=8 contigs; GC=37.25%) using the HiFiasm assembler (v0.15.5). Similarly, we obtained 6,624,997 corrected reads for *S. rubra* (N50=17,577; DP=114X), which we assembled into 747 contigs (N50=16.513Mb; L50=17 contigs; GC=38.50%). HMW DNA was also used to produce optical maps to construct hybrid scaffolds with optical reads produced by two passages of Bionano saphyr. For *D. guianensis*, we obtained 54 hybrid scaffolds (N50=38,450Mb; N=0.76%; 571 gaps), while 515 contigs remained unanchored with a total length of 28,784Mb representing 4.97% of the genome. For *S. rubra*, we obtained 35 hybrid scaffolds (N50=60.458Mb; N=1%; 1.923 gaps), while 609 contigs remained unanchored with a total length of 53.067Mb representing 5.08% of the genome. Genome quality was evaluated using BUSCO and Merqury. We constructed an automated genome annotation workflow that performs: (i) *de novo* and known transposable element (TE) detection, (ii) *de novo* and known gene models detection, and (ii) functional gene annotation. *De novo* TE detection uses RepeatModeler2 (v2.0.3) followed by classification using RepeatClassifier (v2.0.3) and TEclass (v2.1.3). The de novo TEs obtained were merged with the known TE accessions for Viridiplantae from RepBase (v27.07). This consolidated database is used for TE detection in each genome prior to soft masking using RepeatMasker (v2.0.3). Detection of de novo and known gene models is based on BRAKER2 and its dependencies. Finally, functional annotation of candidate genes is based on the Trinotate (v3.2.1) pipeline using TransDecoder (v5.5.0), TMHMM, HMMER, BLAST (v2.13.0), RNAmmer (v1.2), and SignalP (v4.1) with UniProt and Pfam databases (Sup. Note D).

### Leaf and cambium mutation detection

Genomic DNA was extracted from 30 mg of frozen leaf or cambium tissue per sample point for both trees using a CTAB protocol with chloroform-isoamyl alcohol extraction (24:1), isopropanol precipitation and resuspension of the pellet in 1x Low TE (10 mM Tris-HCl + 0.1 mM EDTA). DNA was quantified using a Qubit HS assay (Thermo Fisher Scientific, Waltham, MA, USA) and purified with AMPure XP beads (Beckman Coulter Genomics, Danvers, MA, USA) where necessary to allow library preparation. An Illumina sequencing library was produced for each leaf using an optimised NEBNext Ultra II DNA library protocol (New England Biolabs, Ipswich, MA, USA). Libraries were pooled into multiplexes after independently labelling each library prior to whole genome sequencing (WGS) on an S4 flow cell and NovaSeq 6000 instrument with v1.5 chemistry (2 × 150 PE mode). We obtained 33 cambium and leaf libraries for *D. guianensis* with a sequencing depth of about 160X and 27 libraries with a depth of about 80X for *S. rubra*. We took advantage of a workflow previously developed by some of us to detect somatic mutations from sequencing reads mapped to a genomic reference (6). Paired sequencing reads from each library were quality controlled using FastQC (v0.11.9) before being trimmed using Trimmomatic (v0.39), which retains only paired-end reads without adapters and with a phred score greater than 15 in a 4-base sliding window. The reads are aligned against the reference genome using BWA mem with the option to mark shorter splits (v0.7.17). The alignments are then compressed using Samtools view in CRAM format, sorted by coordinates using Samtools sort, and indexed using Samtools index (v1.10). Duplicate reads in the alignments are marked using GATK MarkDuplicates (v4.2.6.1). Sequencing depth was estimated along the genome using Mosdepth (v0.2.4) globally and over a sliding window of 1 kb. We used Jellyfish (v1.1.12) and GenomeScope to estimate heterozygosity up to 21-mer. We used GATK (HaplotypeCaller, GatherGVCFs, GenomicsDBImport, GenotypeGVCFs) to call heterozygous sites from the previously obtained alignments. We filtered single-nucleotide polymorphisms (SNPs) using bcftools (v1.10.2), GATK VariantFiltration (v4.2.6.1), and plink (v1.90), retaining only biallelic SNPs and discarding those with quality less than 30, quality per depth less than 2, Fisher strand ratio greater than 60, and strand odds ratio greater than three. To eliminate all truly heterozygous sites, we further filtered out SNPs present in all sampled genotypes and tissues (no missing data) and shared by at least all but one tissue. Finally, the workflow uses Strelka2 (v2.9.10) to detect mutations by comparing two samples, a mutated sample and a normal (directional) sample. To detect cambium mutations present at the base of the tree, we compared all potential pairs (6 in total) among the three cambium libraries. To detect leaf mutations, we compared each leaf library to the first cambium library as a reference sample. We filtered candidate leaf mutations discarding previously identified heterozygous sites and all candidate mutations from all cambium comparisons using BEDTools subtract (v2.29.2). We also filtered candidate leaf mutations using the following criteria: (i) no copies of the mutated allele in the reference sample, in this case the cambium sample; (ii) a read depth for both samples between the 5th quantile and the 95th quantile of the corresponding library coverage; and (iii) the presence of the mutation in at least two biological replicates (at least 2 leaves of the crown) We used the same pipeline and compared mutations detected in two pedunculate oaks *Quercus robur* L. (4,10), and in a dataset from a tortuous phenotype of common beech *Fagus sylvatica* (16-17) (Sup. Note E).

### Somatic mutation distributions through physical trees, phylogenies, and with light exposure

We explored mutation distribution along tree architecture by assuming the origin of the mutation in the tree architecture was at the latest the most recent common branching event among all branches harbouring the mutation (11). We further built mutation phylogenies using iqtree rooting the tree with the non-mutated library from the cambium mean genotype without mutations. We compared phylogenies to the physical architecture of both trees with the dendextend R package (Supplementary Note F). We explored the effects of light on the occurrence of mutations in the trees using Student’s T-tests and Kolmogorov-Smirnov tests. We compared the number of mutations detected in branches exposed to high vs. low light conditions using the leaves as an observation. We further compared mutation types (base change) and mutation spectra (mutation context with 5’ and 3’ bases) between high and low light conditions among branches of each tree (Sup. Note G).

### Low-frequency mutations annotation

We explored the allelic fractions of somatic mutations in relation to tree sequencing depth, a known determinant of the sensitivity of somatic mutation detection (8), for the *D. guianensis* tree, the *S. rubra* tree, two pedunculate oaks *Q. robur* (4,10), and data from one tortuous phenotype of common beech *F. sylvatica* (16-17). We further compared mutation annotations in terms of their presence in transposable elements (TE) and genes among trees. We assessed mutation functional impact using SNPeff and related non-synonymous mutations to their functional annotations, gene ontology, and allelic fraction. We finally explored the allelic fraction of mutations depending on genomic contexts using Student’s T-tests (Sup. Note H).

### Detection of fruit mutations

We explored mutation transmission to fruits using amplicon resequencing. We kept as candidate mutations for redetection only mutations present in at least three leaves from the branches that had fruits during sampling for resequencing, which resulted in 160 candidate mutations (124 for *D. guianensis* and 36 for *S. rubra*). Frozen fruits were dissected in 4 tissues: (i) embryo sac, (ii) nucellus, (iii) pericarp, and (iv) fruit base. Genomic DNA was extracted from 10-50 mg of frozen fruit tissue for both trees and additional leaf tissue for positive control with a CTAB protocol with chloroform - isoamyl alcohol (24:1) extraction, isopropanol precipitation and pellet resuspension in 1x Low TE (10 mM Tris-HCl + 0.1 mM EDTA; Doyle and Doyle 1987). DNA was quantified using a Qubit HS assay. Primer3plus was used to design primer pairs targeting candidate mutations (amplicon size between 100 and 200 pb). Only one *D. guianensis* candidate mutation failed to yield a primer pair. Illumina universal tags were added to the 5′ end of the forward and reverse primer sequences respectively. Oligonucleotides were ordered in a plate format from Integrated DNA Technologies (Coralville, IA, USA) with standard desalt purification at 25 nmoles synthesis scale. Each primer pair was tested using simplex PCR amplification of one DNA sample per species in a volume of 10 μL containing 2 μL of 5X Hot Firepol Blend master mix (Solis Biodyne, Tartu, Estonia), 1 µL of 2µM primer pairs, 1 µL of DNA (10 ng/µL), and 6 µL of PCR-grade water. We ran the PCR on a Veriti 96-Well thermal cycler (Applied Biosystems, Waltham, MA, USA) performing an initial denaturation at 95°C for 15 min, followed by 35 cycles of denaturation at 95°C for 20 s, annealing at 59°C for 60 s, extension at 72°C for 30 s, and a final extension step at 72°C for 10 min. We checked the amplification on a 3% agarose gel. A total of 6 *D. guianensis* primer pairs that failed to amplify were discarded at this stage. The remaining 101 *D. guianensis* and 33 *S. rubra* primer pairs, targeting respectively 117 and 36 mutations, were grouped accounting for potential primer dimer formation using PrimerPooler for subsequent multiplex PCR amplification. Four multiplexed PCRs were done for each species in a volume of 10µL using 2 µL of 5X Hot Firepol Multiplex master mix (Solis Biodyne), 1 µL of multiplex primer mix (0.5 µM of each primer), 2 µL of DNA (10 ng/µL), and 5 µL of PCR-grade water. Amplifications were performed on a Veriti 96-Well thermal cycler (Applied Biosystems) using an initial denaturation at 95°C for 12 min followed by 35 cycles of denaturation at 95°C for 30 s, annealing at 59°C for 180 s, extension at 72°C for 30 s, and a final extension step at 72°C for 10 min. The amplicons from the four multiplexed PCRs of each sample were pooled. Illumina (San Diego, CA, USA) adapters and sample-specific Nextera XT index pairs were added to the amplicons by a PCR targeting the Illumina universal tags attached to the locus-specific primers. This indexing PCR was done in a volume of 20 μL using 5X Hot Firepol Multiplex master mix, 5 µL of amplicon, and 0.5 µM of each of the forward and reverse adapters, using an initial denaturation at 95°C for 12 min followed by 15 cycles of denaturation at 95°C for 30 s, annealing at 59°C for 90 s, extension at 72°C for 30 s, and a final extension step at 72°C for 10 min. We then pooled the libraries and purified them with 0.9X AMPure XP beads (Beckman Coulter, Brea, CA, USA). We checked the library quality on a Tapestation 4200 (Agilent Technologies, Santa Clara, CA, USA) and quantified it using QIAseq Library Quant Assay kit (Qiagen, Hilden, Germany) on a LightCycler 480 quantitative PCR (Roche, Basel, Switzerland). The sequencing was done on an Illumina MiSeq sequencer using a V2 flow cell with a 2×150 bp paired-end sequencing kit. Paired-end sequencing reads of each library were trimmed using Trimmomatic (v0.39, Bolger et al., 2014) keeping only paired-end reads without adaptors and a phred score above 15 in a sliding window of 4 bases. Reads were aligned against the reference genome using BWA mem with the option to mark shorter splits (v0.7.17, Li & Durbin, 2009). Alignments were then compressed using Samtools view in CRAM format, sorted by coordinates using Samtools sort, and indexed using Samtools index (v1.10, Li et al., 2009). We used GATK (HaplotypeCaller, GatherGVCFs, GenomicsDBImport, GenotypeGVCFs; Auwera et al., 2013) to call candidate mutations transmitted to fruits from previously obtained alignments. We filtered variants that corresponded to the 160 candidate mutations (124 for the *D. guianensis* tree and 36 for the *S. rubra* tree). For stringency, we highlighted and discussed only candidate mutations that were considered a heterozygous site by the GATK GenotypeGVCFs call (Sup. Note I).

## Data, script and code availability

Genomic and transcriptomic reads from leaf, cambium, and fruits and corresponding genomes are available in GenBank (18). genomeAnnotation and detectMutations pipelines as well as downstream analyses are available on GitHub (19). Results and intermediary files are available on Zenodo (20).

## Acknowledgments

We are grateful to Valentine Alt, Emeline Houël, Laetitia Brechet, Saint-Omer Cazal, and Hadrien Lalagüe for their help with tree climbing and sampling. We thank Olivier Brunaux and Caroline Bedeau for their help in accessing the Regina site and the Office National des Forests data. We are grateful to Nicolas Barbier, Ilona Clocher, and Jean-Louis Smock for their help with lidar acquisition. PacBio HiFi reads were produced at Gentyane Genomic Facility (INRAE Clermont-Ferrand, France). Bionano Saphir reads and genome *de-novo* assembly were produced at INRAE-CNRGV Plant Genomic Centre (Toulouse, France). Whole genome re-sequencing was performed at Genoscope National Sequencing Centre (Evry, France) with the help of Eric Mahieu, Corinne Cruaud, and Pedro H. Oliveira. Sequence-based genotyping was performed at the Bordeaux Genome Transcriptome Facility PGTB (doi:10.15454/1.5572396583599417E12) with the help of Zoé Delporte. We are grateful to the GenoToul bioinformatics facility (Castanet-Tolosan, Toulouse, Occitanie, France, doi:10.15454/1.5572369328961167E12) for providing help, computing and data storage resources The climate data were provided by Météo-France to the Unité Mixte de Recherche EcoFoG for research purposes within the framework of a MétéoFrance-INRAE AgroClim agreement. This study was funded by an ANR Investissement d’Avenir grant: CEBA (ANR-10-LABEX-0025).

## Author Contributions

MH, NT, TL, PH, CP & JC conceived the ideas; VT, JC, NT, PH, and SS conducted the fieldwork; GV and OM characterised individuals’ light conditions; MB, EC, CC, EG, OL, JL, WM, VT, and MH produced genetic data; SS, TL, NT, and MH analysed the data and led the writing of the manuscript. All authors contributed critically to the drafts and gave final approval for publication.

## A - Choice of species and individuals

### Candidate species selection

To study the process of mutations, we wanted to focus on a species that fulfilled a maximum of these criteria:

- Ecological relevance: we aimed to work on common species, so the results of our study would be ecologically relevant.
- Economic relevance: we aimed to work on species where the availability of high quality genomes would contribute to further studies on genomics, evolutionary ecology, and sustainable management of these species.
- Species tree architecture: we aimed to have emergent canopy trees with typical architectures that allowed us to establish a sampling design where pairs of branches with contrasting sun exposure could be sampled on the same bough, and this sampling strategy repeated on different boughs across the whole tree canopy.
- Predictable phenology: we aimed to study species where we could predictably expect fruits for the individual tree, so that we could assess the transmission of mutations from somatic tissues to the embryo tissues.
- Genome size: we aimed to work on species with smaller genomes, which would reduce the amount of sequencing needed to identify low-frequency mutations.

The study was conducted in the Amazon forest, in the coastal forests of French Guiana. A database of 710 species of trees from French Guiana containing information on the functional group, maximum diameter at breast height, architectural model, presence of wood rings, reproductive phenology, and ecological and economical importance was constructed. A set of 15 candidate species was thus selected based on these criteria.

### Genome size estimation

To obtain information on genome size of the candidate species we collected leaf tissue from 15 individuals from the 15 selected species at Paracou Research station, and conserved the tissues in RNAlater (Qiagen) or in silica gel until flow cytometric analyses. Nuclear suspensions were obtained following Galbraith et al. (1983) by chopping RNA-later conserved tissue of the studied species and fresh leaf tissue of *Pisum sativum* ‘Ctirad’ (internal reference standard, 2C = 9.09 pg; Doležel et al. 1998) in 1 ml of WPB buffer (Loureiro et al. 2007). Then, the nuclear suspension was filtered using a 50 µm nylon mesh and 50 µg/ml of propidium iodide (PI, Fluka, Buchs, Switzerland) and 50 µg/ml of RNAse (Fluka, Buchs, Switzerland) were added to stain the nuclear DNA and remove dsRNA, respectively. Samples were analysed in a Sysmex CyFlow Space flow cytometer equipped with a 532 nm green solid-state laser and operating at 30 mW, and results were acquired using FloMax software v2.4d.

Regardless of whether preserved in RNAlater or in silica gel, we were able to obtain reliable haploid genome size estimates ranging from 419 Mbp (in *Laetia procera*) to 1836 Mbp (in *Symphonia globulifera*) (Table A1). According to the genome size categories of Leitch et al. (1998), most of the candidate species present a very small genome size (1C ≤ 1,372 Mbp). Only *Moronobea coccinea* with 1C = 1508 Mbp and *Symphonia globulifera* with 1C = 1,836 Mbp, presented a small genome size. The genome size estimation of *Eschweilera coriacea* obtained here is slightly larger than the one given in Heuertz et al. (2020), but it is in the same range of values as other estimations obtained in the genus (Heuertz et al., 2020).

Based on all of the above information we chose to work on *Dicorynia guianensis* (Amshoff) and *Sextonia rubra* (Mez) van der Werff, which are common in French Guiana, and they are ecologically and economically important species, being respectively the first and second most harvested species in French Guiana. They are shade-tolerant, canopy to emergent trees. *D. guianensis* forms growth-rings (Detienne 1995), while *S. rubra* produces seasonal variation in wood chemistry (Ponton *et al*., 2016). Both species have very small genomes.

### Individual selection

We surveyed *D. guianensis* trees at the Paracou research station and the Office National des Forêts plots between Régina and Saint George (Secteur Saint George). We aimed to find trees as large as possible (DBH > 100cm) but without evidence of dieback or senescence. This was to maximise the potential of mutations, assuming that the number of mutations is correlated with the number of cellular divisions. We avoided senescent trees that may have lost many of their main branches, as well as trees with evidence of damage from nearby treefalls, or trees whose crowns were intermingled with other trees. We examined the architecture of the trees with binoculars and by climbing the trees, and chose trees that would allow an experimental design where several boughs could be selected (bough being the main branch attached to the trunk of the tree), and where in each bough we could sample pairs of branches with contrasting light exposure. Other practical aspects to consider were ease of access from the road to transport equipment and the ease and safety of climbing the trees. We finally settled for a *D. guianensis* tree, hereafter named Angela, in the secteur Saint George (4°01’N, 51°59’W), that complied with all our requirements and was fruiting at the moment of sampling. The secteur Saint George experiences an annual rainfall of 3,665 mm and a mean air temperature of 27°C. The choice of the *Sextonia rubra* individual was dictated by a collaboration with another project, TREE-D, that aims at looking into chemical and holobiont heterogeneity across the whole tree. The tree, hereafter named Sixto, was just outside of the Paracou Research Station (5°18’N, 52°53’W), and complied with the requirements stated for the *D. guianensis* and was fruiting at the moment of sampling. The Paracou research station experiences an average annual rainfall of 3,041 mm and a mean air temperature of 25.7°C (Aguilos *et al*. 2018).

## B - Sampling

We surveyed Sixto on the 13 October 2020 by climbing the tree and selecting suitable boughs that had branches with contrasting light exposure, and then proceeded to sample: 1) Cambium: three cambium tissue samples were collected from the base of the trunk, approximately 1.3m above ground and equidistantly across the perimeter of the tree trunk; 2) Leaves: three leaves from the same twig per branch from eight branches were sampled, the branches were selected in three pairs each in a different bough, plus two independent branches in two other boughs. Branches on boughs were chosen so that each pair of branches contained one light-exposed and one shaded branch, the shade being mostly a result of self-shading (see section C).; and 3) Fruits: fruits were collected from 5 different branches where leaves were collected (Fig. B3). All boughs and branch lengths, and basal and terminal diameters were measured for branches with basal diameters above 10 cm, in addition to the trunk. Sixto was felled on the 14th of October 2020 for analysis of chemical and microbiome heterogeneity across the whole tree for another project, thus the tree is not available for future studies. The sampling of Angela was similar: On 22 April 2021, we collected: 1) Cambium: three cambium tissue samples were sampled from the base of the trunk of Angela, approximately 1.3m above ground and equidistantly across the perimeter of the tree trunk; 2) Leaves: three leaves from the same twig per branch from ten branches were sampled, the branches were selected as five pairs of branches each in a different bough. Branches on boughs were chosen so that each pair of branches contained one light-exposed and one shaded branch, the shade being mostly a result of self-shading; and, 3) fruits from 4 different branches where leaves were collected (Fig. B4). On 15, 16 and 22 November 2021, we described the architecture of Angela: all branch lengths, and basal and terminal diameters were measured for branches with basal diameters above 10 cm, in addition to the trunk. On 21, 22, and 23 March 2022, 30 wood cores were sampled using a borer across the crown and in the trunk of Angela. Angela was not felled, and is not among the trees planned for felling by the ONF, so it will be accessible for future studies. In both species, leaves,cambium, and fruit samples were flash-frozen in liquid nitrogen and stored at −80°C until DNA and RNA extraction.

## C - Characterisation of light conditions

A linear PAR ceptometer (AccuPar, Decagon Devices, Pullman, WA, USA) was used on each sampling position on both trees during sampling to measure direct incident light around midday in comparison to open incident light measured on the closest road (Tab C1). In addition, for Angela, a terrestrial and drone lidar campaign (TLS, Faro Focus3D 120; DLS, Yellowscan Vx20-100) was carried out on 3 May 2021 to map the canopy transmittance in order to derive an estimate of the irradiance in different locations in the canopy. TLS scans were done horizontally from 0-360° and vertically from −60-90° resulting in 174.8 millions of points per scan for 10 scans in a forest gap close to the tree and 4 scans from the nearby road. DLS scans were taken at 35 m above the focal tree in 2 perpendicular flights with flight lines 10 m apart in a circular area 150 m in diameter above the focal tree, resulting in 46.6 million points. Before lidar acquisition, reflective strips were put on sampling points by tree climbers to detect sampling points in the lidar cloud (Fig. C1-A). The AMAPvox software (Vincent *et al*. 2017) was used to compute an annual illumination index from the lidar point cloud. The plant area density (PAD, m2/m3) was calculated for the focal tree in context (with a diameter of 30m surrounding the tree) using 1 m3 voxels. An estimate of the yearly proportion of the above canopy solar radiation received at the sampling point was simulated considering a clearness index of 0.5 and a latitude of 5 degrees. Transmittance uncertainty due to sampling point locations uncertainties was further evaluated by randomly sampling 10 positions around sampling points up to 0.5 metres (Fig. C1-B) and revealed low variations in transmittance (Table C2). The estimates were in agreement with the light/shade classification of branches identified by the tree climbers (Table C2).

## D - Genome assemblies and annotations

The genomes used in the project were extracted from the same individuals as the ones used to detect mutations, ensuring the best possible correspondence between the reference genomes and the resequencing libraries. High molecular weight (HMW) DNA was extracted from 2g of frozen leaves using QIAGEN Genomic-tips 500/G kit (Qiagen, MD, USA). We followed the tissue extraction protocol. Briefly, 2g of young leaf material was ground in liquid nitrogen with mortar and pestle. After 3h of lysis and one centrifugation step, the DNA was immobilised on the column. After several washing steps, DNA was eluted from the column, then desalted and concentrated by Isopropyl alcohol precipitation. A final wash was done with 70% ethanol before resuspending the DNA in the EB buffer. DNA quantity and quality were assessed with NanoDrop and Qubit (Thermo Fisher Scientific, MA, USA). DNA integrity was also assessed using the Agilent FP-1002 Genomic DNA 165 kb on the Femto Pulse system (Agilent, CA, USA). Hifi libraries were constructed using SMRTbell® Template Prep kit 2.0 (Pacific Biosciences, Menlo Park, CA, USA) according to PacBio recommendations (SMRTbell® express template prep kit 2.0 - PN: 100-938-900). HMW DNA samples were first purified with 1X Agencourt AMPure XP beads (Beckman Coulter, Inc, CA USA), and sheared with Megaruptor 3 (Diagenode, Liège, BELGIUM) at an average size of 20 kb. After end repair, A-tailing and ligation of SMRTbell adapter, the library was size-selected on a BluePippin System (Sage Science, MA,USA) using a size range of 10-50kb. The size and concentration of libraries were assessed using the Agilent FP-1002 Genomic DNA 165 kb on the Femto Pulse system and the Qubit dsDNA HS reagents Assay kit. Sequencing primer v5 and Sequel® II DNA Polymerase 2.2 were annealed and bound, respectively, to the SMRTbell libraries. Each library was loaded on 2 SMRTcell 8M at an on-plate concentration of 90pM. Sequencing was performed on the Sequel® II system at Gentyane Genomic Platform (INRAE Clermont-Ferrand, France) with Sequel® II Sequencing kit 3.0, a run movie time of 30 hours with an Adaptive Loading target (P1 + P2) at 0.75. We obtained 1,898,004 corrected reads for *D. guianensis* (N50=21,233; DP=58.7X), that we assembled in 562 contigs (N50=37.76Mb; L50=8 contigs; GC=37.25%) using *HiFiasm assembler* (v0.15.5, Cheng *et al*., 2021). *Hifiasm* is able to produce 2 haplotype-aware assemblies where the alleles are separated in two files. All the metrics reported correspond to the haplotype1 file and only this one was used for follow-up analyses. Similarly, we obtained 6,624,997 corrected reads for *S. rubra* (N50=17,577; DP=114X), that we assembled in 747 contigs (N50=16.513Mb; L50=17 contigs; GC=38.50%).

HMW DNA was additionally used to produce optical maps for hybrid scaffolding. The HMW DNA was labelled and stained according to the Direct Label and Stain (DLS) protocol (BNG). Briefly, labelling was achieved by incubating 750 ng genomic DNA with 1× DLE-1 Enzyme (BNG) for 2 hours in the presence of 1× DL-Green (BNG) and 1× DLE-1 Buffer (BNG). Following proteinase K digestion and DL-Green clean-up, the DNA backbone was stained by mixing the labelled DNA with DNA Stain solution (BNG) in the presence of 1×Flow Buffer (BNG) and 1× DTT (BNG), and incubating overnight at room temperature. The DLS DNA concentration was measured with the Qubit dsDNA HS Assay (Invitrogen, Carlsbad, CA, USA). Labelled and stained DNA was loaded on 1 Saphyr chip for each species. The chips were loaded and run on the BNG Saphyr System according to the Saphyr System User Guide. Digitalised labelled DNA molecules were assembled to optical maps using the BNG Access software. A total of 972 Gb (*D. guianensis*) and 720 Gb (*S. rubra*) of molecules with a size larger than 150kb, the threshold for map assembly, were generated representing 1,500x and 720X of genome coverage, respectively. The *D. guianensis* assembly produced 511 genome maps with a N50 of 15.7 Mb for a total genome map length of 903 Mbp. The *S. rubra* assembly produced 314 genome maps with a N50 of 14,95 Mb for a total genome map length of 1329 Mbp. Finally, a hybrid scaffolding was assembled between the sequence assembly and the optical genome maps with hybridScaffold pipeline (https://bionano.com/wp-content/uploads/2023/01/30073-Bionano-Solve-Theory-of-Operation-Hybrid-Scaffold.pdf). For *D. guianensis*, we obtained 54 hybrid scaffolds (N50=38.450Mb; N=0.76%; 571 gaps), while 515 contigs remained unanchored with a total length of 28,784Mb representing 4.97% of the genome. For *S. rubra*, we obtained 35 hybrid scaffolds (N50=60.458Mb; N=1%; 1,923 gaps), while 609 contigs remained unanchored with a total length of 53,067Mb representing 5.08% of the genome. Genome quality was assessed with BUSCO (Seppey *et al.,* 2019) and Merqury (Rhie *et al.,* 2020). Specifically, alongside the contiguity and error rates, we focused on the following criteria: <5% false duplications, >90% k-mer completeness, >90% sequences assigned to candidate chromosome, >90% single copy conserved genes complete and single, and >90% transcripts from the same organism mappable. In case of issue with duplication we further looked into Merqury copy number spectrum plots.

We built an automated workflow for genome annotation named *genomeAnnotation* (see script availability). We used both *singularity* containers (Kurtzer *et al.,* 2017) and the *snakemake* workflow engines (Köster *et al.,* 2012) to make the workflow highly reproducible (FAIR), and scalable. The workflow accomplishes: (i) *de novo* and known transposable element (TE) detection, (ii) *de novo* and known gene models detection, and (ii) gene functional annotation. *De novo* TE detection uses *RepeatModeler2* (v2.0.3; Flynn *et al*., 2020) followed by classification using *RepeatClassifier* (v2.0.3; Flynn *et al*., 2020) and *TEclass* (v2.1.3; Abrusán *et al*., 2009). The obtained *de novo* TE are merged with known TE accessions for *Viridiplantae* from *RepBase* (v27.07; Kapitonov and Jurka 2008). This consolidated database is used for TE detection in each genome before soft masking using *RepeatMasker* (v2.0.3; Flynn *et al*., 2020). *De novo* and known gene models detection rely on *BRAKER2* and its dependencies (Brůna *et al*., 2020). Finally, functional annotation of candidate genes is based on the *Trinotate* pipeline (v3.2.1; Bryant *et al*., 2017) using *TransDecoder* (v5.5.0; https://github.com/TransDecoder/TransDecoder), *TMHMM* (Krogh *et al*., 2001), *HMMER* (Finn *et al*., 2011), *BLAST* (v2.13.0; Altschul *et al*., 1990), *RNAmmer* (v1.2; Lagesen *et al*., 2007), and *SignalP* (v4.1; Petersen *et al*., 2011) with *UniProt* (Bateman *et al*., 2015) and *Pfam* databases (Punta *et al*., 2012).

Two high quality genomes were obtained for *Dicorynia guianensis* and *Sextonia rubra* with a total length of 550 and 991 Mb and 54 and 35 super-scaffolds anchored with optical maps, respectively, including 20 super-scaffolds > 1 Mb in each species. Evaluation of completeness for the two assemblies was done with BUSCO (Seppey *et al.,* 2019), revealing near complete assemblies (>99% of the *Viridiplantae* housekeeping genes, 352 and 409 complete single-copy genes for *Dicorynia guianensis* and *Sextonia rubra,* respectively, Fig. D1). Assessment of genome quality with Merqury (Table D1, Fig. D8-9, Rhie *et al.,* 2020) revealed only good scores according to the defined criteria (Table D1), with the exception of duplication for *D. guianensis*, which was higher (14%) than the 5% criterion. A closer examination of the Merqury copy number spectrum plots for haploid and diploid assemblages in *D. guianensis* (Fig. D8b) shows that alleles are well separated and distributed in both haplotypes (similar black and red curves), indicating that the 14% duplication is due to true duplication. Genome heterozygosity (π) was estimated based on K-mer distributions and was found to be 0.9% and 0.7% for *Dicorynia guianensis* and *Sextonia rubra,* respectively (Fig. D2 & D3). Consistent with the smaller genome size of *D. guianensis*, we detected fewer genes (15,490) and a lower transposable elements content (50.8%) in *D. guianensis* as compared to *S. rubra* (21,412 and 63.8%, respectively). Genomic distributions of TE were highly heterogeneous across super-scaffolds (Fig. D4 & Fig. D5). TE annotations revealed a majority of long terminal repeat elements (16%-23.3%) and DNA transposons (3.4%-16%, Fig. D6 & Fig. D7).

## E - Leaf and cambium mutation detection

Genomic DNA was extracted from 30 mg of frozen leaf or cambium tissue per sampling point for both trees with a CTAB protocol with chloroform - isoamyl alcohol (24:1) extraction, isopropanol precipitation and pellet resuspension in 1x Low TE (10 mM Tris-HCl + 0.1 mM EDTA; Doyle and Doyle 1987). DNA was quantified using a Qubit HS assay (Thermo Fisher Scientific, Waltham, MA, USA) and purified with AMPure XP beads (Beckman Coulter Genomics, Danvers, MA, USA) when required to allow library preparation. An Illumina sequencing library was produced for each leaf using an optimised NEBNext Ultra II DNA library protocol (New England Biolabs, Ipswich, MA, USA). Libraries were pooled in multiplexes after tagging each library independently before whole genome sequencing (WGS) on a S4 flow cell and in a NovaSeq 6000 instrument with v1.5 chemistry (2 × 150 PE mode). We obtained 33 cambium and leaf libraries for Angela with a sequencing depth of about 160X and 27 libraries with a depth of about 80X for Sixto (Fig. E1).

We built a workflow to detect somatic mutations from mapped sequencing reads on a genome reference named *detectMutations* (see script availability and Schmitt *et al*., 2022). A simulation approach has already been used to verify the somatic mutation detection workflow (Schmitt *et al.,* 2022) by generating Illumina-like sequence reads spiked with simulated mutations at different allelic fractions to compare the performance of seven commonly-used variant callers to recall them. S*ingularity* containers (Kurtzer *et al.,* 2017) and the *snakemake* workflow engines (Köster *et al.,* 2012) were used to ensure an automated, highly reproducible (FAIR), and scalable workflow. Paired-end sequencing reads of every library are quality checked using *FastQC* (v0.11.9) before trimming using *Trimmomatic* (v0.39, Bolger *et al.,* 2014) keeping only paired-end reads without adaptors and a phred score above 15 in a sliding window of 4 bases. Reads are aligned against the reference genome using BWA *mem* with the option to mark shorter splits (v0.7.17, Li & Durbin, 2009). Alignments are then compressed using *Samtools view* in CRAM format, sorted by coordinates using *Samtools sort,* and indexed using *Samtools index* (v1.10, Li *et al.,* 2009). Duplicated reads in alignments are marked using *GATK MarkDuplicates* (v4.2.6.1, Auwera *et al.,* 2013). Sequencing depth is estimated along the genome using *Mosdepth* (v0.2.4, Pedersen *et al.,* 2018) globally and on a 1-kb sliding window.

We used both a K-mer based and an alignment based method to estimate heterozygosity and detect heterozygous sites. We used *Jellyfish* (v1.1.12; Marçais and Kingsford 2011) and *GenomeScope* (Vurture *et al.,* 2017) to estimate heterozygosity up to 21-mer. We used *GATK (HaplotypeCaller, GatherGVCFs, GenomicsDBImport, GenotypeGVCFs;* Auwera *et al.,* 2013) to call heterozygous sites from previously obtained alignments. We filtered single nucleotide polymorphisms (SNP) using *bcftools* (v1.10.2, Danecek *et al.,* 2021), *GATK VariantFiltration* (v4.2.6.1, Auwera *et al.,* 2013), and *plink* (v1.90, Chen *et al.,* 2019). To filter candidate variants, we kept only biallelic SNPs with a quality above 30, a quality by depth above 2, a Fisher strand ratio below 60 and a strand odds ratio below 3. To remove all sites that were truly heterozygous, we further filtered SNPs present in all genotypes and sampled tissues (no missing data) and shared by at least all tissues but one. Finally, the workflow uses *Strelka2* (v2.9.10, Kim *et al.,* 2018) to detect mutations, a variant caller developed initially for cancer research. This software has been shown to perform best in our study design (Schmitt *et al*., 2022). *Strelka2* identifies mutations by comparing two samples, one mutated and one normal sample (directional). To detect cambium mutations present at the base of the tree we compared all potential pairs (6 in total) among the three cambium libraries. To detect leaf mutations we called each leaf library against the first cambium library (T2 for Anglea and T1 for Sixto) as the normal sample. We filtered from leaf candidate mutations previously identified heterozygous sites and all candidate mutations from all the cambium comparisons (Fig. E2) using *BEDTools subtract* (v2.29.2, Quinlan & Hall, 2010). We further filtered mutations using the following criteria: (i) no copy of the mutated allele in the reference sample being here the cambium sample; (ii) a read depth for the two samples between the 5^th^ quantile and the 95^th^ quantile of the coverage of the corresponding library; and (iii) the presence of the mutation in at least two biological replicates (at least 2 leaves from the crown). We produced two datasets with the filtered mutations: (i) all filtered mutations; and (ii) a more stringent dataset that passed the empirical variant score (EVS) filtering of *Strelka2* (v2.9.10, Kim *et al.,* 2018). The different steps are summarised in Fig. E3. The results with the stringent dataset based on the empirical variant score (EVS) are shown in supplementary note K. We used the same pipeline and compared detected mutations on two pedunculate oaks *Quercus robur* (Schmitt *et al*., 2022) named Napoleon (Schmid-Siegert *et al*. 2017) and 3P (Plomion *et al*. 2018), and on a dataset from one tortuous phenotype of common beech *Fagus sylvatica* (Aury Plomion 2023ab). We detected a total of 15,066 unique somatic mutations in Angela and 3,208 in Sixto by comparison with the cambium reference samples. Similarly, we found 2,356 and 13,976 unique somatic mutations in Napoleon and 3P, respectively, and 6,560 unique somatic mutations in the tortuous beech.

## F - Mutations along the tree architecture

We explored mutation distribution along tree architecture by assuming the origin of the mutation in the tree architecture was at the latest the most recent common branching event among all branches harbouring the mutation (Duan *et al.,* 2022). Among the 15,066 and 3,208 unique somatic mutations identified in Angela and in Sixto, respectively, 10,849 (72.0%) and 1,381 (43.0%) of the mutations originated at the latest from the base of the crown, a proportion far more important than from the tips (824 (5.5%) and 283 (8.8%) respectively; Fig. F1). Nevertheless most mutations were only weakly shared among sampling points, *i.e.* only by a few branches among all branches in common (Fig. F2 & Fig. F3), resulting in low correlations among sampling points (Tab. F1 & Tab. F2). We further built mutation phylogenies using *iqtree* rooting the tree with the non-mutated library from the cambium mean genotype without mutations (Nguyen *et al.,* 2015), and found very divergent sampling points in both individuals with low support for the tree topology (Fig. F4). We compared phylogenies to the physical architecture of both trees with the *dendextend* R package (Galili 2015). Contrary to the general expectation in plants, the observed phylogeny at the somatic mutation positions do not follow the tree architecture, neither in Angela nor Sixto (Fig. F5 and Fig. F6). For stringency, we further constructed the phylogeny with mutations shared by all 3 leaves of each branch, and found no more consistency with the tree architecture (Fig. F7). Finally, we reuse the code of Orr et al., (2020) using the *rmtopology* and *dist.topo* functions from *ape* R package to compare the path distance of the inferred phylogenies from the physical trees compared to all possible physical trees typologies with the same number of tips. The phylogeny observed was not significantly closer to the physical tree topology than expected by chance (Fig. F8).

## G - Light and somatic mutations

We explored the effects of light on the occurrence of mutations in the trees using Student’s T-tests and Kolmogorov-Smirnov tests. We compared the number of mutations detected in branches exposed to high vs. low light conditions using the leaves as an observation. Student’s T-tests revealed no differences in the number of accumulated mutations between light and shadow conditions in both trees (Fig. G1), while Kolmogorov-Smirnov tests revealed non-identical distributions between the two conditions (Fig. G2). We further compared mutation types (base change, Fig. G3) and mutation spectra (mutation context with 5’ and 3’ bases, Fig. G4 & Fig. G5) between high and low light conditions among branches of each tree. Student’s T-tests again revealed no differences of mutation types and spectra between light and shadow conditions (Fig. G3, Fig. G4 & Fig. G5). We further used Bayesian methods to address small sample problems. Bayesian methods do not rely on asymptotics, a property that can be a hindrance when employing frequentist methods in small sample contexts. Joint distributions of the mutation spectrum were inferred within each species separately. We used a softmax regression within a conjugated Dirichlet Process and Multinomial distribution to infer the distribution of spectrum jointly and not independently:

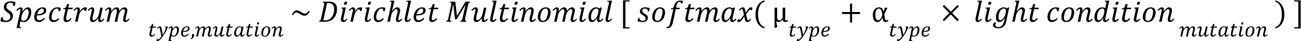

Where *Spectrum _type,mutation_* represents the spectrum for every mutation as a simplex of 0 s and 1 s (i.e. 0 for all other types and 1 for the considered spectrum type), *light condition _mutation_* is the light condition (shadow or light) of the branch where the mutation is observed, μ*_type_* is the vector of mutations mean frequencies (intercept), and α*_type_* is the vector of light effect on spectrum frequency. Since each distribution (one per triplet of the spectrum) intercepts the zero value (Fig. G6), our result is consistent with no significant effect of light on the spectrum frequencies for both species.

## H - Low-frequency mutations

We explored the allelic fractions of somatic mutations in relation to tree sequencing depth (Fig. E1), a known determinant of the sensitivity of somatic mutation detection (Schmitt *et al*. 2022), for Sixto; Angela; two pedunculate oaks, *Quercus robur*, named Napoleon (Schmid-Siegert *et al*. 2017) and 3P (Plomion *et al*. 2018); and a dataset from one tortuous phenotype of common beech *Fagus sylvatica* named Verzy (Aury Plomion, 2023). Most detected mutations were at low frequency (Fig. H1). The discrepancy in the number of observed mutations among sampled trees can be explained by an enrichment in low-fraction mutations in Angela (Fig. H1) detected thanks to the deeper sequencing (Fig. E1). For an allelic fraction above 0.25 (*i.e.,* medium-frequency to fixed mutations), we found the two tropical trees, Angela and Sixto, to have the lowest rate of somatic mutations, with 3 and 6 somatic mutations on 10 and 8 branches for Angela and Sixto, respectively, versus 56 to 421 somatic mutations on 2 to 3 branches for oak and beech (Fig. H1). However, as the number of individuals is small and differences can be linked to species identity with species from very distant lineages, tree ages (with unknown ages for tropical trees), and other life history traits, we did not feel comfortable interpreting these differences.

We further compared mutation annotations in terms of their presence in transposable elements (TE) and genes among trees. We assessed mutation functional impact using *SNPeff* (Cingolani *et al.,* 2012) and related non-synonymous mutations to their functional annotations, gene ontology, and allelic fraction. Focusing on coding regions, we detected 314 and 9 non-synonymous mutations and 567 and 31 synonymous mutations in Angela and in Sixto, respectively (Fig. H2). Gene ontology enrichment of genes associated to non-synonymous mutations was observed at genes coding for viral processes and in biotic interactions with other organisms in Angela (Fig. H3). Unfortunately, the 9 non-synonymous mutations in Sixto were not sufficient for similar analyses. We finally explored the allelic fraction of mutations depending on synonymy and found a significant negative effect (p=1.47*10^-14^) of non-synonymy on the allelic fraction of mutations in both Angela and Sixto, indicating that novel non-synonymous mutations increase in allele frequency less frequently than synonymous ones, which suggests a potential overall negative intra-individual selection (Fig. H4).

## I - Fruit mutations redetections

We explored mutation transmission to fruit using amplicon resequencing. We kept as candidate mutations for redetection only mutations present in at least three leaves from the branches that had fruits during sampling for resequencing, which resulted in 160 candidate mutations (124 for Angela and 36 for Sixto). Frozen fruits were dissected in 4 tissues: (i) embryo sac, (ii) nucellus, (iii) pericarp, and (iv) fruit base. Genomic DNA was extracted from 10-50 mg of frozen fruit tissue for both trees and additional leaf tissue for positive control with a CTAB protocol with chloroform - isoamyl alcohol (24:1) extraction, isopropanol precipitation and pellet resuspension in 1x Low TE (10 mM Tris-HCl + 0.1 mM EDTA; Doyle and Doyle 1987). DNA was quantified using a Qubit HS assay. Primer3plus (Untergasser et al., 2012) was used to design primer pairs targeting candidate mutations (amplicon size between 100 and 200 pb). Only one Angela candidate mutation failed to yield a primer pair. Illumina universal tags 5′-TCGTCGGCAGCGTCAGATGTGTATAAGAGACAG-3′ and 5′-GTCTCGTGGGCTCGGAGATGTGTATAAGAGACAG-3′ were added to the 5′ end of the forward and reverse primer sequences respectively. Oligonucleotides were ordered in a plate format from Integrated DNA Technologies with standard desalt purification at 25 nmoles synthesis scale. Each primer pair was tested using simplex PCR amplification of one DNA sample per species in a volume of 10 μL containing 2 μL of 5X Hot Firepol Blend master mix (Solis Biodyne), 1 µL of 2µM primer pairs, 1 µL of DNA (10 ng/µL), and 6 µL of PCR-grade water. We amplified the PCR on a Veriti 96-Well thermal cycler (Applied Biosystems) which consisted in an initial denaturation at 95°C for 15 min, followed by 35 cycles of denaturation at 95°C for 20 s, annealing at 59°C for 60 s, extension at 72°C for 30 s, and a final extension step at 72°C for 10 min. We checked the amplification on a 3% agarose gel. A total of six Angela primer pairs that failed to amplify were discarded at this stage. The remaining 101 Angela and 33 Sixto primer pairs, targeting respectively 117 and 36 mutations, were grouped accounting for potential primer dimer formation using Primer Pooler (Brown et al., 2017) for subsequent multiplex PCR amplification. Four multiplexed PCRs were done for each species in a volume of 10µL using 2 µL of 5X Hot Firepol Multiplex master mix (Solis Biodyne), 1 µL of multiplex primer mix (0.5 µM of each primer), 2 µL of DNA (10 ng/µL), and 5 µL of PCR-grade water. The amplification was done a Veriti 96-Well thermal cycler (Applied Biosystems) using an initial denaturation at 95°C for 12 min followed by 35 cycles of denaturation at 95°C for 30 s, annealing at 59°C for 180 s, extension at 72°C for 30 s, and a final extension step at 72°C for 10 min. The amplicons from the four multiplexed PCRs of each sample were pooled. Illumina adapters and sample-specific Nextera XT index pairs were added to the amplicons by a PCR targeting the Illumina universal tags attached to the locus-specific primers. This indexing PCR was done in a volume of 20 μL using 5X Hot Firepol Multiplex master mix (Solis Biodyne), 5 µL of amplicon, and 0.5 µM of each of the forward and reverse adapters, using an initial denaturation at 95°C for 12 min followed by 15 cycles of denaturation at 95°C for 30 s, annealing at 59°C for 90 s, extension at 72°C for 30 s, and a final extension step at 72°C for 10 min. We then pooled the libraries and purified them with 0.9X Agencourt AMPure XP beads (Beckman Coulter, the UK). We checked the library quality on a Tapestation 4200 (Agilent) and quantified it using QIAseq Library Quant Assay kit (Qiagen, Hilden, Germany) in a Roche LightCycler 480 quantitative PCR. The libraries were sequenced on an Miseq sequencer (Illumina, San Diego, CA, USA) using a V2 flow cell with a 2×150 bp paired-end sequencing kit. Paired-end sequencing reads of every library were trimmed using *Trimmomatic* (v0.39, Bolger *et al.,* 2014) keeping only paired-end reads without adaptors and a phred score above 15 in a sliding window of 4 bases. Reads were aligned against the reference genome using *BWA mem* with the option to mark shorter splits (v0.7.17, Li & Durbin, 2009). Alignments were then compressed using *Samtools view* in CRAM format, sorted by coordinates using *Samtools sort,* and indexed using *Samtools index* (v1.10, Li *et al.,* 2009). We used *GATK (HaplotypeCaller, GatherGVCFs, GenomicsDBImport, GenotypeGVCFs;* Auwera *et al.,* 2013) to call candidate mutations transmitted to fruits from previously obtained alignments. We filtered variants that corresponded to the 160 candidate mutations (124 for Angela and 36 for Sixto). The different steps are summarised in Figure E3.

For stringency, we highlighted and discussed only those candidate mutations with at least a positive heterozygous call detected by *GATK GenotypeGVCFs* (red border in Fig. I1). We found 23 candidate mutations present in Angela’s fruit reads and 9 in Sixto’s fruit reads distributed across the different fruits and their respective tissues. For almost all mutations (97% of mutations x library pairs), we observe that the mutations recovered in the fruit have a higher allele fraction than in the plant, as expected for inherited mutations (as visually recovered in Fig. I1). However, the allele frequency is rarely higher than 40%, as we would have expected for a heterozygous site and given the high coverage of these markers (60±42X). There are several possible explanations:

- amplification bias during PCR due to random or amplicons affinity,
- amplification bias during PCR due to multiplexing of amplicons (including amplicons matching the other, non-focal species) in a single PCR,
- lack of knowledge of the origin of fruit tissue in fruit development with regard to which tissues are derived from the mother and which from the embryo,
- tissue contamination between those expected to be derived from mother and those from embryo during dissection due to the small size of the fruits.

Mutation “validation” should remain considered as a challenging task, and we encourage future work based on plant tissues derived from germinated seeds (*e.g.* first leaves), rather than directly dissected seeds, to overcome the two last potential limitations indicated just above.

In plants, the timing of germline segregation remains debated (Lanfer, 2018), our study provided limited additional information to this interesting and fundamental question. However, by recovering somatic mutations inherited in the progeny, our results appear consistent with a late germline segregation scenario in the two species, in line with some other recent investigations in trees (Prunus: Wang et al., 2019; Quercus: Plomion et al., 2018). But our work was not designed to tackle this question in detail and this point would require further investigations.

## J - Empirical Variant Score Filtering

As described in Kim et al., (2018): “Strelka2’s EVS models are pre-trained on data from a variety of sequencing conditions to improve robustness, and produce a single aggregate score which can be used to set application-specific precision levels or prioritise variants for follow-up”. All germline and RNA-seq datasets are from an individual for which a gold standard truth set is available from the Platinum Genomes project (Eberle et al., 2017). Consequently, the EVS was built on human mutation data with impure normal samples and very high sequencing depth with conditions and assumptions very different from the plant dataset with pure normal samples and lower sequencing depth that we used here. Consequently, we thus choose not to use EVS as a filtering criterion but to filter based on several other criteria that EVS is supposed to summarise and an additional biological one:

i. no copies of the mutated allele in the reference sample, in this case the cambium sample;
ii. a read depth for both samples between the 5th quantile and the 95th quantile of the corresponding library coverage; and (iii) the presence of the mutation in at least two biological replicates (at least 2 leaves of the crown). Moreover, looking at Schmitt *et al.,* (2022) reanalysis revealed that among the 12 somatic mutations detected by Schmid-Siegert *et al.,* (2017) validated by Sanger resequencing, 3 would have been rejected based on the EVS scores. Consequently, we question the use of EVS for plant applications. Nevertheless, to test the robustness of our results, we reproduced our analysis with mutations passing EVS filtering only, and found that:

- Phylogeny does not match the tree architecture (congruent);
- Student’s T-tests revealed no differences of mutation types and spectra between light and shadow conditions (congruent, Fig. K1a-b) except for C>T mutation (incongruent) but in the opposite direction from what we could have expected assuming UV as the mutagen driving this pattern;
- Low-frequency mutations account for the vast majority of within-individual somatic diversity in plants (congruent, Fig. K2);
- Low-frequency mutations are transmitted to the next generation (congruent, Fig. K2a-b);
- We could not conclude that the average allelic fraction at non-synonymous sites was lower than those at synonymous sites (inconclusive, Fig. K2c).

Consequently, our main conclusions, namely the abundance of low frequency mutations, their transmission, the absence of a pattern matching the tree architecture and an absence of a UV-associated spectra would be robust to the introduction of a EVS criterion. In addition, somatic mutations in leaves that are also observed in fruit had significantly higher EVS scores than those not observed in fruit (9.3 vs. 2.1, p < 2e-16), arguing for their robustness and heritability despite their low frequency.

## K - Somatic mutations and population diversity

Any new mutation in the population appears at half the effective population size (*N_e_*), regardless of whether the mutation is of germline or somatic origin, or whether it was initially at a low or high frequency. If the mutation is transmitted, the evolution of the allele frequency will depend on the evolutionary forces, with a particular importance on genetic drift for selectively neutral mutations. Consequently, the identification of new mutations requires the sequencing of families rather than populations, as it would take thousands or tens of thousands of generations for the mutation to increase in allele frequency in the population. Future studies could combine sequencing of families (e.g. trios) with the germline mutation rates in order to know which proportion of the mutation rates has a somatic origin.

---

**Figure B1:**
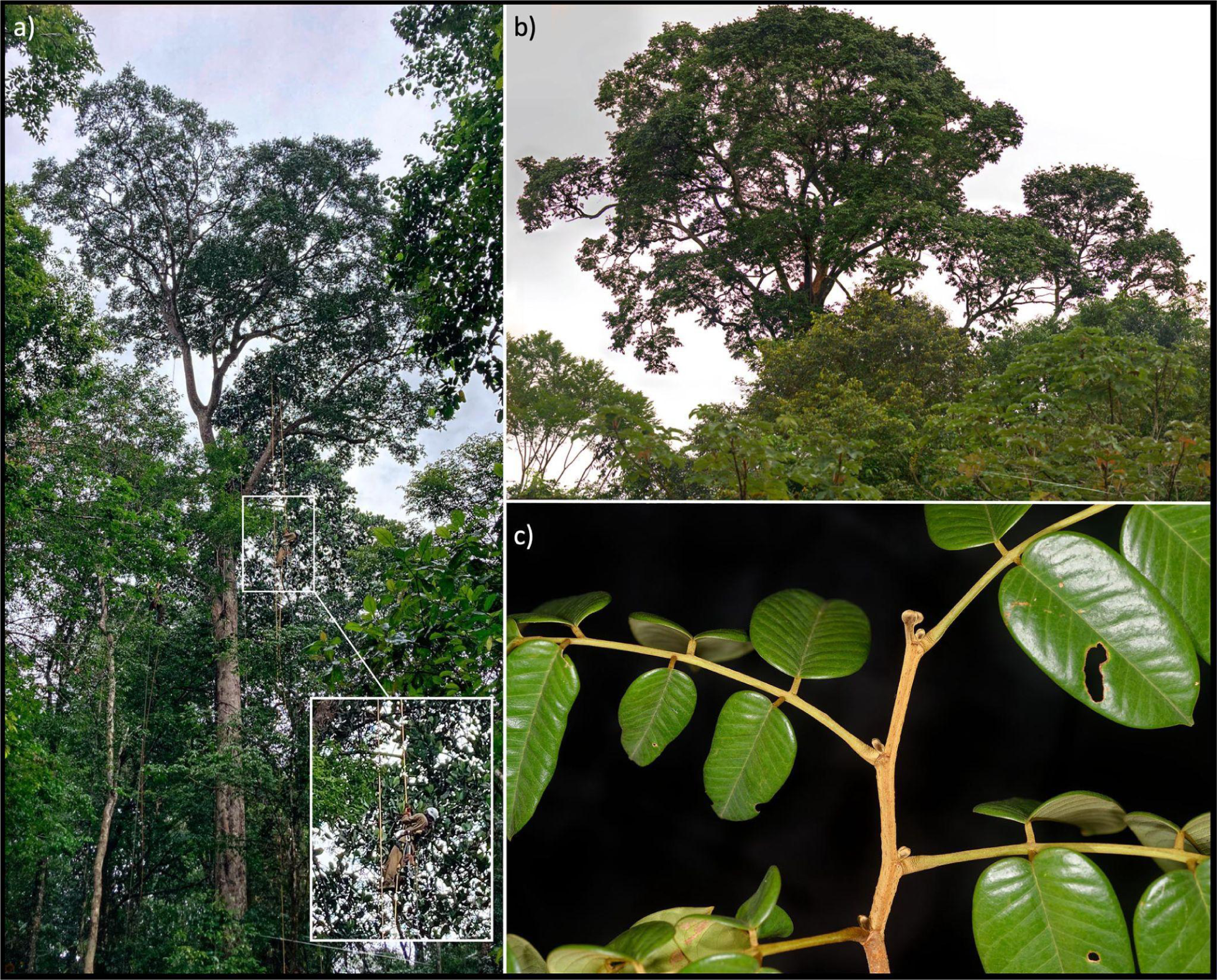
Angela: the Dicorynia guianensis tree sampled in the Régina Saint-Georges forest (4°01’N, 51°59’W). **a)** Angela from the forest gap with an insert showing a tree climber, note that the branch B is hidden by the trunk. **b)** Angela’s crown from the road with the off-centre branch B on the right. **c)** Detail of branch morphology showing compound leaves, alternate phyllotaxy and buds formed by stipules.

**Figure B2:**
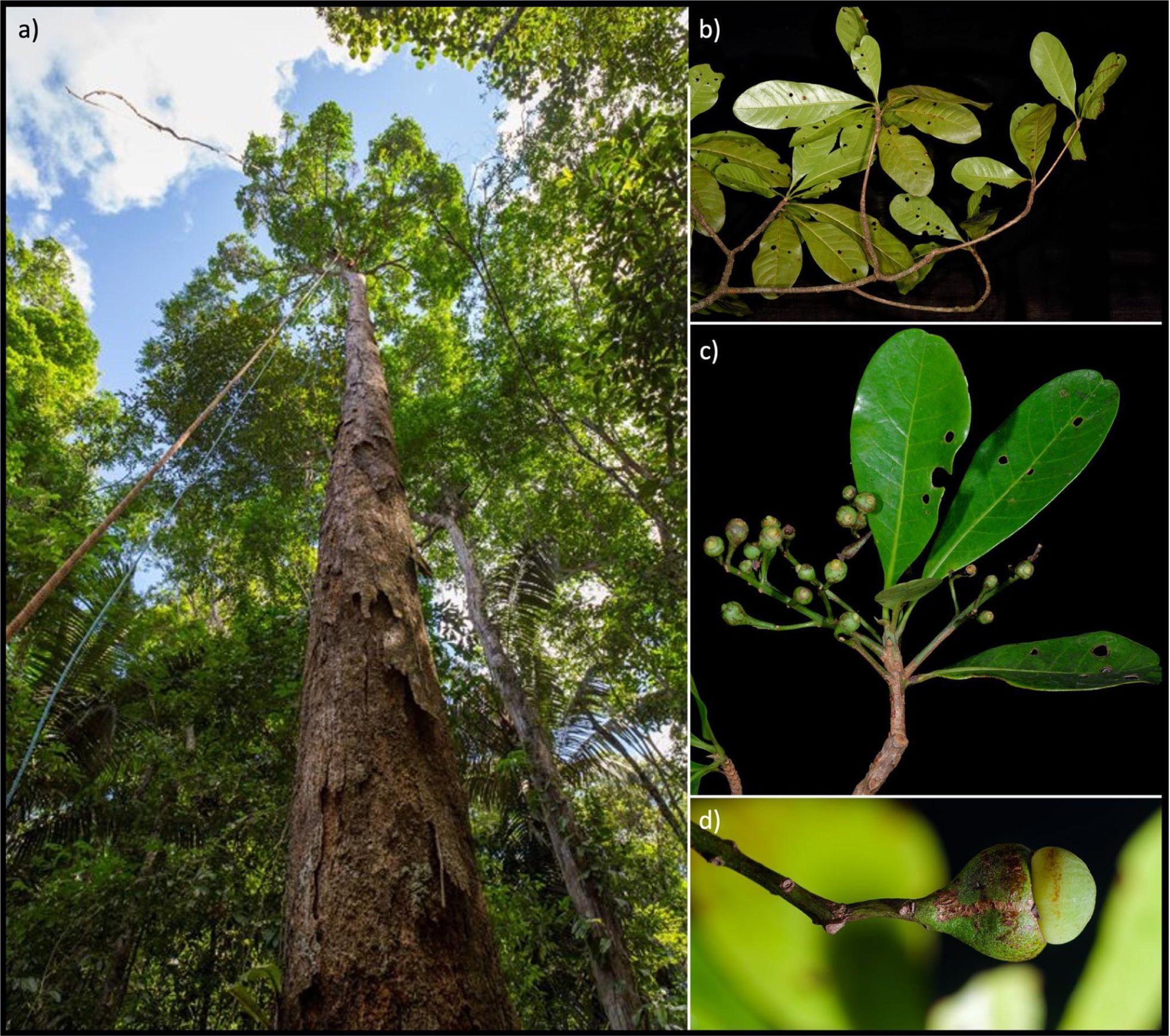
Sixto: the Sextonia rubra tree sampled near the Paracou research station *(5°18’N, 52°53’W)*. **a)** Sixto from the ground. **b-c)** Detail of branch morphology showing simple leaves and alternate phyllotaxy (a, b) and fruit morphology (b,c).

**Figure B3:**
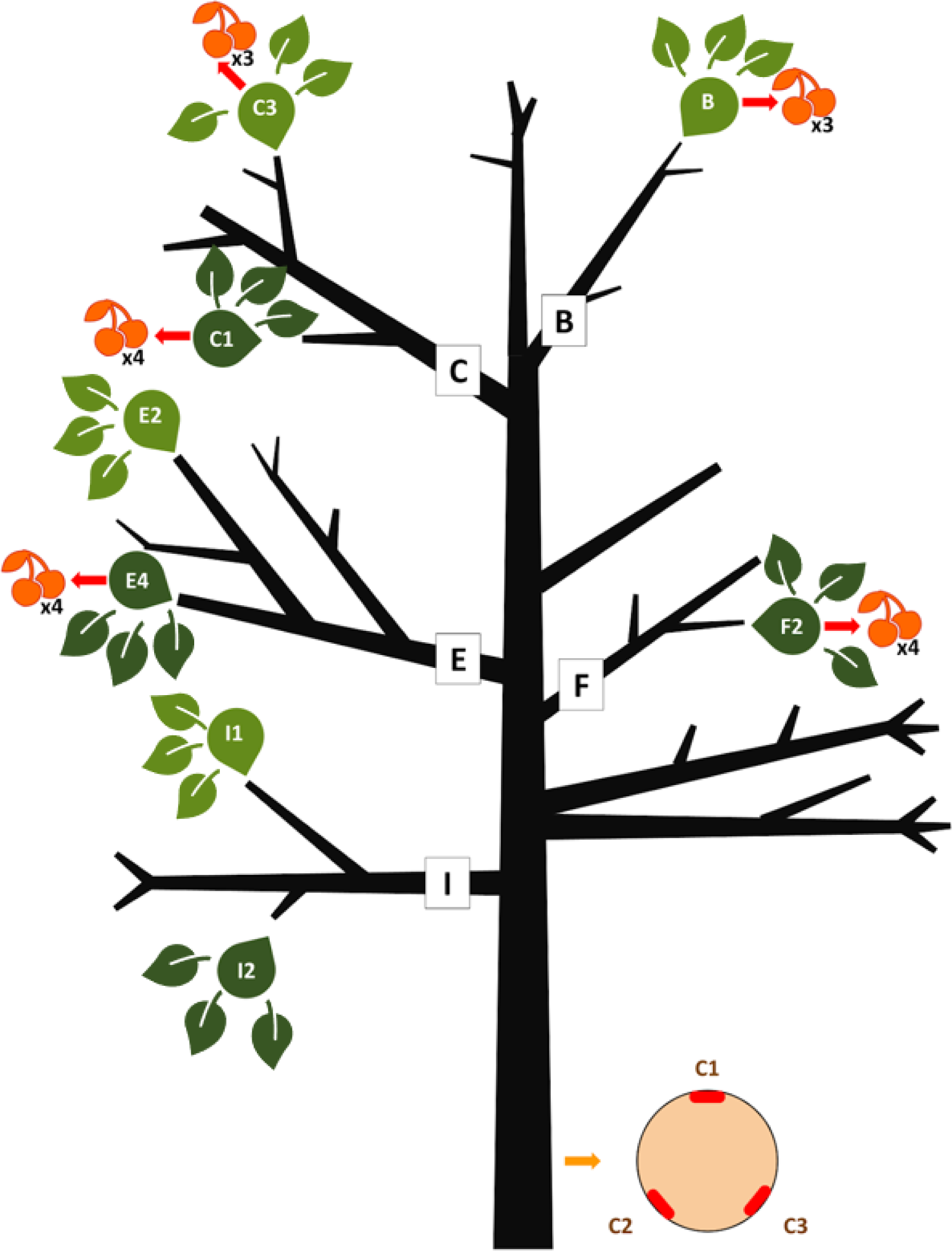
Sixto sampling scheme: Boughs are labelled along the trunk from B to I. The sampling included (i) three cambium samples from the trunk taken at 1m30 height shown with the salmon-coloured circle facing respectively east, north, and southwest, (ii) three leaves per light condition (light green for light-exposed, dark green for shaded branches) in each pair of branches, and (iii) fruits close to leaf samples B, C1, C3, E4, and F2.

**Figure B4:**
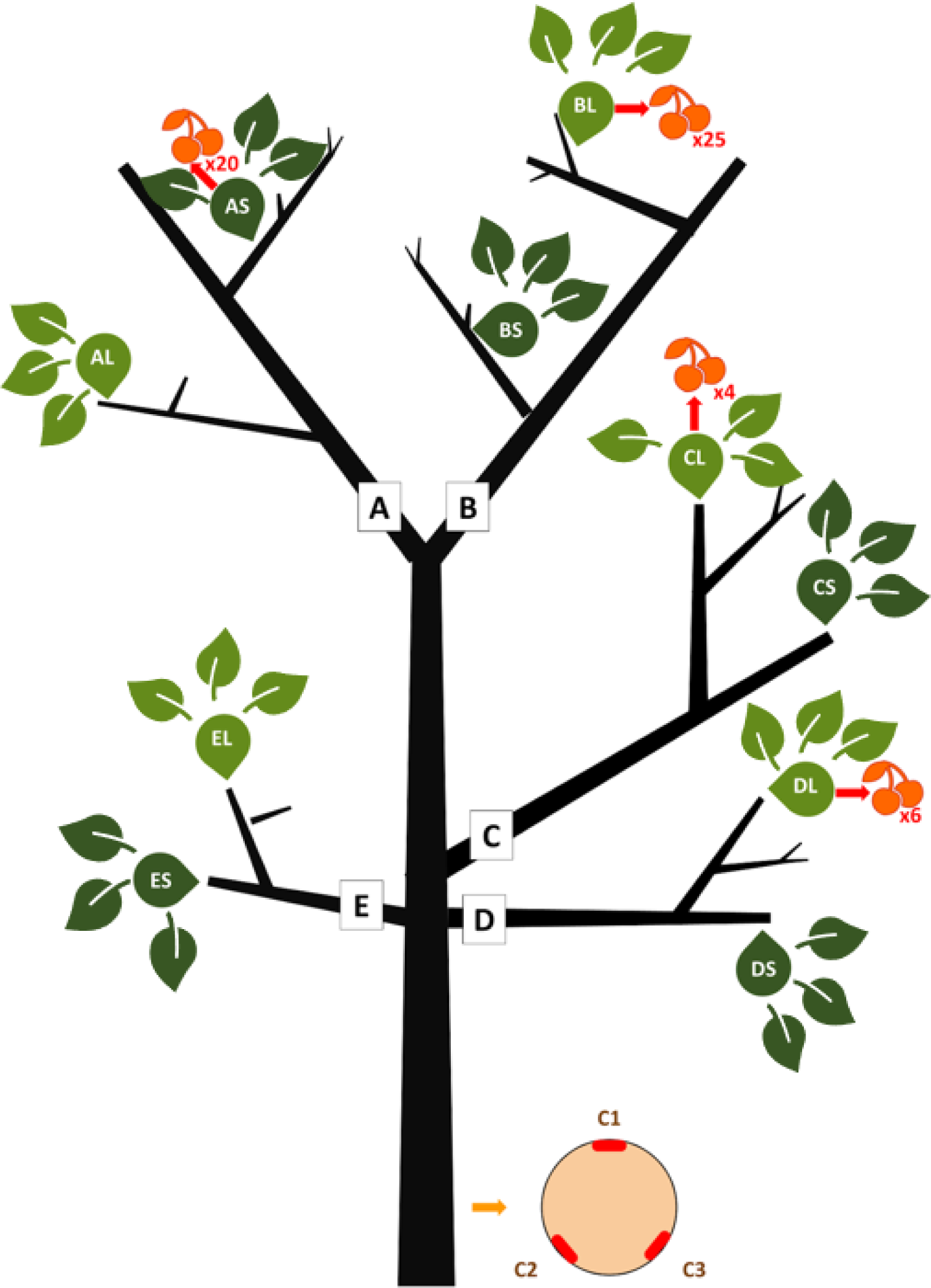
Angela sampling scheme. Boughs are labelled along the trunk from A to E. The sampling included (i) three cambium samples from the trunk taken at 1m30 height shown with the salmon-coloured circle facing respectively east, north, and southwest, (ii) three leaves per light condition (light green for light-exposed, dark green for shaded branches) in each pair of branches, and (iii) fruits close to leaf samples AS, BL, CL, and DL.

**Figure C1:**
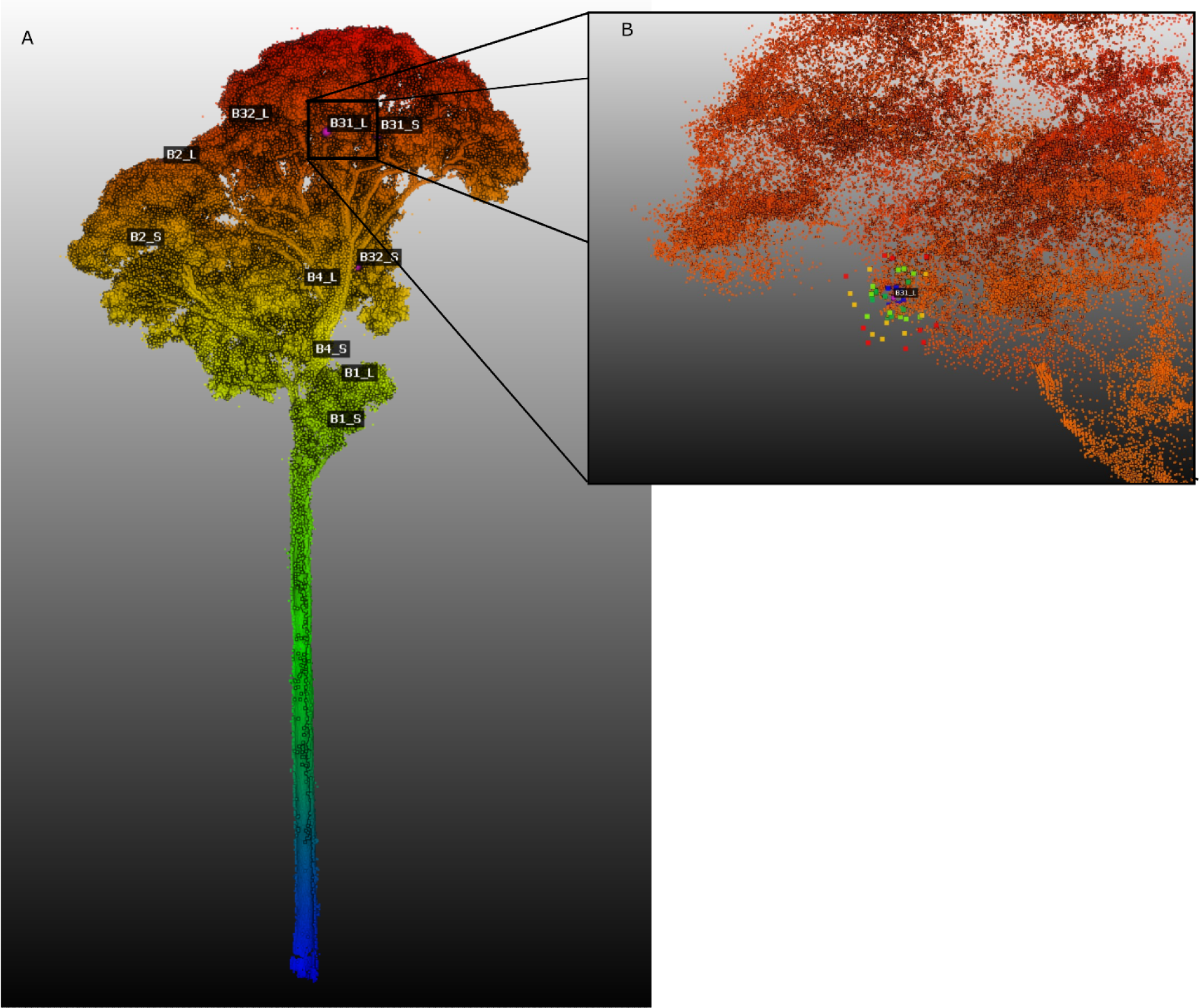
*Dicorynia guianensis* lidar point cloud with sampling points. Lidar point cloud was obtained with terrestrial and drone lidar done on the 3rd of May 2021 to characterise annual transmittance. A: the sampling point; B: The random positions from 0.1m (blue) to 0.5m (red) every 10cm around the B31 Light sampling point. Refer to table 1 for the correspondence between lidar and genomic IDs.

**Figure D1:**
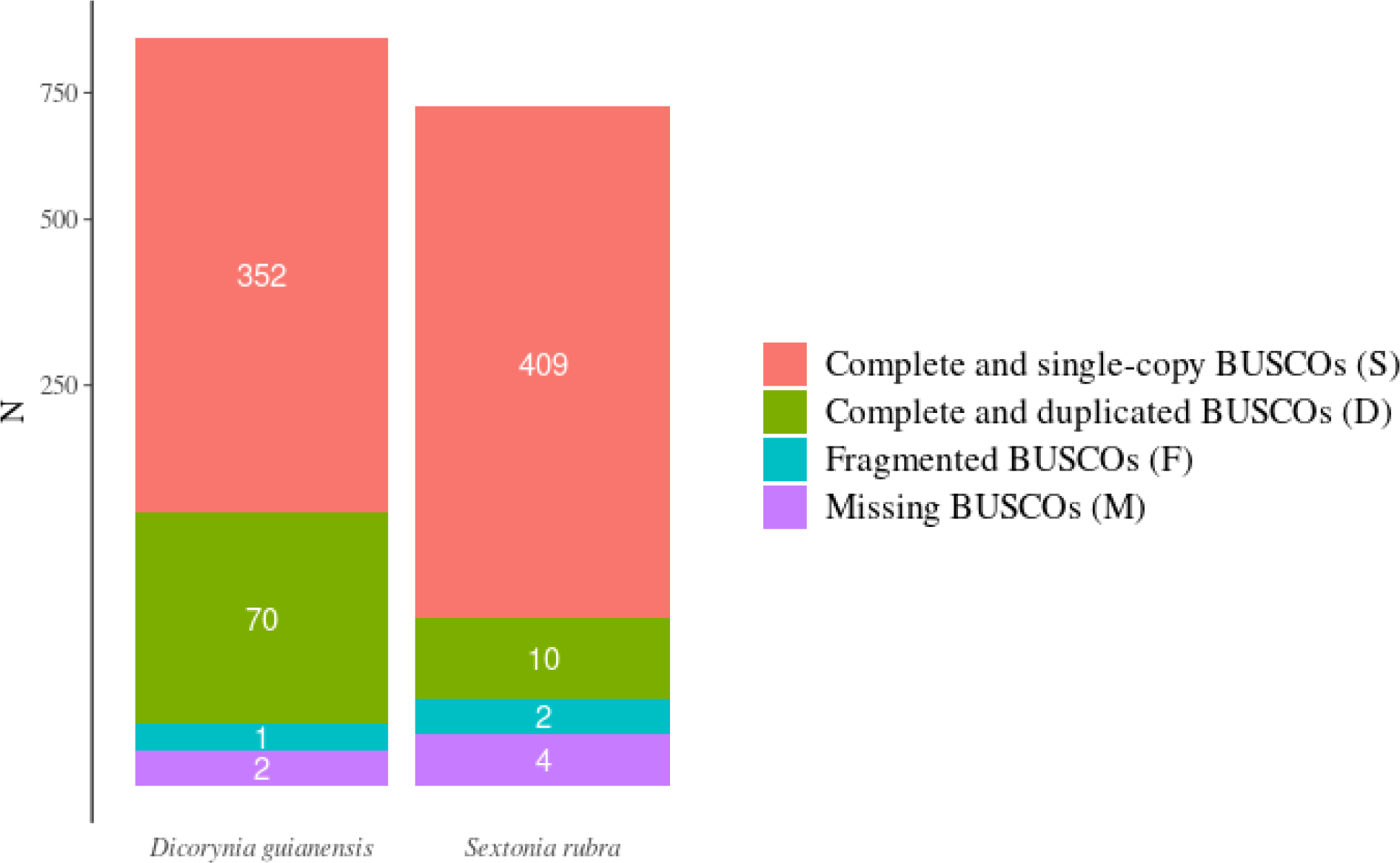
Dicorynia guianensis *and* Sextonia rubra *genome completeness assessed by BUSCO analyses (Seppey e al., 2019) with Viridiplantae housekeeping genes*.

**Figure D2:**
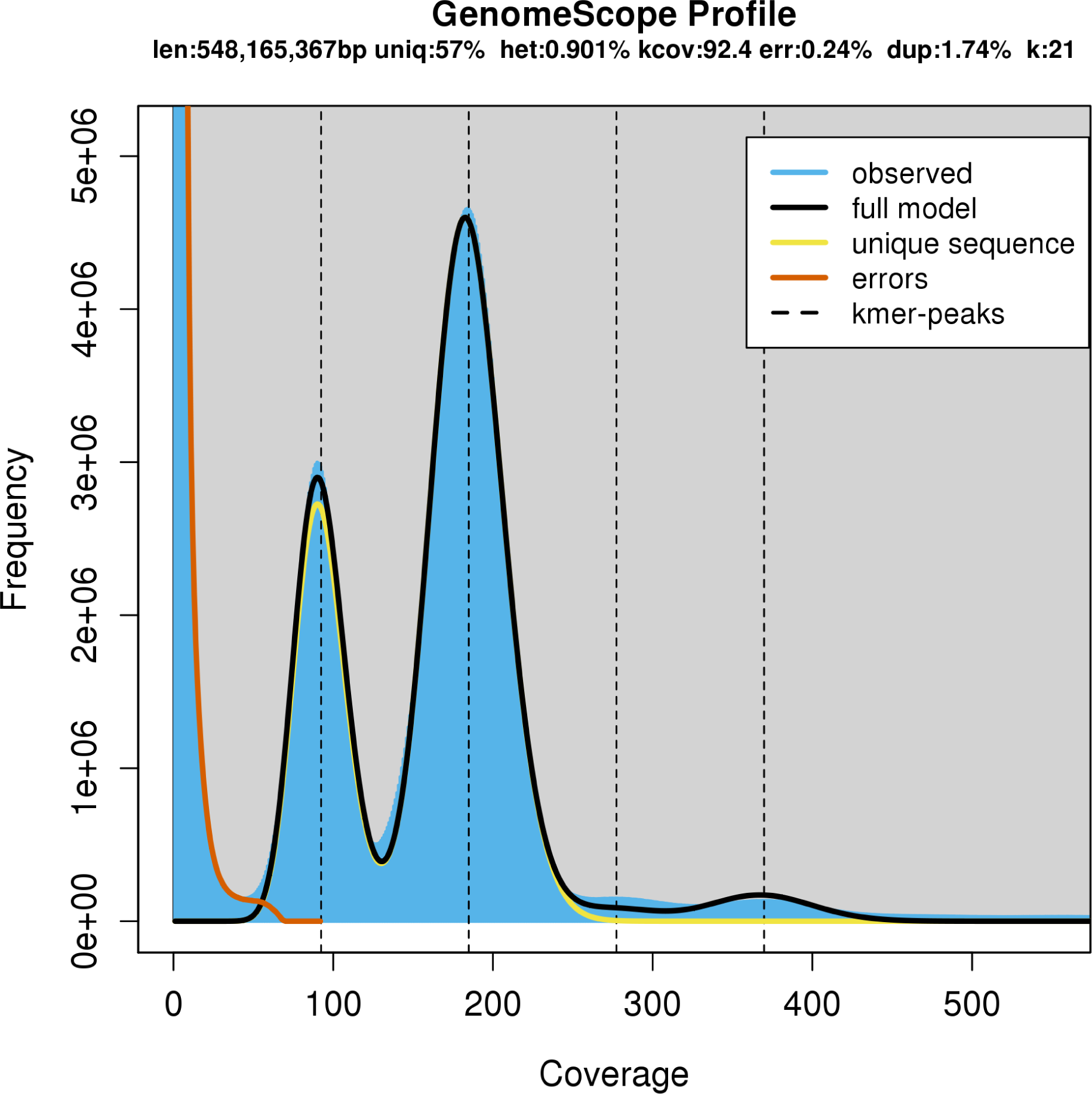
Dicorynia guianensis *heterozygosity estimated at 0.901% with a K-mer based method using Jellyfish (v1.1.12; Marçais and Kingsford 2011) and the GenomeScope (Vurture et al., 2017) with up to 21-mer*.

**Figure D3:**
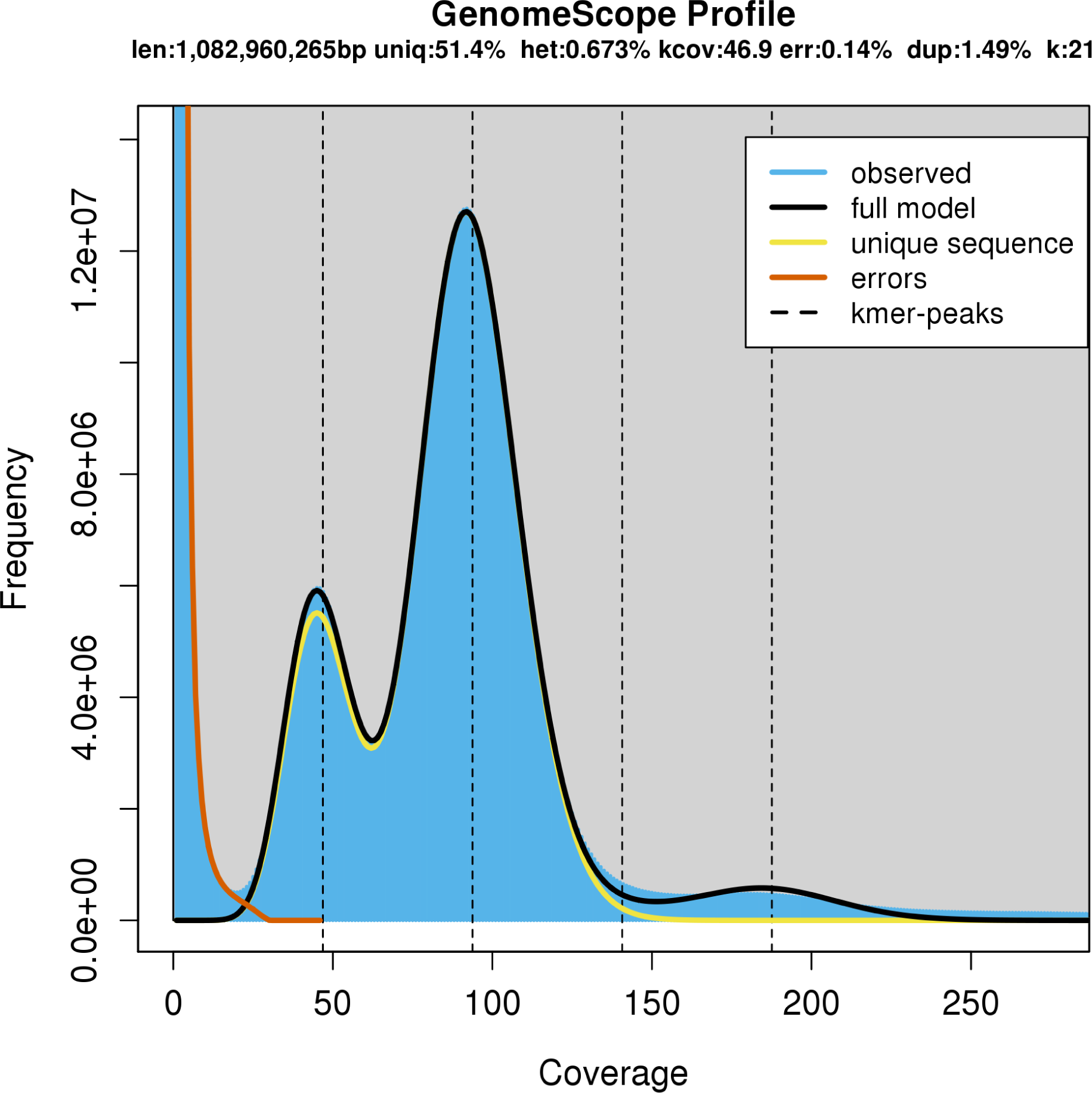
Sextonia rubra *heterozygosity estimated at 0.673% with a K-mer based method using Jellyfish (v1.1.12; Marçais and Kingsford 2011) and the GenomeScope (Vurture et al., 2017) with up to 21-mer*.

**Figure D4:**
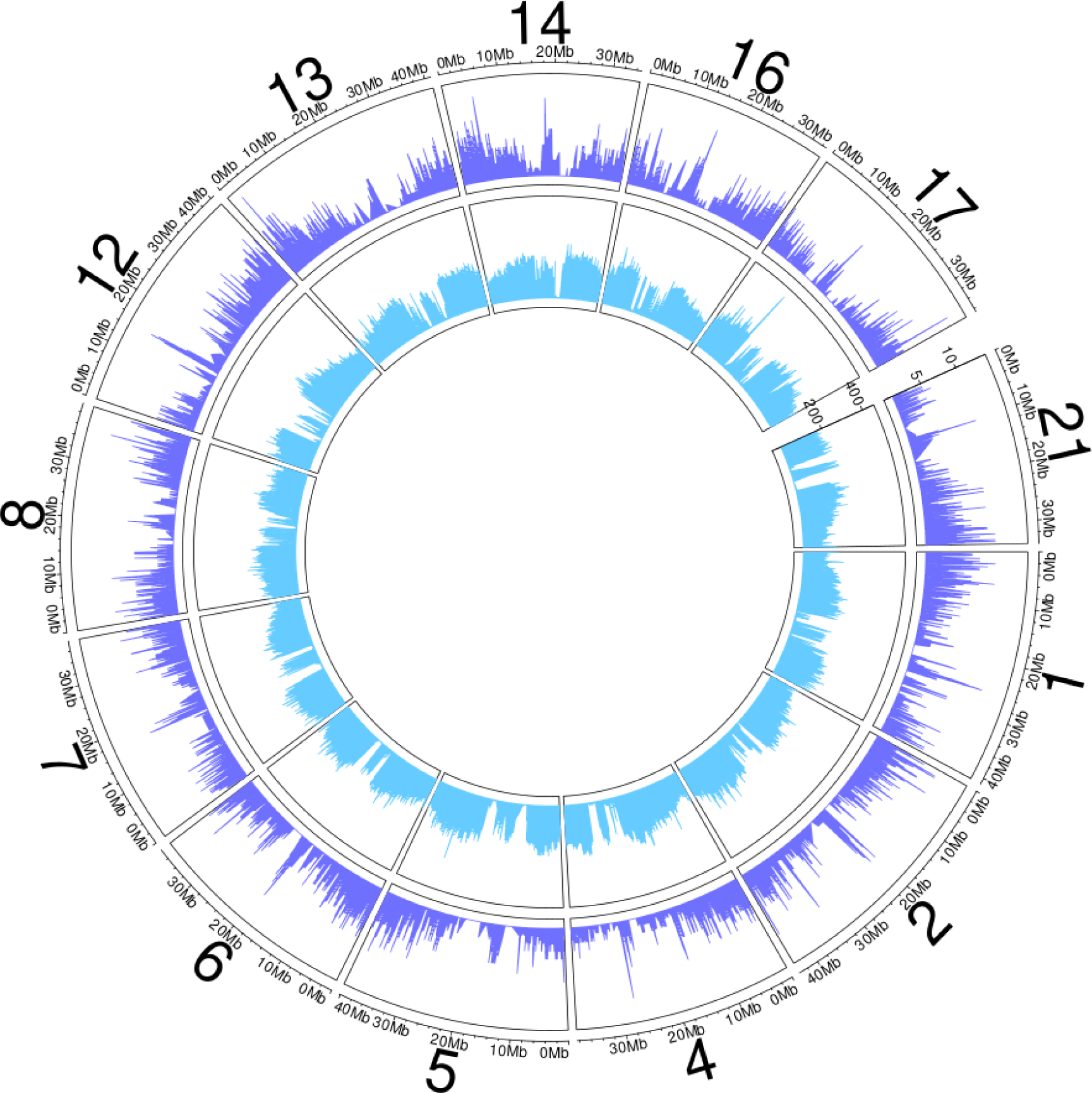
Transposable elements (TE) and gene distribution across the genome of *Dicorynia guianensis* on a 0.1MB sliding window. The outer circle shows genes count in purple, and the inner circle shows TE count blue. Number indicates super-scaffolds id. Only scaffolds with a length above 20Mb are shown.

**Figure D5:**
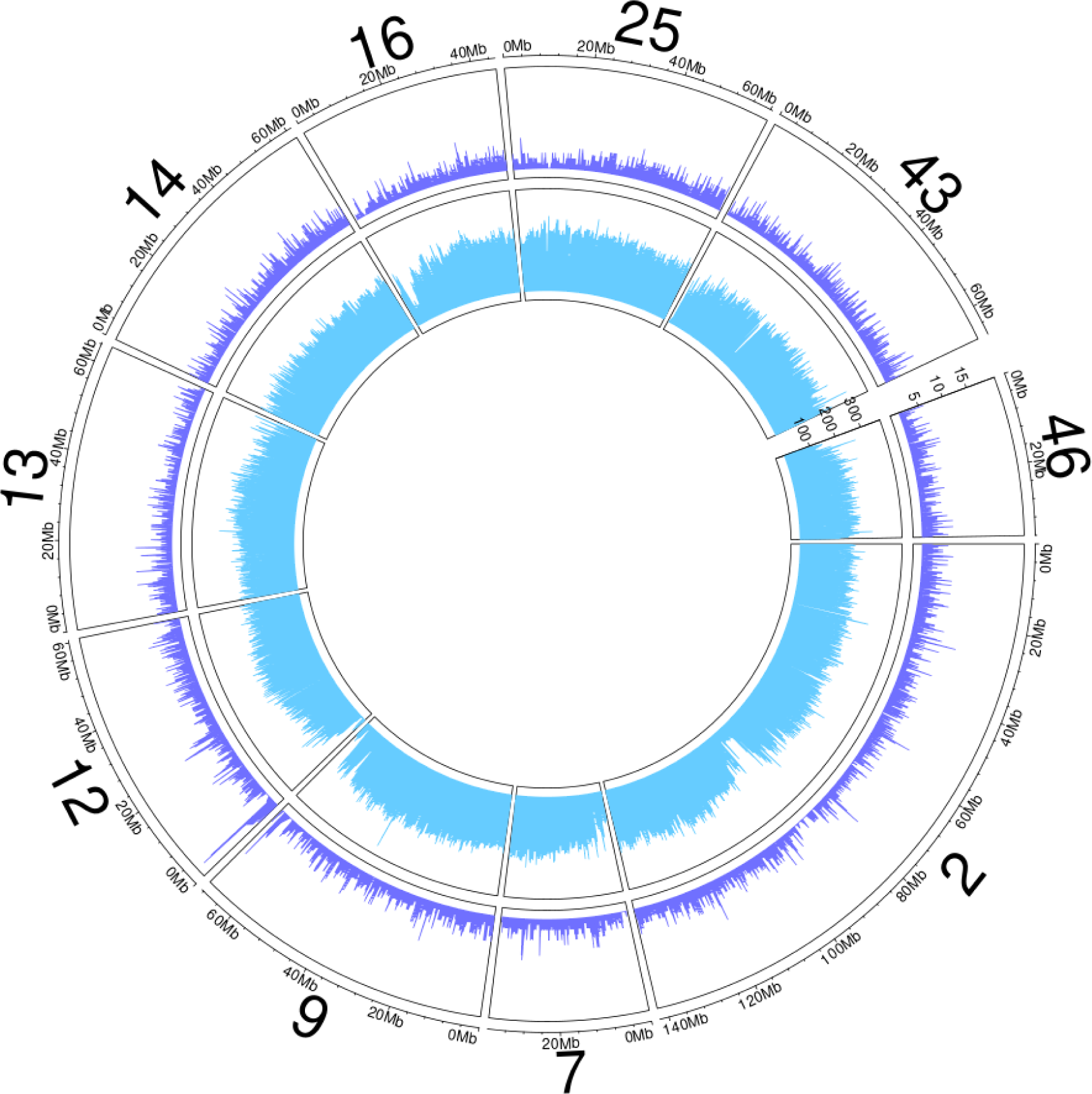
Transposable elements (TE) and gene distribution across the genome of *Sextonia rubra*. on a 0.1MB sliding window. The outer circle shows genes count in purple, and the inner circle shows TE count blue. Number indicates super-scaffolds id. Only scaffolds with a length above 35Mb are shown.

**Figure D6:**
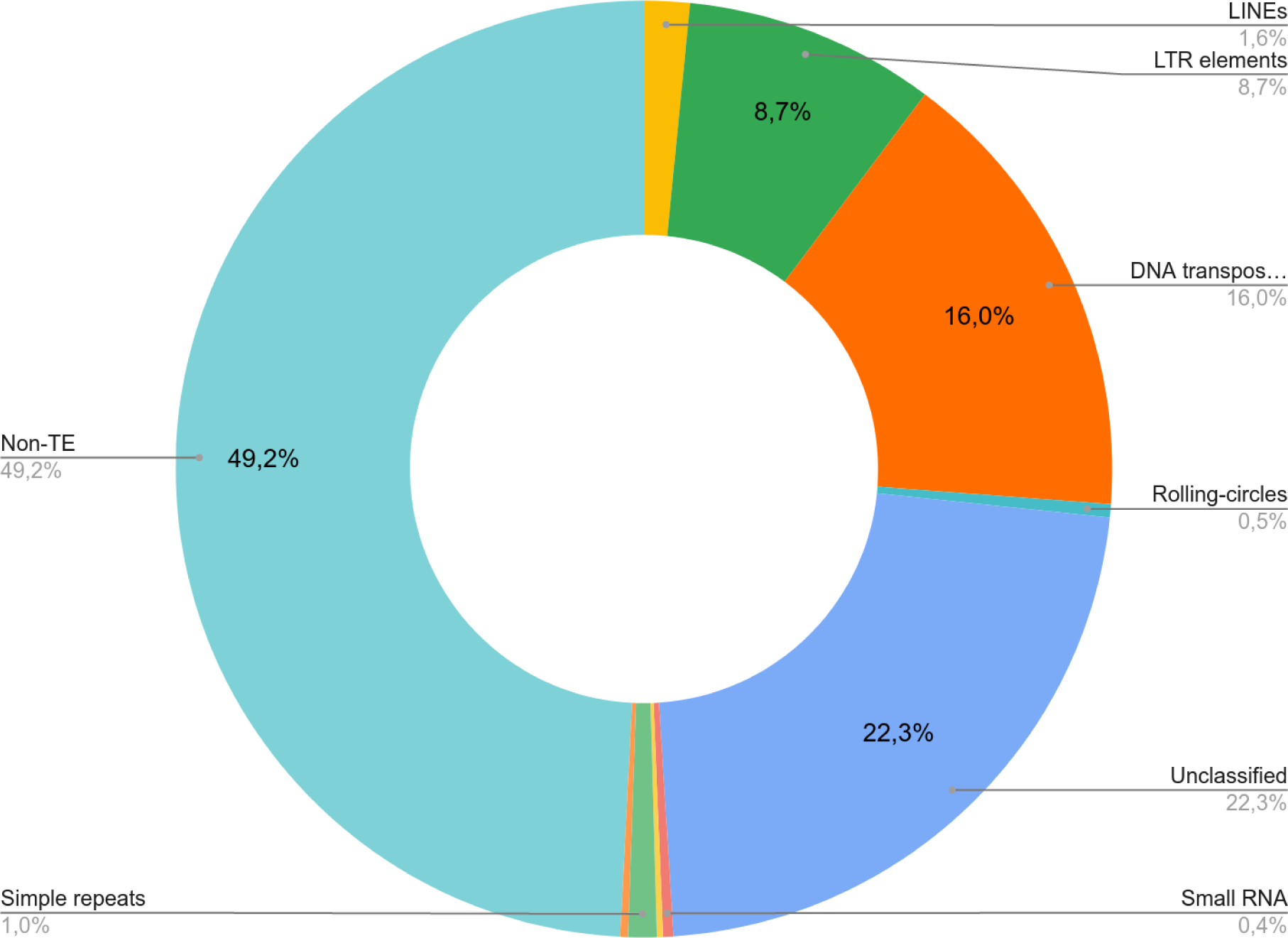
Transposable elements (TE) annotation for *Dicorynia guianensis*. Percentage of genome total length in non-TE, low-complexity DNA, repeats, satellites, small RNA, unclassified TE, rolling circles, DNA transposons, long terminal repeats (LTR) elements, and long interspersed nuclear elements (LINEs). De novo detection of TE resulted in 22.3% of unclassified TE by RepeatClassifier.

**Figure D7:**
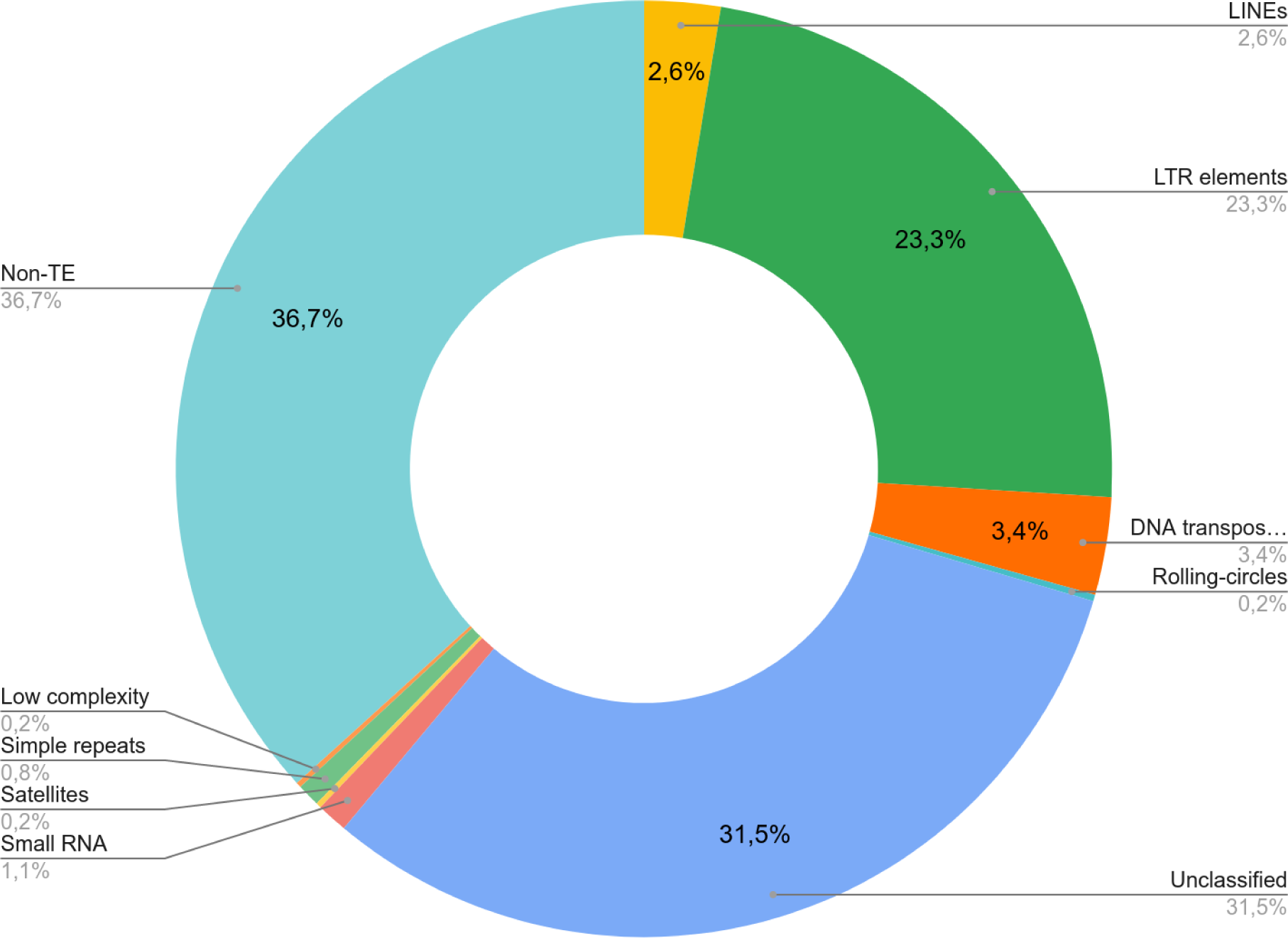
Transposable elements (TE) annotation for *Sextonia rubra*. Percentage of genome total length in non-TE, low-complexity DNA, repeats, satellites, small RNA, unclassified TE, rolling circles, DNA transposons, long terminal repeats (LTR) elements, and long interspersed nuclear elements (LINEs). De novo detection of TE resulted in 22.3% of unclassified TE by RepeatClassifier.

**Figure D8:**
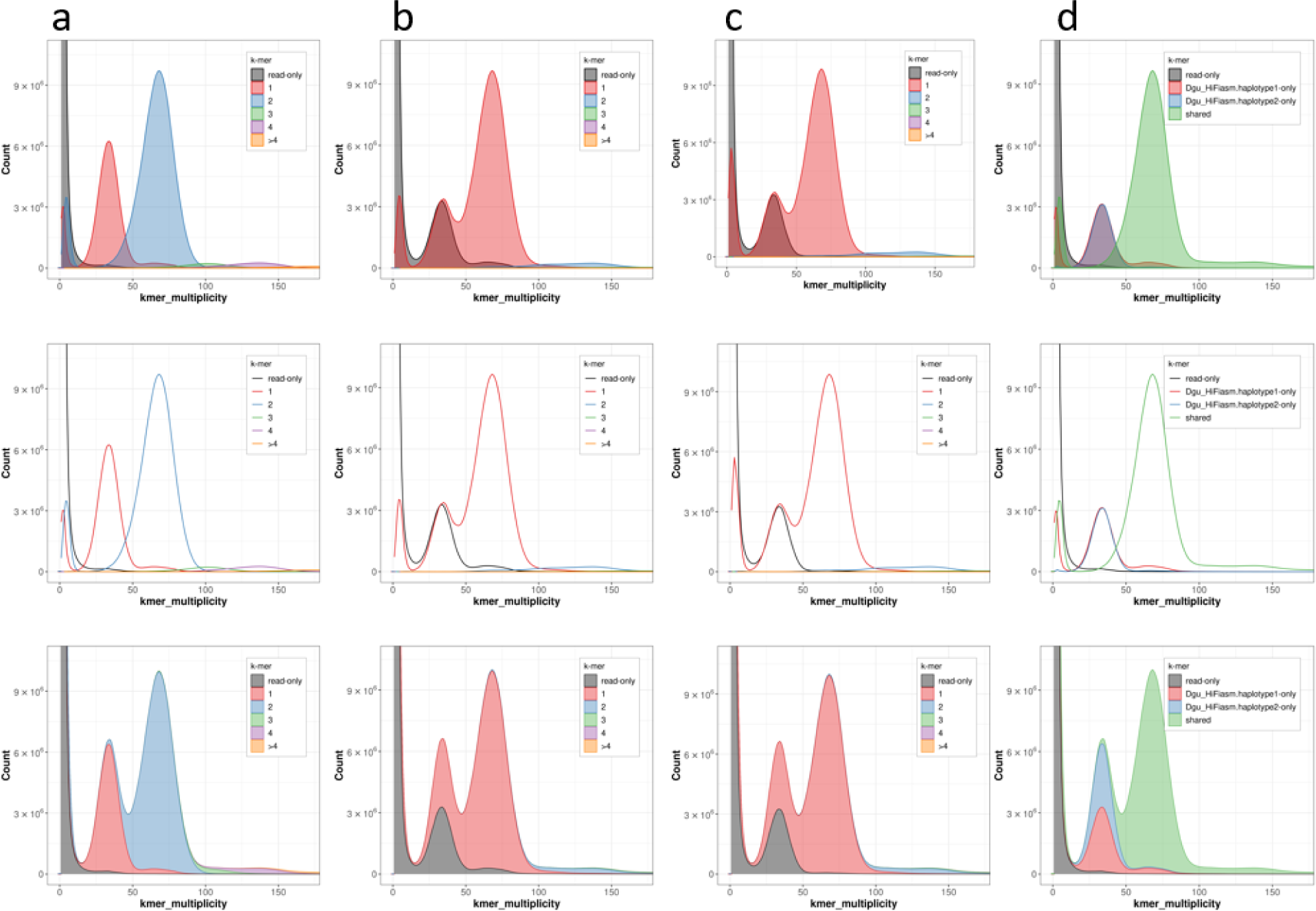
Merqury copy number and assembly spectrum plots for *D. guianensis*. ***a***) Histogram of k-mer multiplicity collected from PacBio reads. The first peak (red) represents 1-copy (heterozygous) kmers in the genome, and the second peak (blue) represents 2-copy k-mers originating from homozygous sequence or haplotype-specific duplications. Depth of sequencing coverage determines where these peaks appear. In this example, sequencing coverage is approximately 70x, corresponding to the 2-copy peak. **b and c**) Spectra-cn plot of haplotype 1 and 2 assembly respectively. Half the single copy k-mers are missing and found in the other haplotype (black). Two-copy k-mers are found once (red) in each haplotype assembly. **d**) K-mers are coloured by their uniqueness in the reads and haplotype assemblies. Distinct k-mer assembly spectrum (spectra-asm) plot of both hap1 and hap2 assemblies. This plot shows the unique (red and blue) and shared portion of k-mers (green).

**Figure D9:**
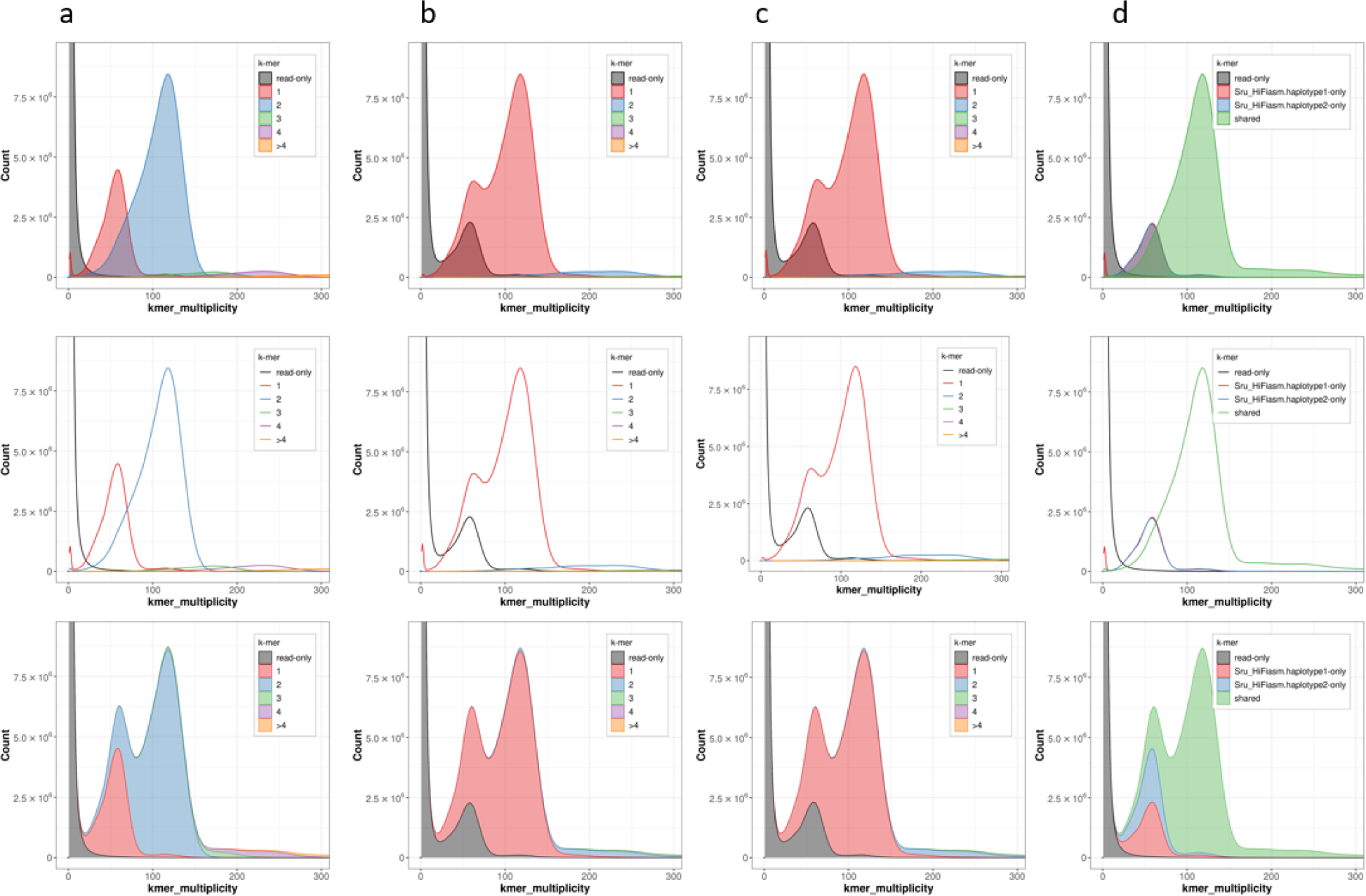
Merqury copy number and assembly spectrum plots for *S. rubra*. ***a***) Histogram of k-mer multiplicity collected from PacBio reads. The first peak (red) represents 1-copy (heterozygous) kmers in the genome, and the second peak (blue) represents 2-copy k-mers originating from homozygous sequence or haplotype-specific duplications. Depth of sequencing coverage determines where these peaks appear. In this example, sequencing coverage is approximately 70x, corresponding to the 2-copy peak. **b and c**) Spectra-cn plot of haplotype 1 and 2 assembly respectively. Half the single copy k-mers are missing and found in the other haplotype (black). Two-copy k-mers are found once (red) in each haplotype assembly. **d**) K-mers are coloured by their uniqueness in the reads and haplotype assemblies. Distinct k-mer assembly spectrum (spectra-asm) plot of both hap1 and hap2 assemblies. This plot shows the unique (red and blue) and shared portion of k-mers (green).

**Figure E1:**
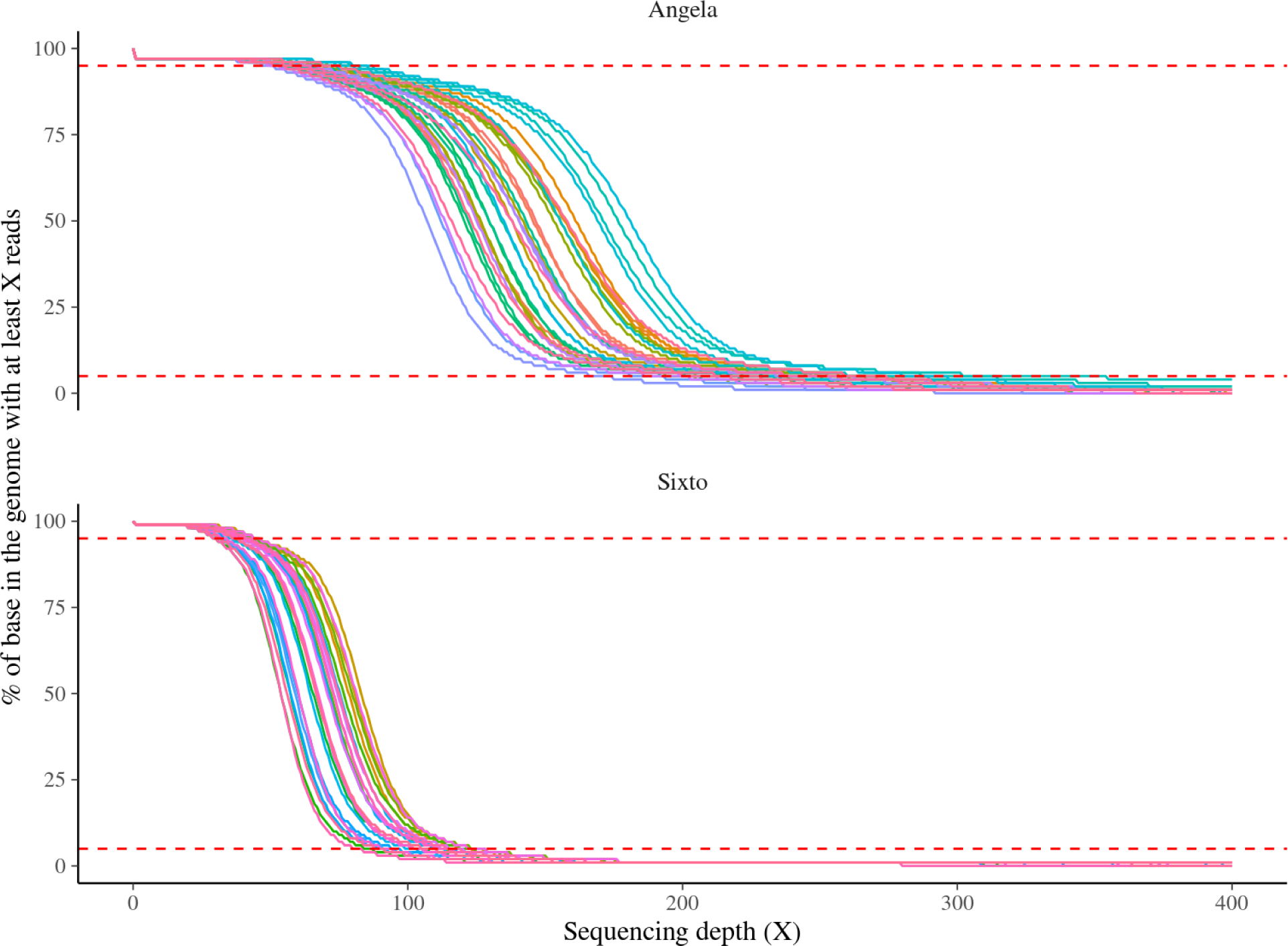
Sequencing depth for Angela and Sixto libraries (cambium and leaves). The x-axis shows the sequencing depth while the y-axis shows the percentage of bases with at least the corresponding sequencing depth. Metrics were calculated over sliding windows of 1 kb.

**Figure E2:**
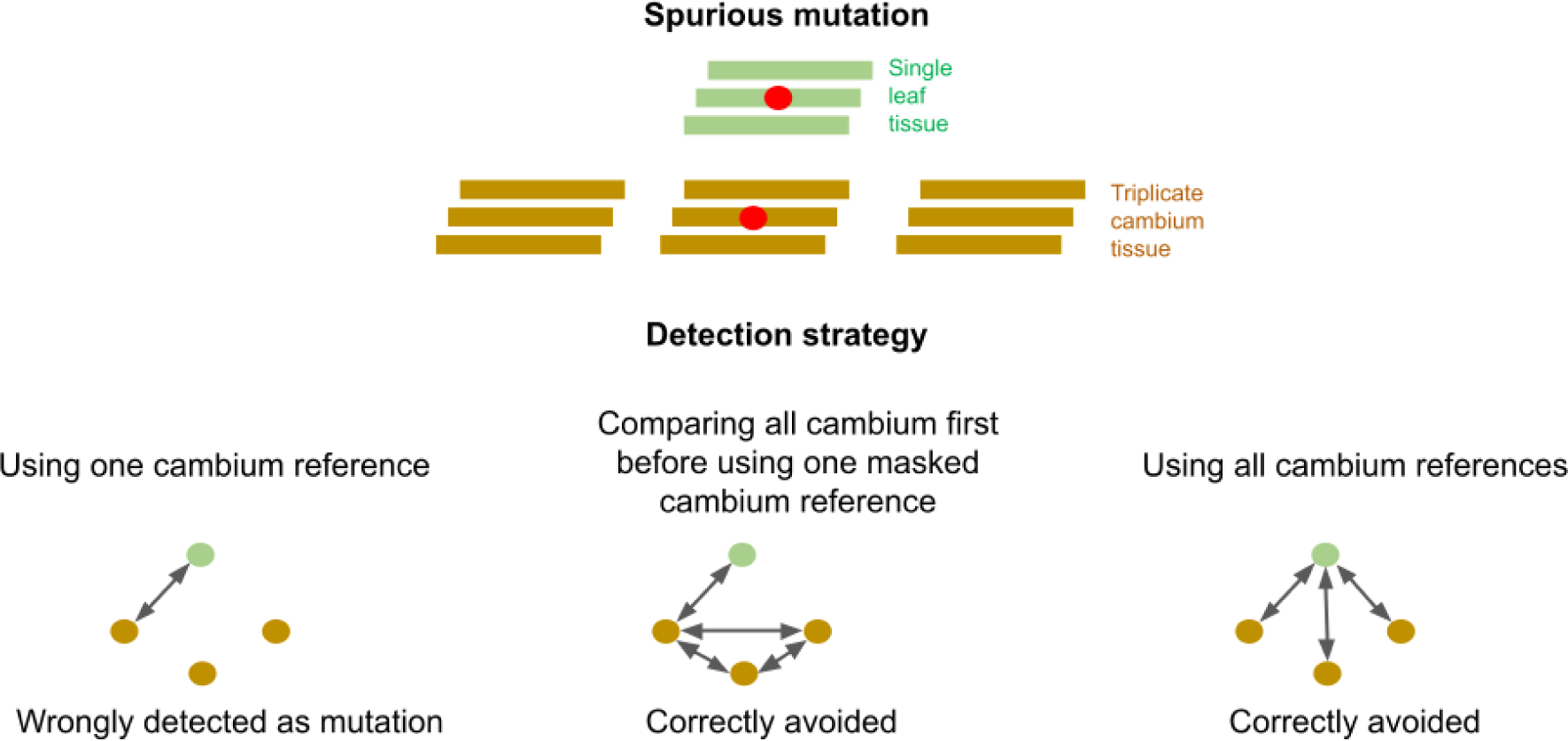
Example of strategies taking advantage of multiple cambial libraries to avoid the detection of spurious mutations. When using only one cambium library (brown) as the normal sample, as shown in the first example, the cambial reads might not include a mutation present in other cambial libraries. Thus the pipeline would wrongly assign a variant discovered in a leaf as a de novo mutation (green). Comparing the leaf library with each cambial library avoids this mistake, as shown in the last example, but takes a lot of computing resources for numerous leaf libraries. Instead, in this study we first compared all cambial libraries against each other to first identify all candidate mutations among cambial libraries, as shown in the second example. We then used the first cambial library as the normal sample against the leaf library, but further masked already identified variant sites from the results to avoid spurious mutations.

**Figure E3:**
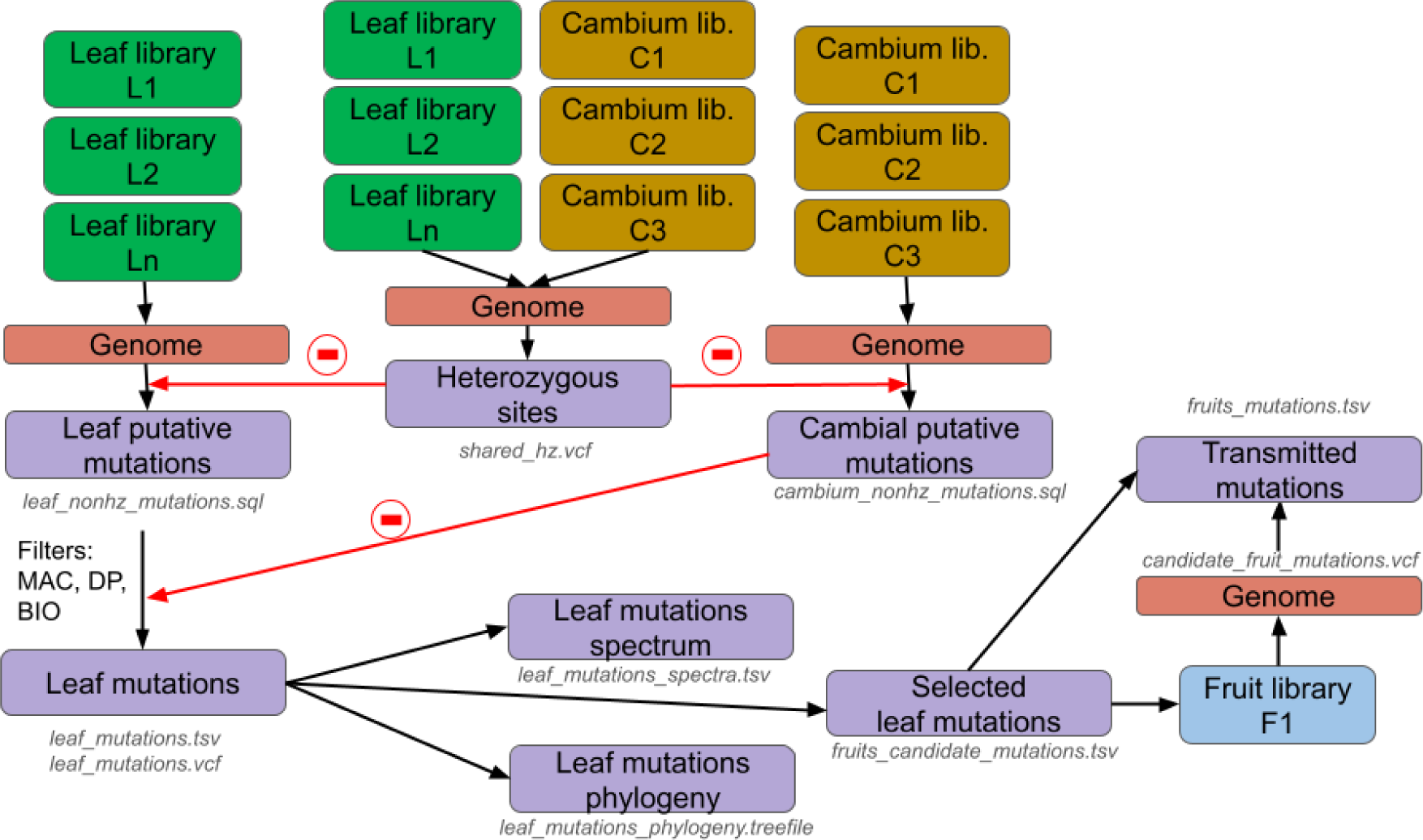
Scheme of the different steps to detect crown mutations and mutations transmitted to fruits with associated resulting files in grey and italic. Heterozygous sites are called for all libraries against the genome (file: shared_hz.vcf). Cambial putative mutations are called for all cambial libraries against the genome and masked for heterozygous sites (file: cambium_nonhz_mutations.sql). Leaf putative mutations are called for all leaf libraries against the genome and masked for heterozygous sites (file: leaf_nonhz_mutations.sql). Leaf mutations are further filtered based on minimum allelic count in the normal sample (MAC=0), sequencing depth of normal and tumour samples (DP) and the existence of at least two biological replicates across the crown (BIO) and masked for putative cambial mutations (file: leaf_mutations in TSV and VCF). Leaf mutations are annotated with their spectra (file: leaf_mutations_spectra.tsv) and their phylogeny built (file: leaf_mutations.treefile). A set of candidates for mutation transmission to fruits is selected (file: fruits_candidate_mutations.tsv) and resequenced using amplicon sequencing. The resulting libraries are aligned against the genomes and the candidate fruit mutations are called (file: candidate_fruit_mutations.vcf) before filtering the candidates according to expected mutations, resulting in the transmitted mutations summarised in the table fruits_mutations.tsv.

**Figure F1:**
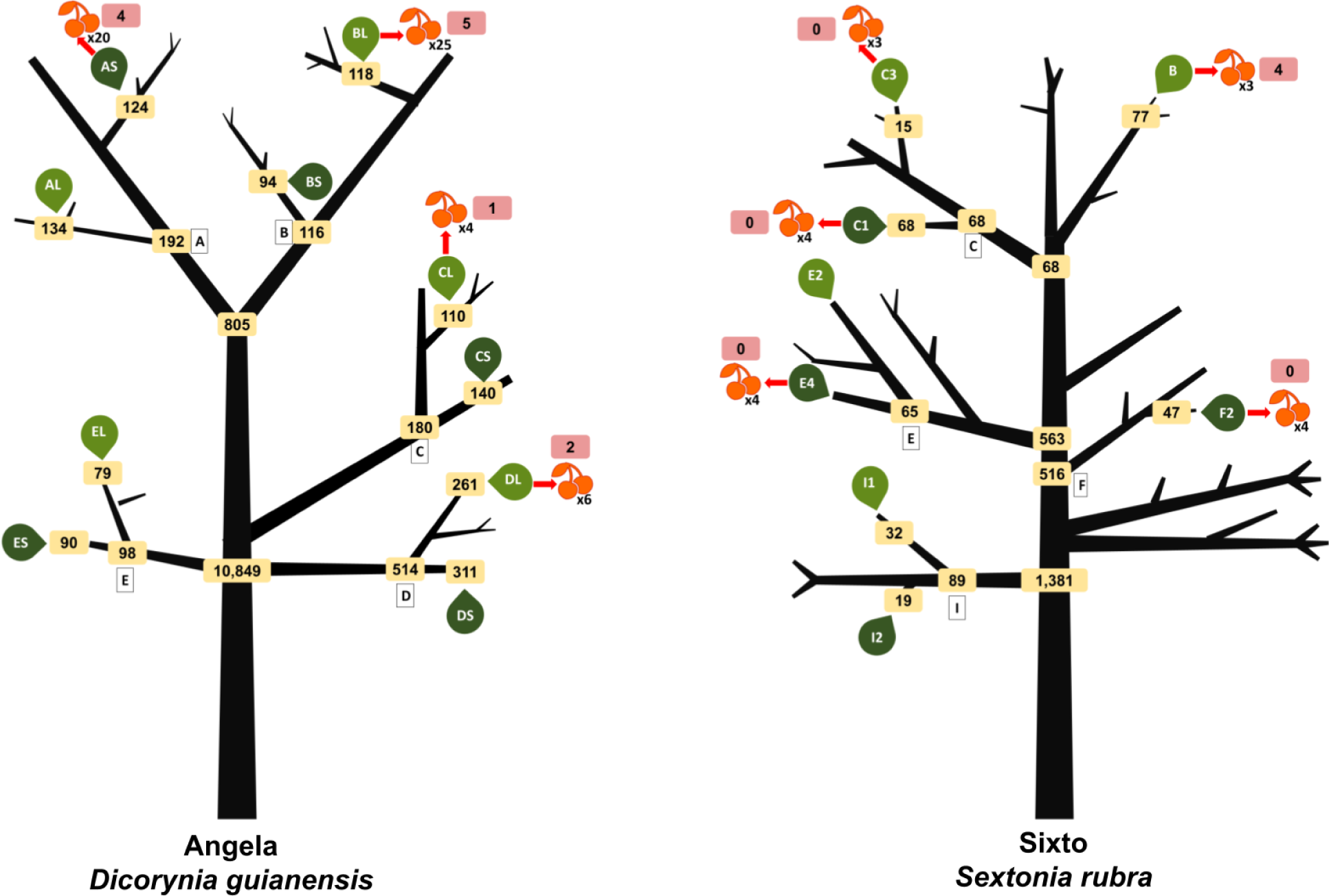
Origin of mutations in the crown of Angela and Sixto. After filtering out mutations with a minimum of 5 copies, no copies in the reference sample, a sequencing depth between the 5^th^ and 95^th^ quantile of the corresponding library and present in at least two samples in the crown, we classified their origin as the most recent branching event of the shared mutation by all samples carrying it. Numbers in yellow boxes give the corresponding number of mutations originating from this branching event.

**Figure F2:**
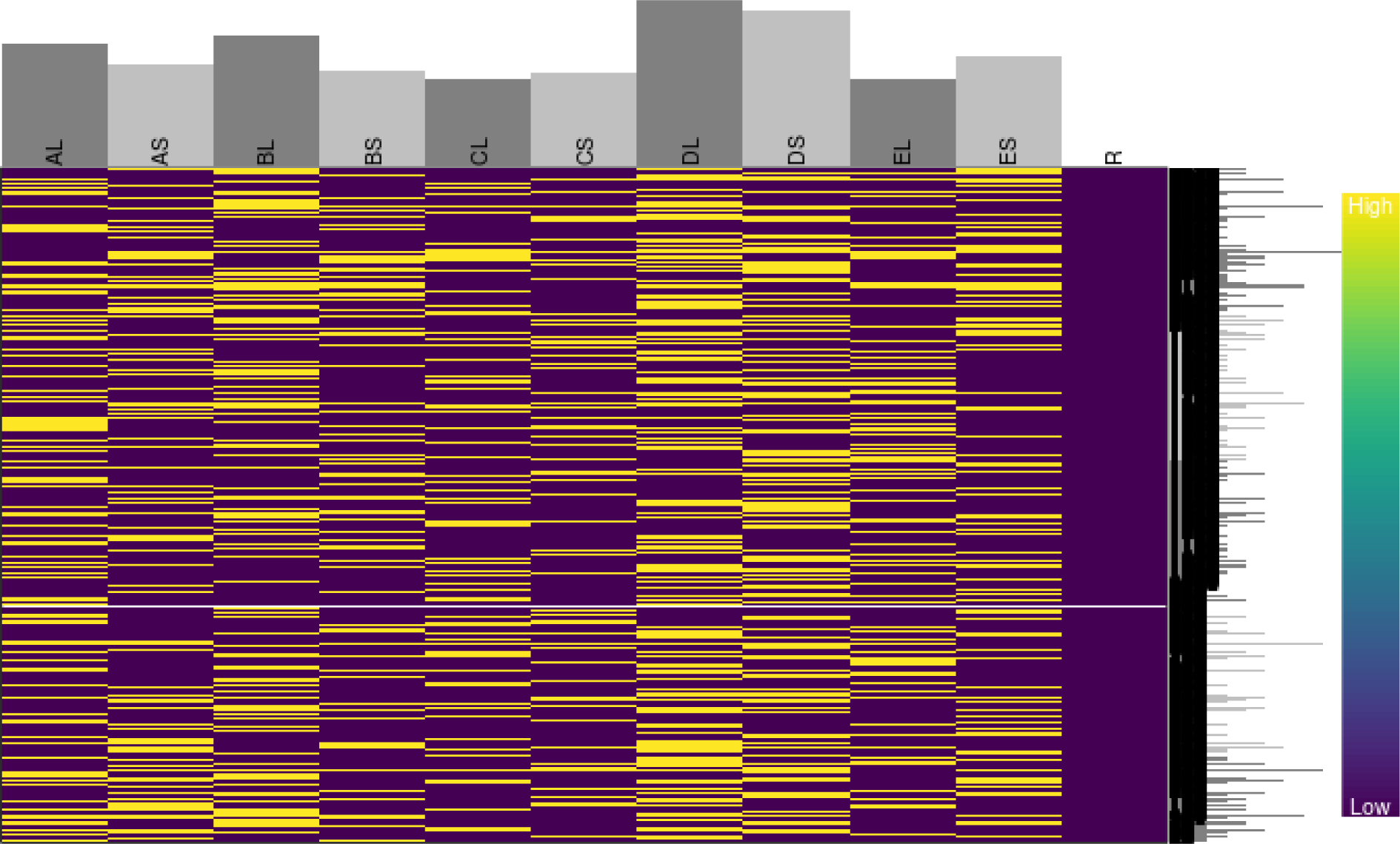
Heat map of somatic mutations on super-scaffold 1 in Angela (lines) in each sample point (lines) with bar graphs representing the total number of accumulated mutations. Yellow represents mutations while purple represents the ancestral state with R representing the fully ancestral root.

**Figure F3:**
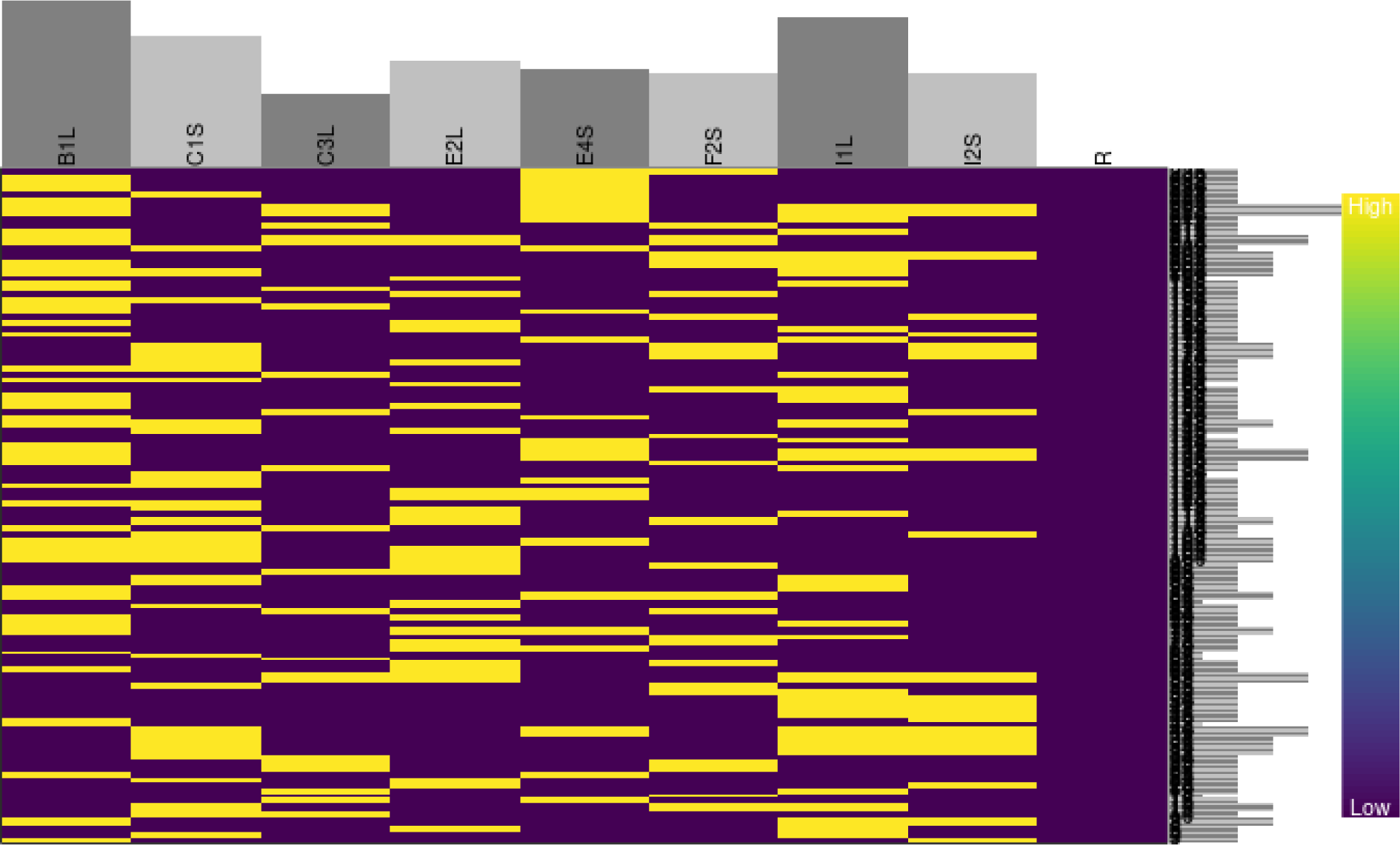
Heat map of somatic mutations on super-scaffold 10 in Sixto (lines) in each sample point (columns) with bar graphs representing the total number of accumulated mutations. Yellow represents mutations while purple represents the ancestral state with R representing the fully ancestral root.

**Figure F4:**
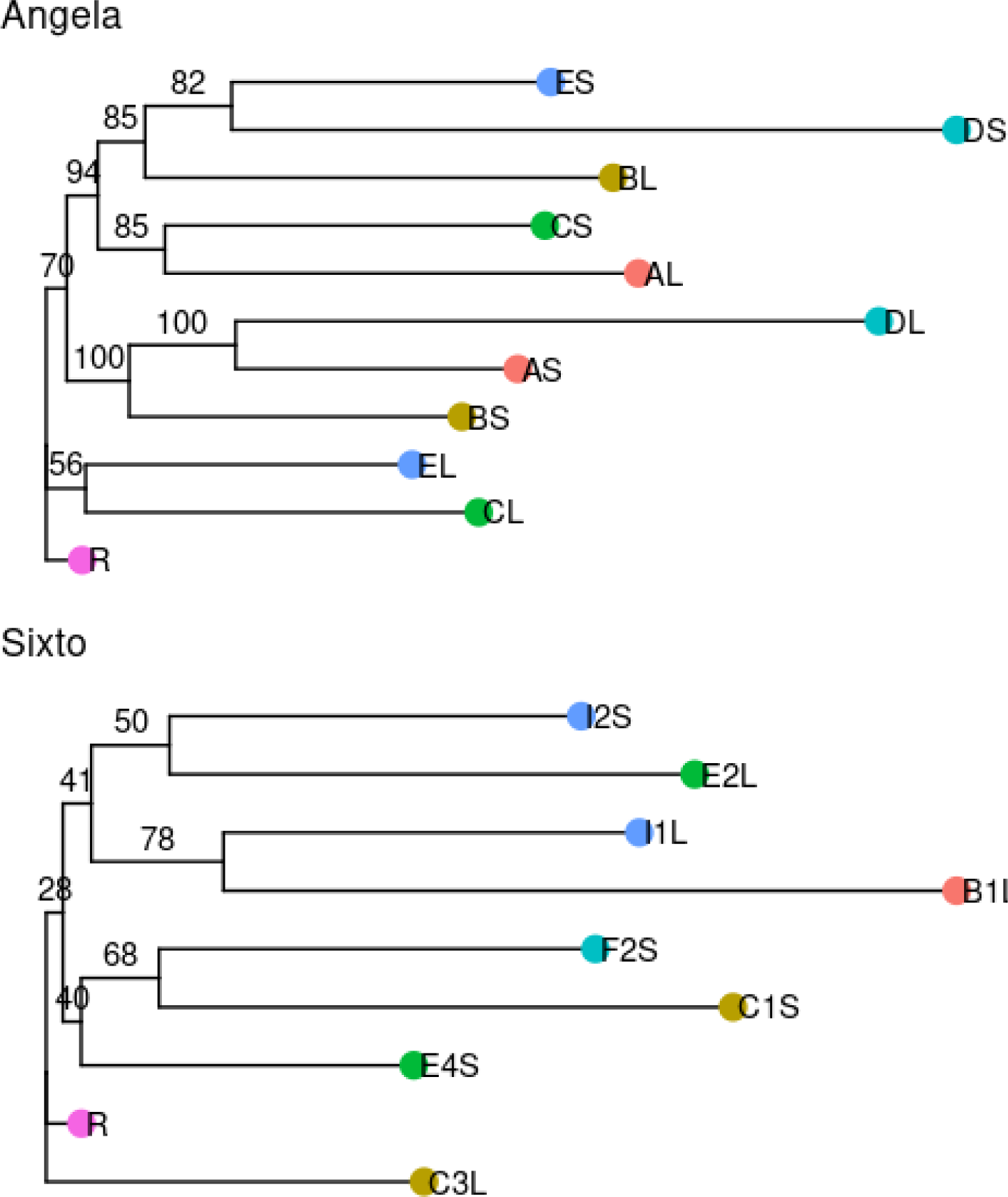
Phylogenies of the Angela and Sixto mutations. The phylogeny of the mutations was constructed with iqtree (Nguyen et al., 2015) with R defined as the root. Node labels indicate node confidence in percent assessed with 1000 bootstraps in iqtree.

**Figure F5:**
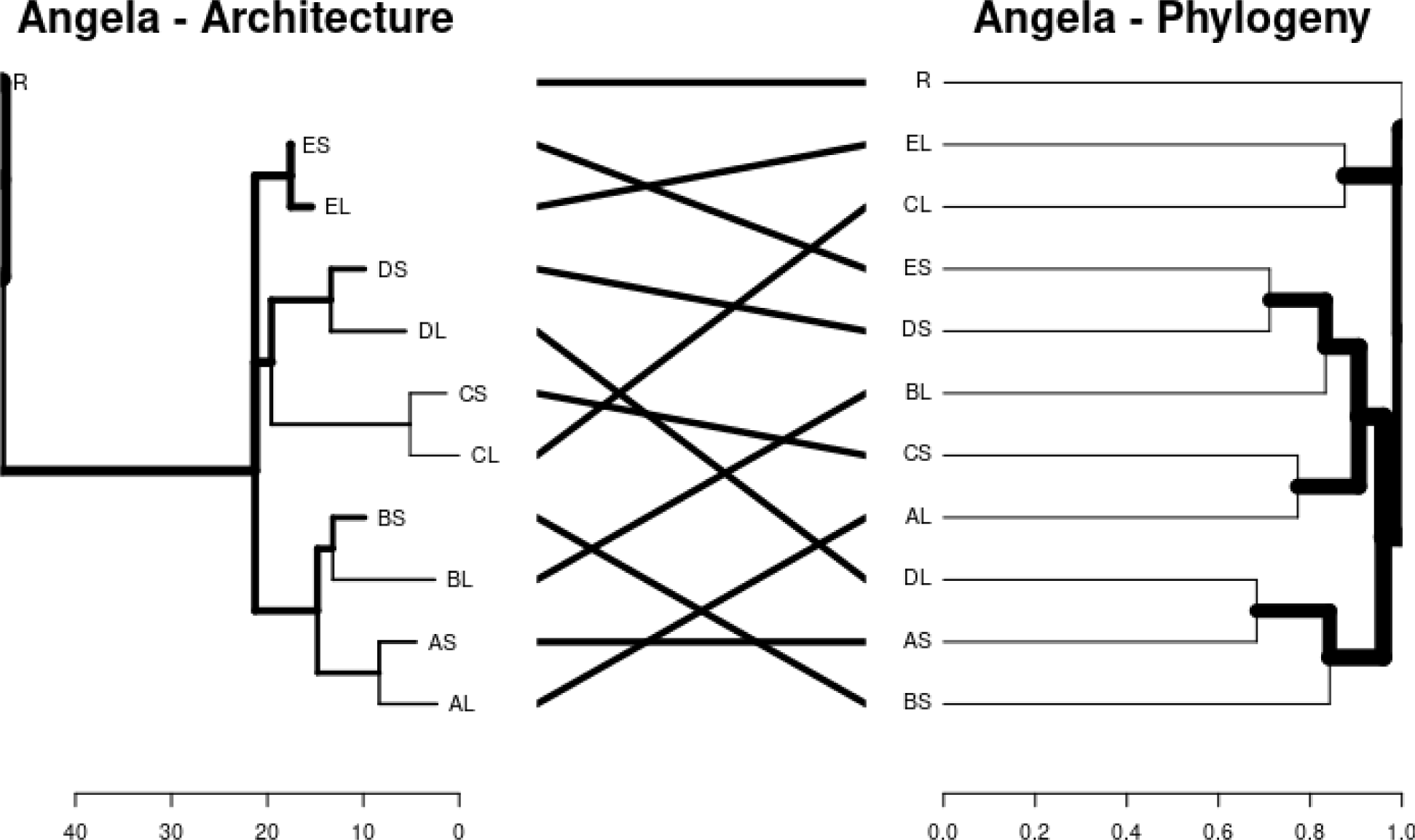
Comparison of the dendrograms of Angela’s architecture with the phylogeny of Angela’s mutations. X-axes represent distances in metres for the architecture and in substitution per base for the phylogeny.

**Figure F6:**
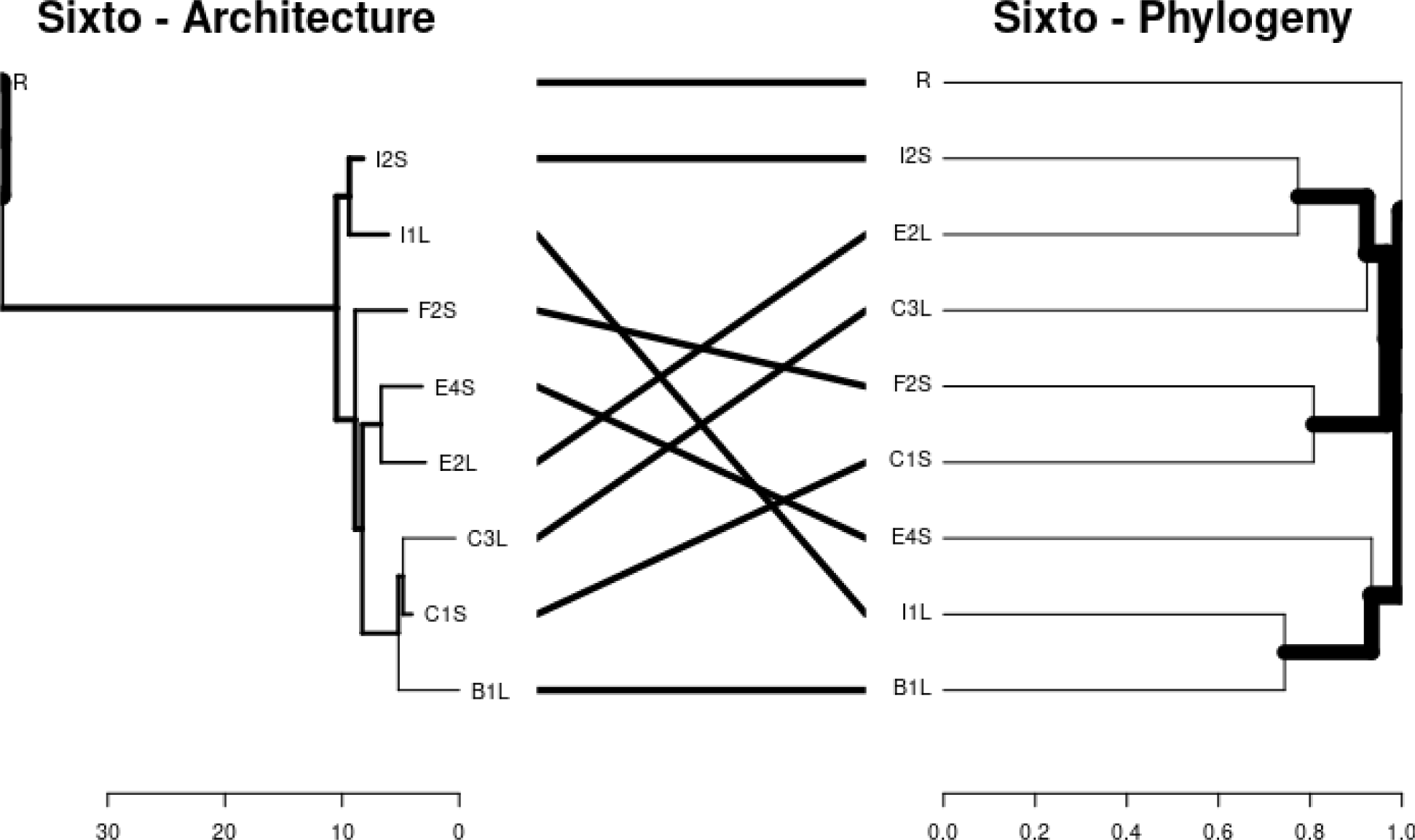
Comparison of the dendrograms of Sixto’s architecture with the phylogeny of Sixto’s mutations. X-axes represent distances in metres for the architecture and in substitution per base for the phylogeny.

**Figure F7:**
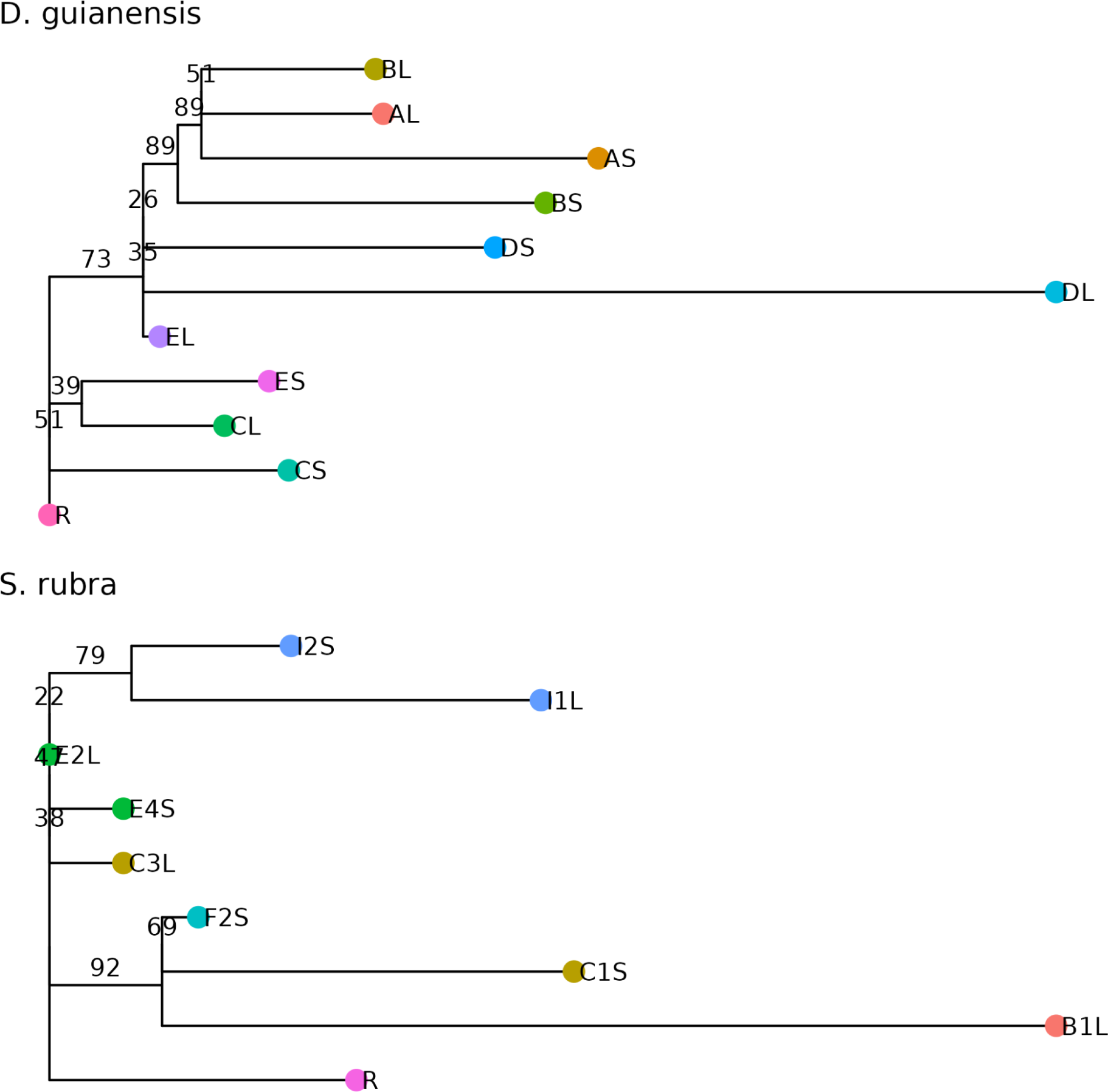
Phylogenies of the Angela and Sixto mutations. for mutations detected in all three leaves of each sampled branch. The phylogeny of the mutations was constructed with iqtree (Nguyen et al., 2015) with R defined as the root. Node labels indicate node confidence in percent assessed with 1000 bootstraps in iqtree.

**Figure F8:**
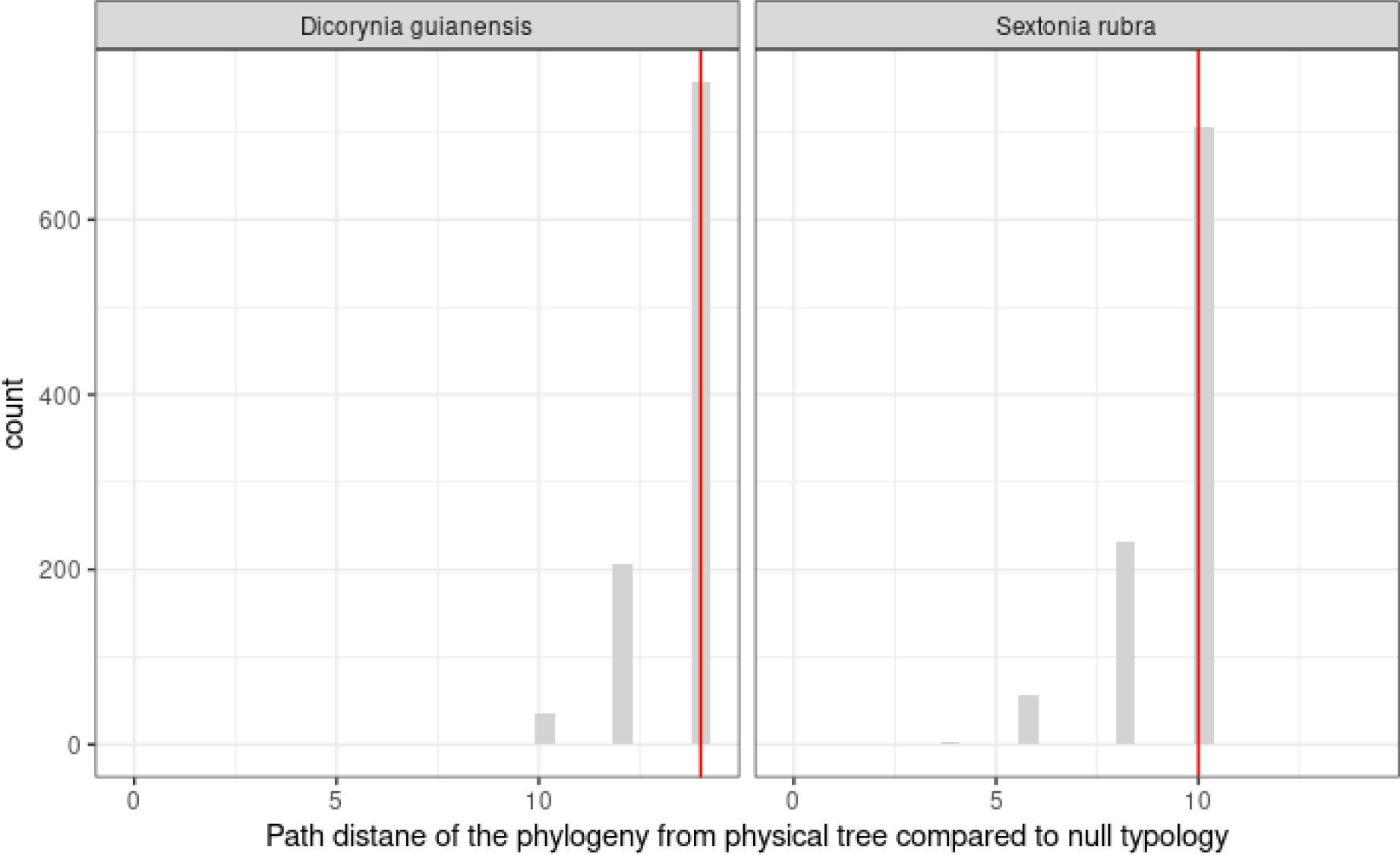
The phylogenetic trees reconstructed from somatic mutations do not resemble the physical structure of the tree more closely than expected by chance. The path distance between the physical tree and 1,000 possible phylogenetic trees is shown with the grey histogram. The path distance between the physical tree and the phylogeny inferred from somatic mutations is shown with the red vertical line.

**Figure G1:**
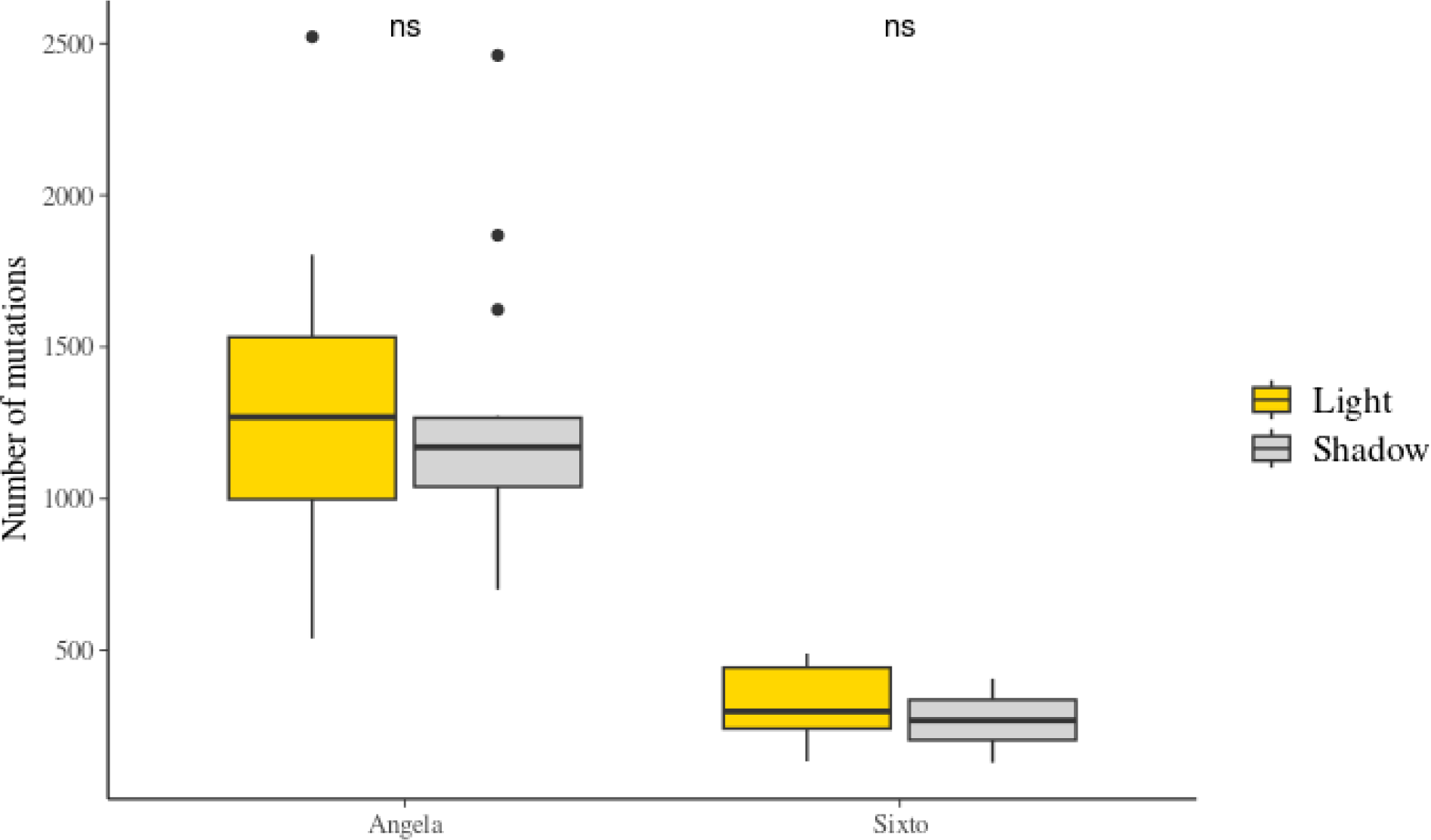
Effect of light exposure on somatic mutation accumulation in Angela and Sixto. Gold represents the number of mutations accumulated in leaves of the light-exposed branches and grey in leaves of the shaded branches. The “ns” labels indicate non-significant differences in the Student’s T-tests.

**Figure G2:**
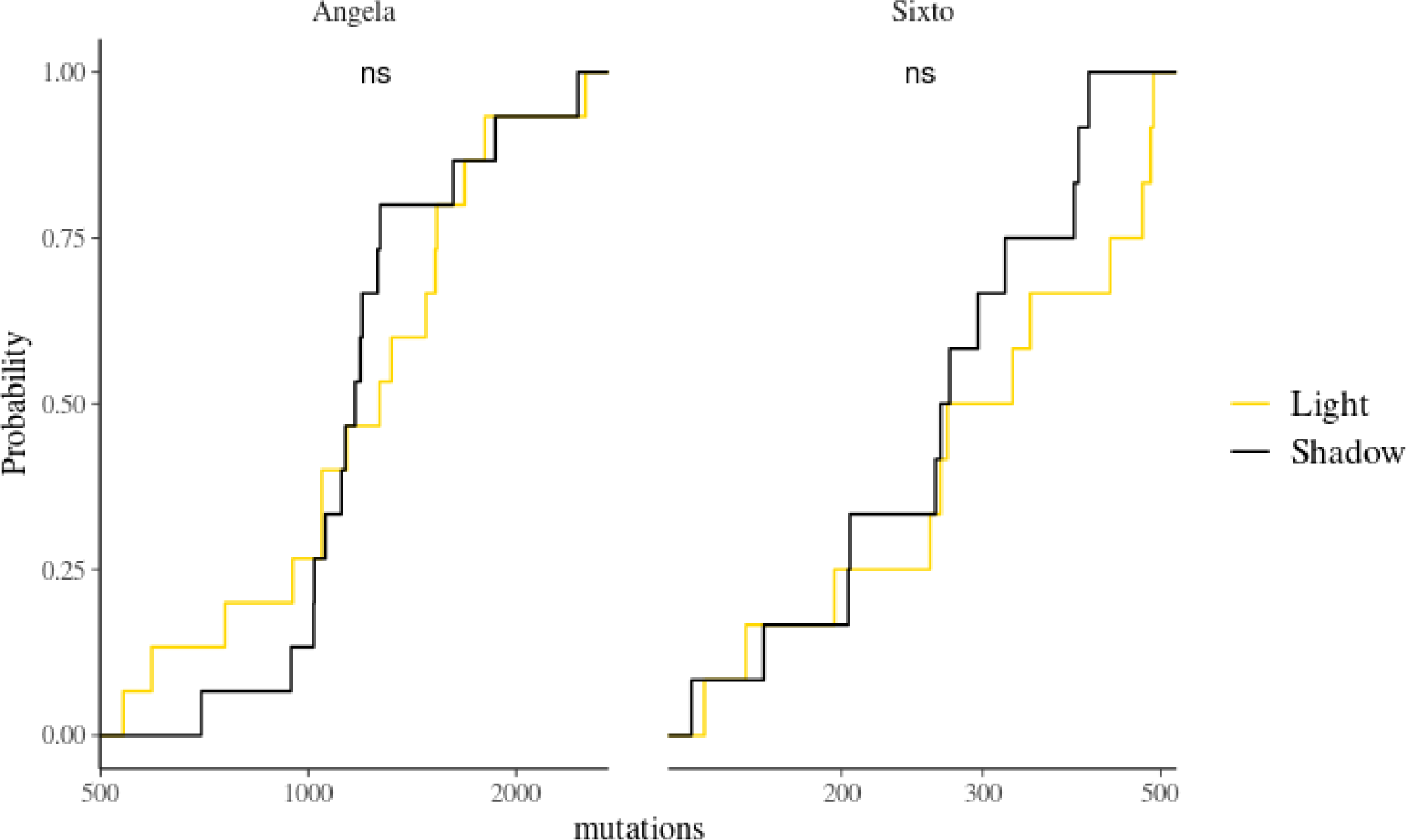
Empirical cumulative distribution of somatic mutations between leaves exposed to light and shade in Angela and Sixto. Gold represents the number of mutations accumulated in leaves of the light-exposed branches and black in leaves of the shaded branches. The “ns” labels indicate non-significant similarity in the Kolmogorov-Smirnov tests.

**Figure G3:**
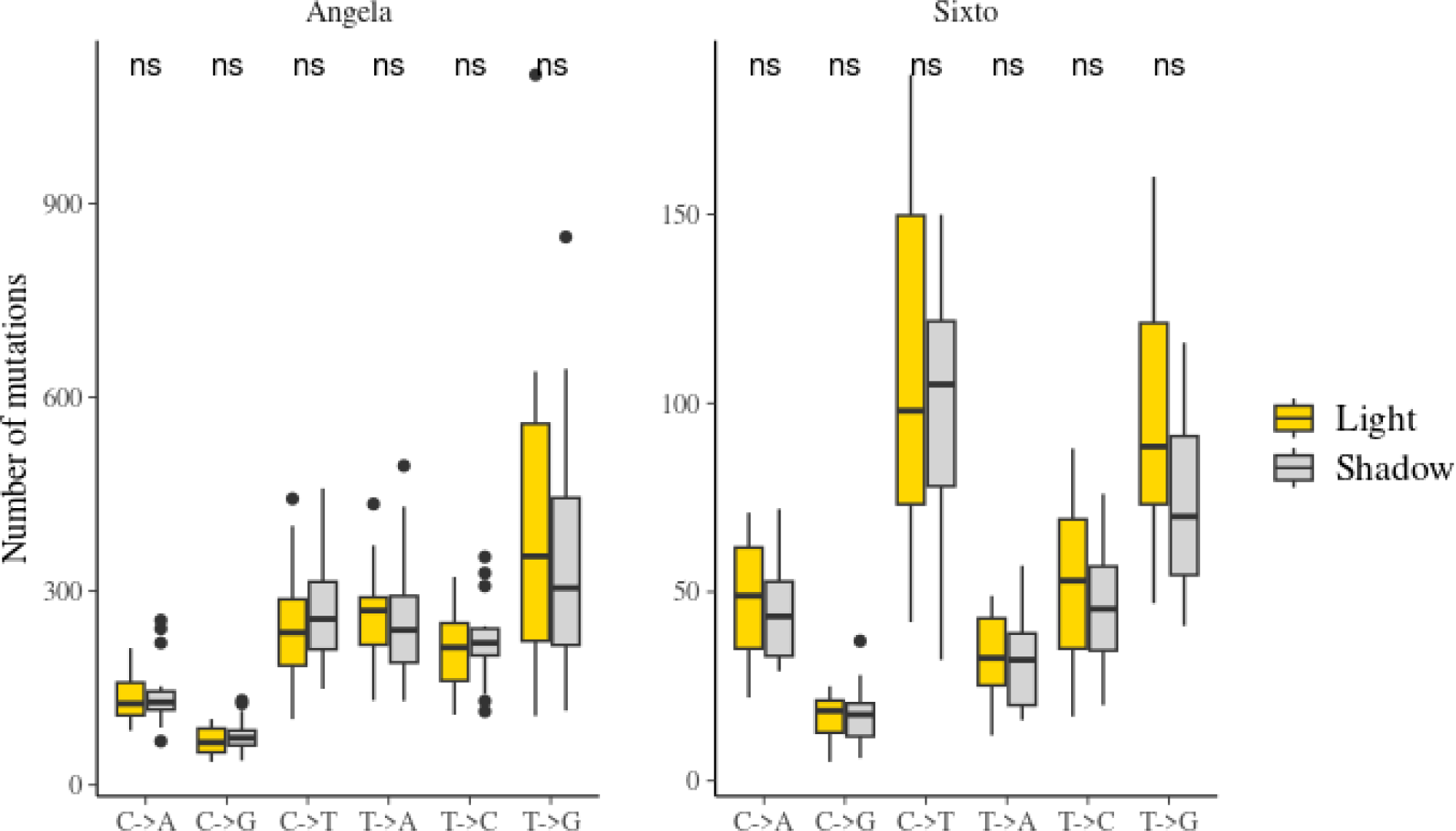
Effect of light exposure on somatic mutation accumulation in Angela and Sixto depending on mutation type. Gold represents the number of mutations accumulated in all leaves of the lighted tips and grey in all leaves of the shaded tips. The “ns” labels indicate non-significant differences in Student’s T-tests.

**Figure G4:**
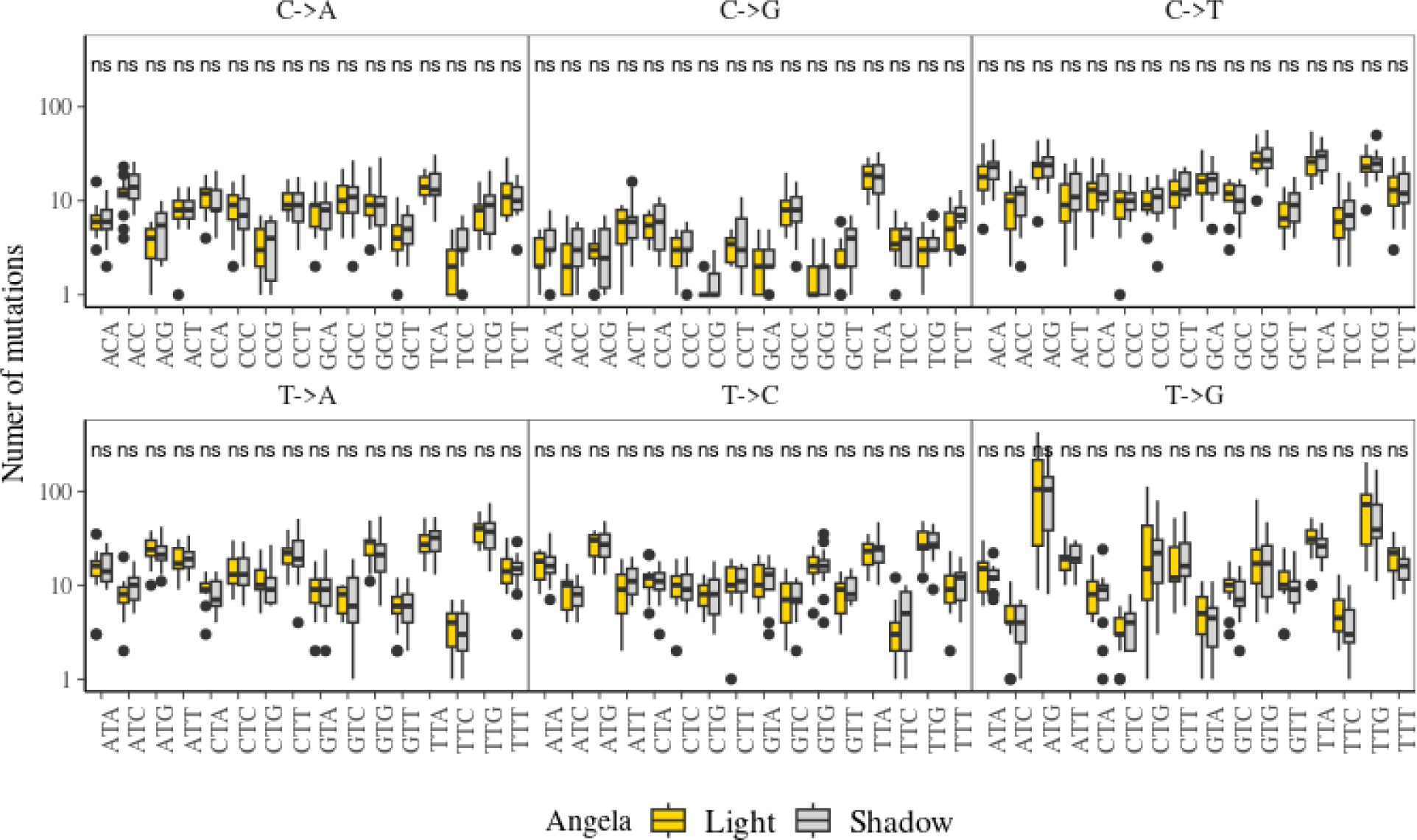
Effect of light exposure on somatic mutation accumulation in Angela depending on mutation spectra (mutation context with 5’ and 3’ bases). Gold represents the number of mutations accumulated in all leaves of the lighted tips and grey in all leaves of the shaded tips. The “ns” labels indicate non-significant differences in Student’s T-tests.

**Figure G5:**
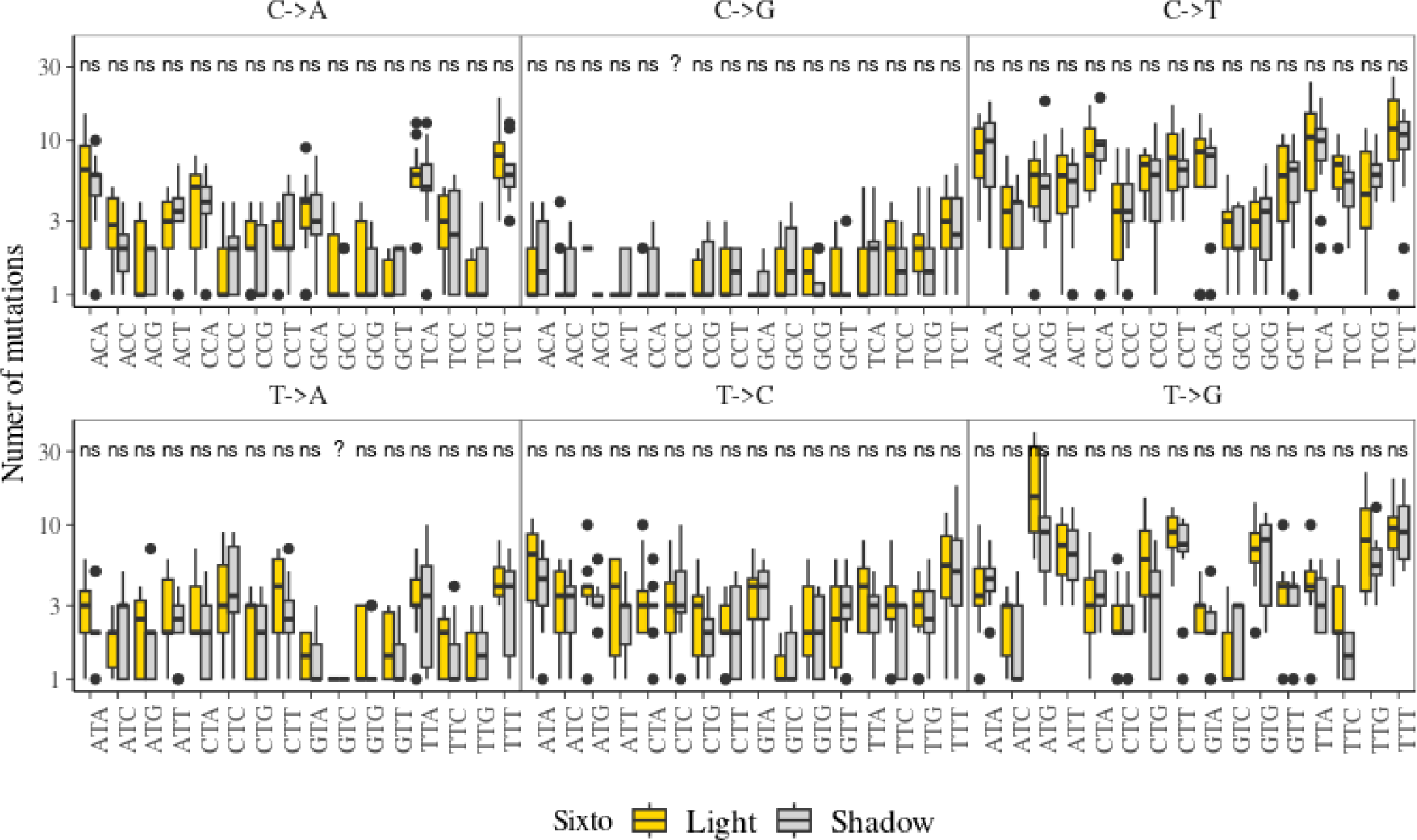
Effect of light exposure on somatic mutation accumulation in Sixto depending on mutation spectra (mutation context with 5’ and 3’ bases). Gold represents the number of mutations accumulated in all leaves of the light-exposed branches and grey in all leaves of the shaded branches. The “ns” labels indicate non-significant differences in Student’s T-tests.

**Figure G6:**
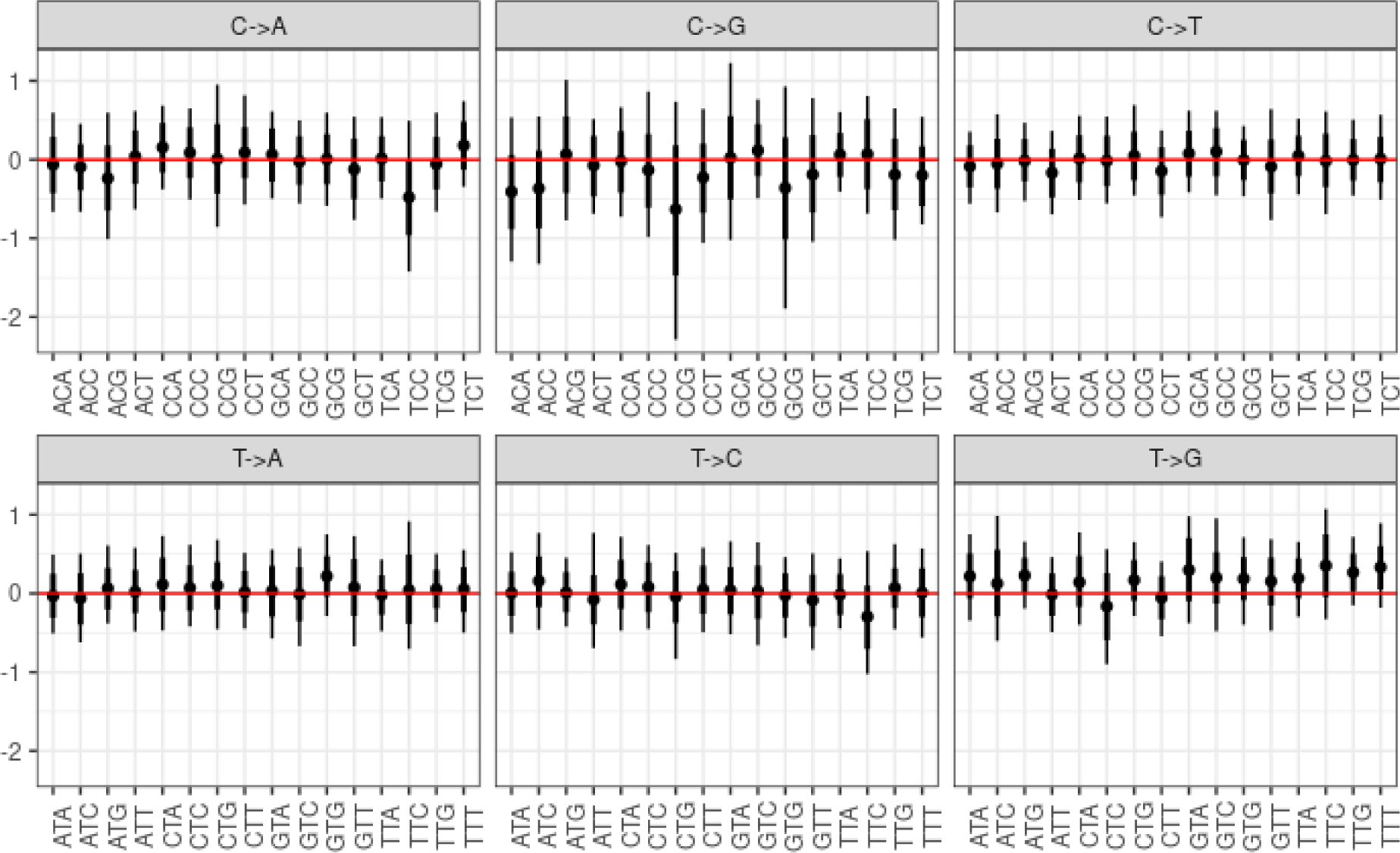
Effect of light exposure on somatic mutation accumulation on mutation type (X>X) spectra (mutation context with 5’ and 3’ bases). We used a Dirichlet multinomial model to test the effect of light on the number of mutations accumulated per spectra. The posterior distribution of light effect for each spectrum within each species is represented with the point representing the mean value of the parameter posterior, thick lines the 50% confidence interval and thin lines the 95% confidence interval. The red horizontal line represents a null effect intercepted by all posterior distributions and indicates that there is no significant effect of light on spectrum frequencies.

**Figure H1:**
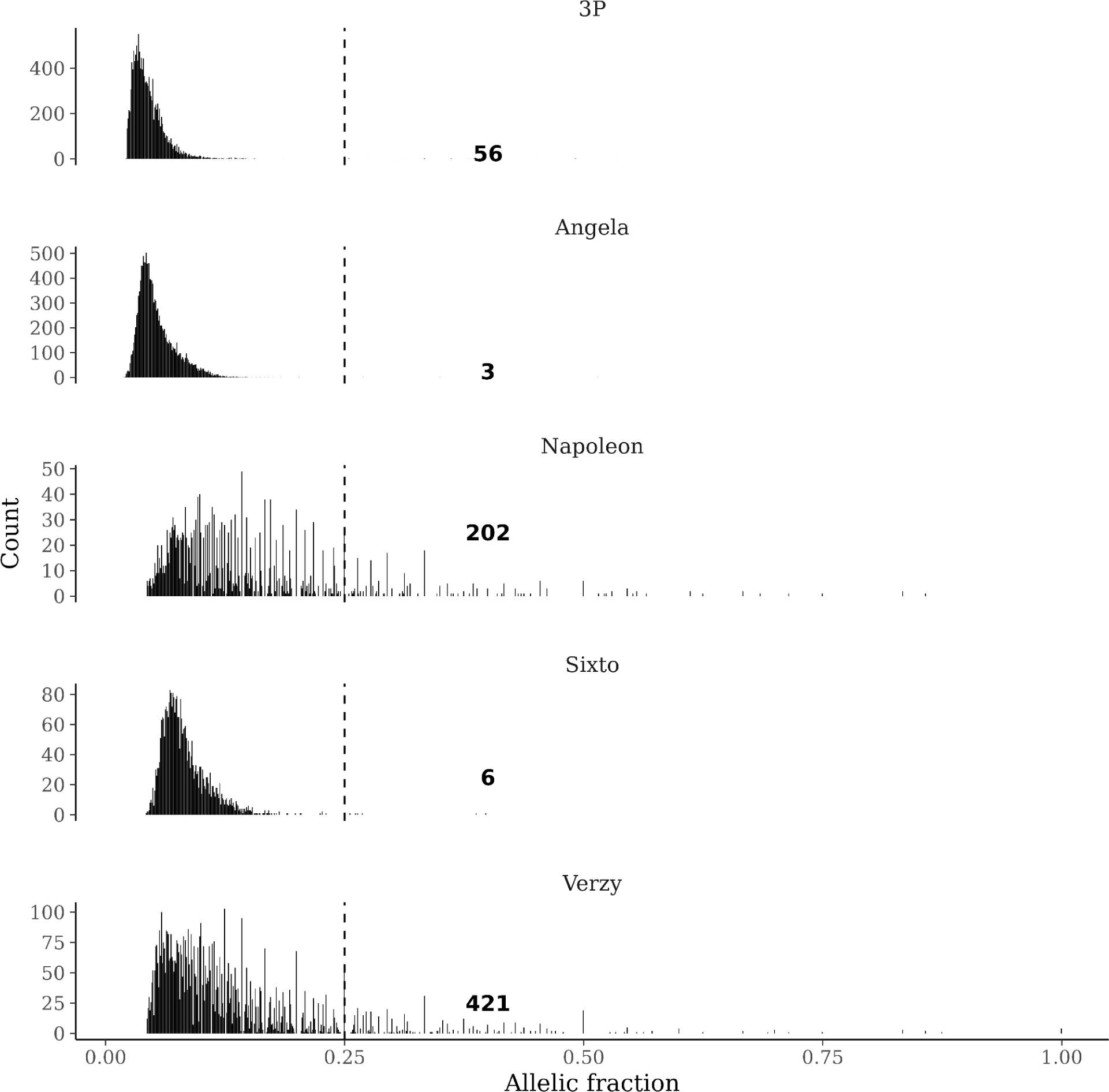
Distribution of allelic fractions for the Angela and Sixto mutations; for the two pedunculate oaks *Quercus robur* reanalysed with the same pipeline: 3P and Napoleon (Schmitt et al., 2022); and an dataset from one tortuous phenotype of common beech *Fagus sylvatica* analysed with the same pipeline. The lower sequencing depth of Napoleon and Verzy explain their smaller left distribution.

**Figure H2:**
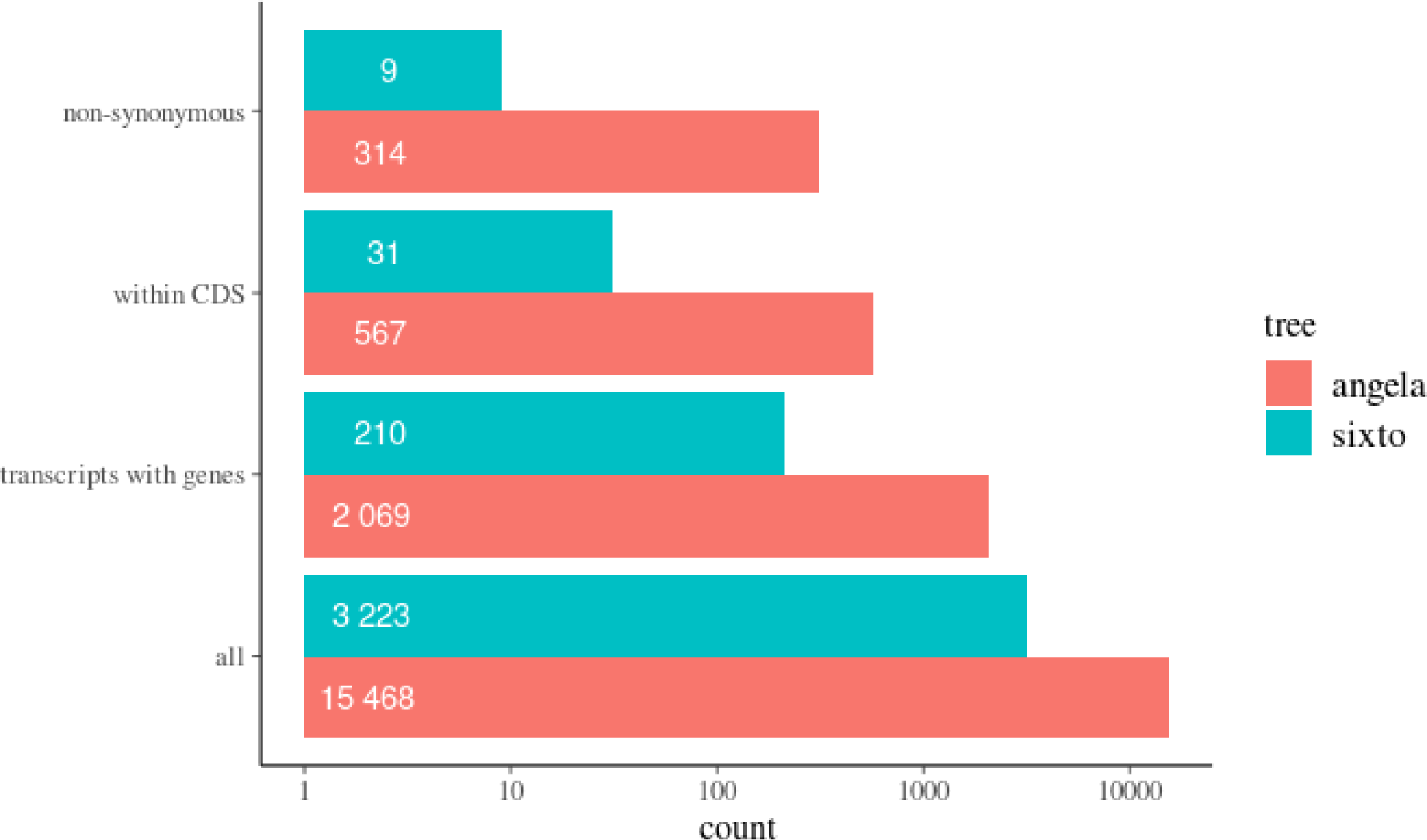
Mutation classified by functional impact with position and SNPeff (Cingolani et al., 2012) for Angela (red) and Sixto (blue). Mutations are classified into all, in a transcript with a coding DNA sequence, within the coding DNA sequence (CDS), and with at least one non-synonymous effect on the coding DNA sequences.

**Figure H3:**
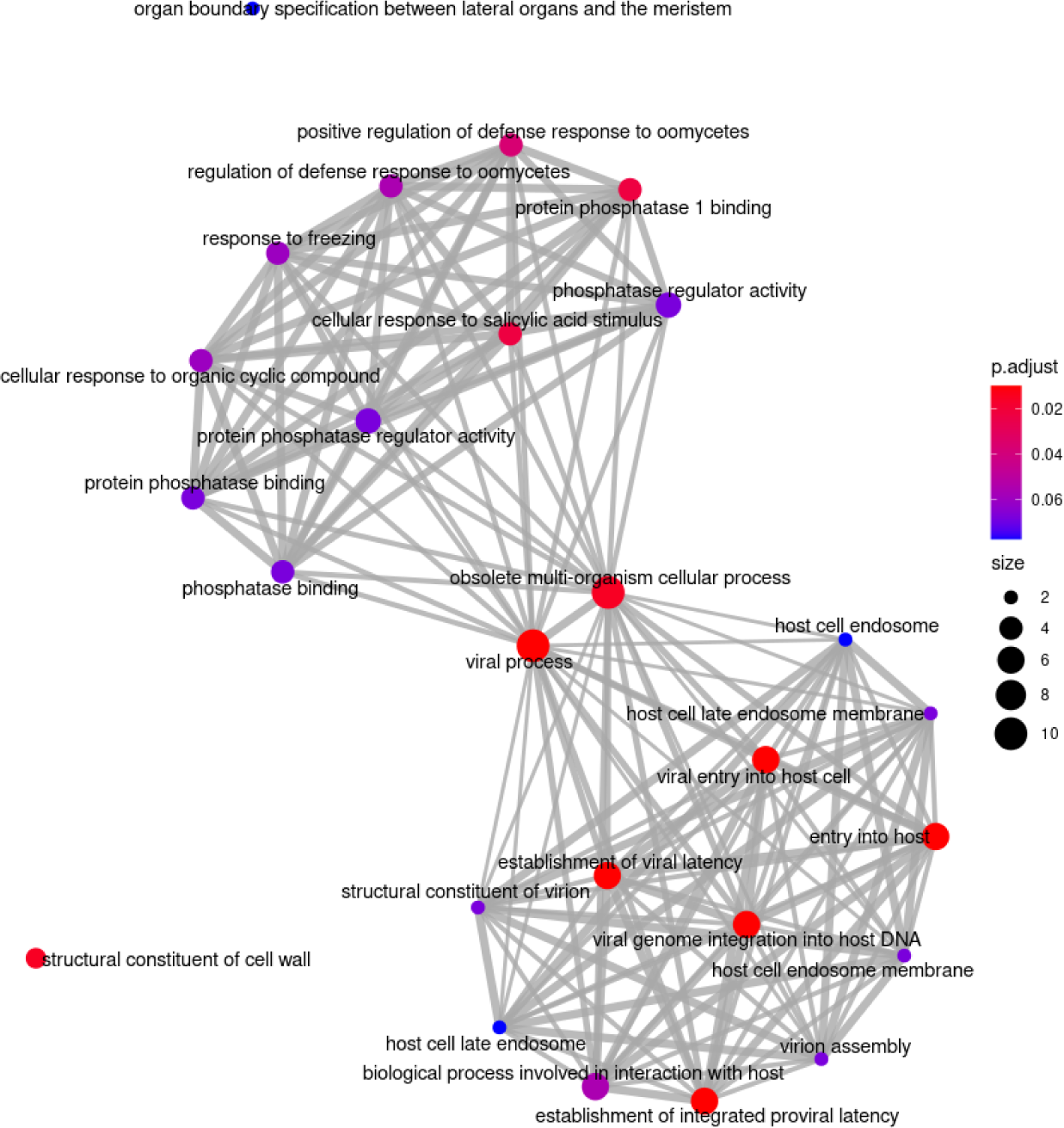
Enrichment map of the result of the over-representation of the enrichment analysis of gene sets with non-synonymous mutations in Angela. The enrichment in the ontology of genes carrying non-synonymous mutations was tested against all background gene ontologies. The colour represents the adjusted p-value and the size the number of connections to other enriched gene ontologies.

**Figure H4:**
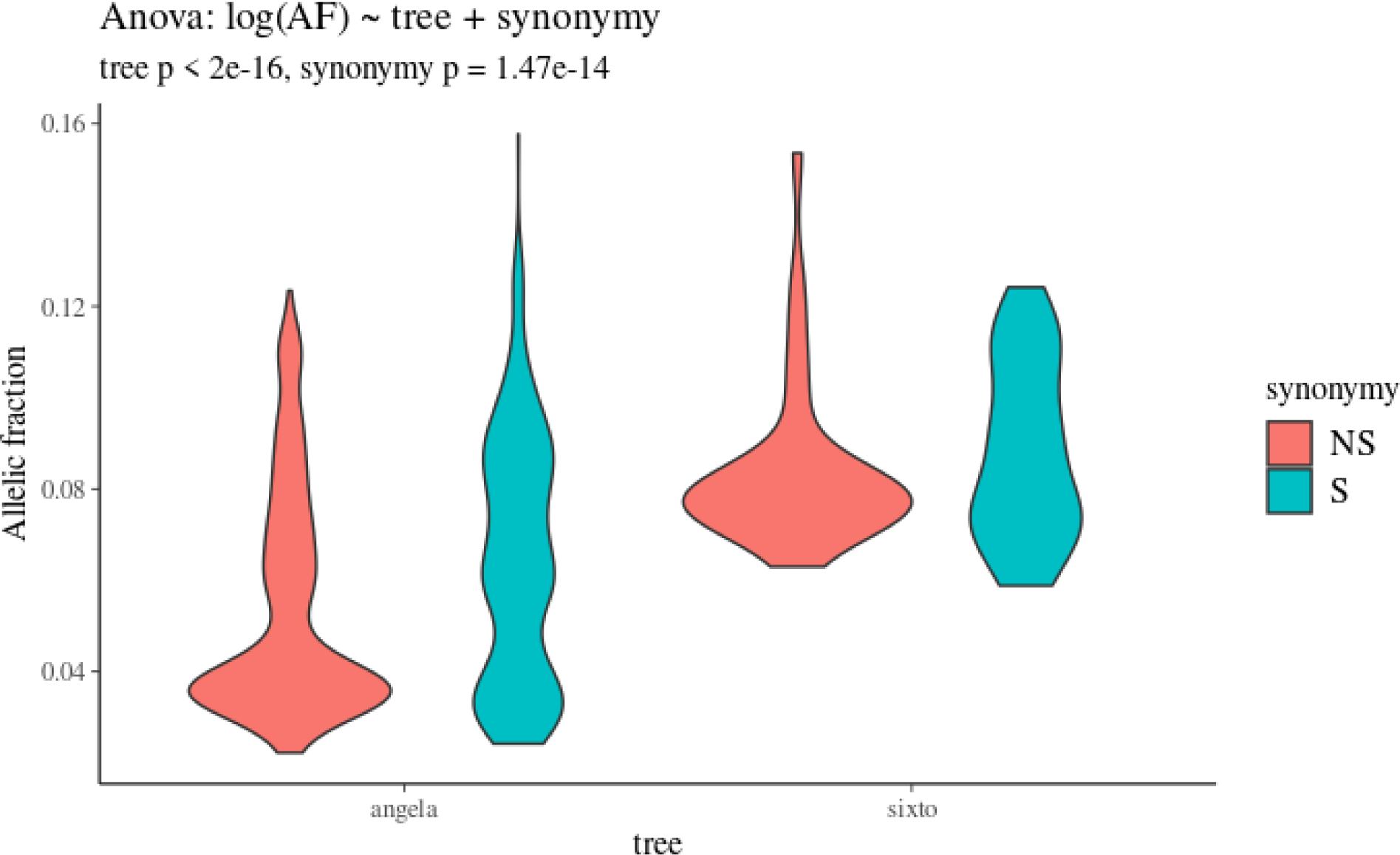
Allelic fractions of synonymous (S, blue) and non-synonymous (NS, red) mutations for Angela and Sixto. Type II ANOVA testing differences in the log allelic fraction with tree and synonymy revealed a significant negative effect (p=1.47*10-14) of non-synonymy on the allelic fraction. The effect of trees is methodologically expected due to the greater sequencing depth in Angela allowing detection of lower allelic fractions.

**Figure I1:**
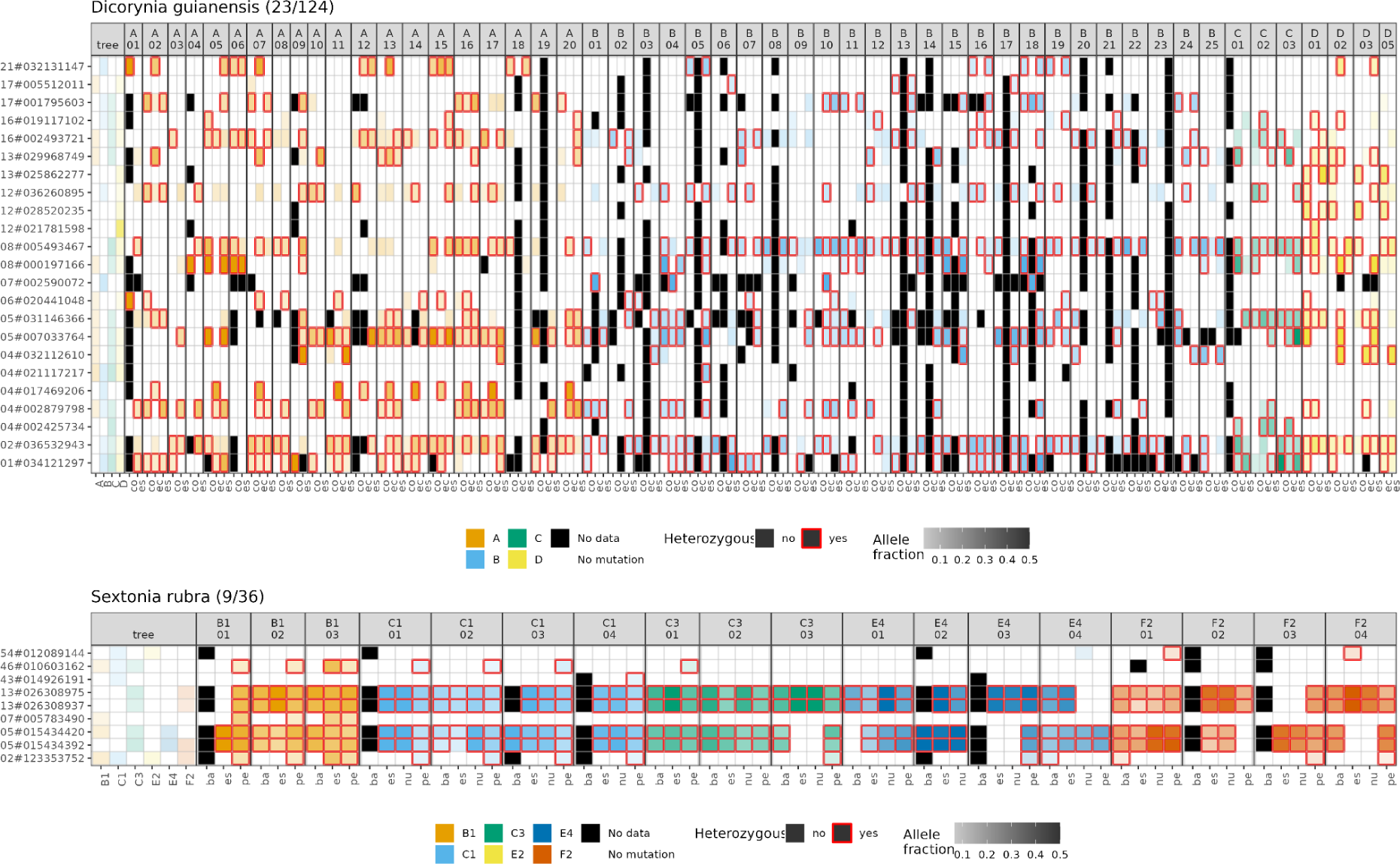
Candidate mutations transmission in fruit tissues for Dicorynia guianensis and Sextonia rubra. Every column represents either the tree crown or a fruit labelled with its branch and the number of the fruit. In each column mutation positions are given on the y-axis and the branch or fruit tissue on the x-axis. Colours indicate the branch origin, while white cells represent sequenced tissues with absence of mutations and black cells unsequenced tissues. The colour transparency represents the allelic fraction of the mutation from AF=0.1 with low-intensity colour to AF=0.5 with high intensity colour. The red border indicates sites called as heterozygous by GATK GenotypeGVCFs. The figure showcases the increase of allelic fraction in fruit tissues compared to the tree tissues from initial mutation detection. The different tissues are embryo sac (es), nucellus (nu), pericarp (pe), and fruit base (ba).

**Figure K1:**
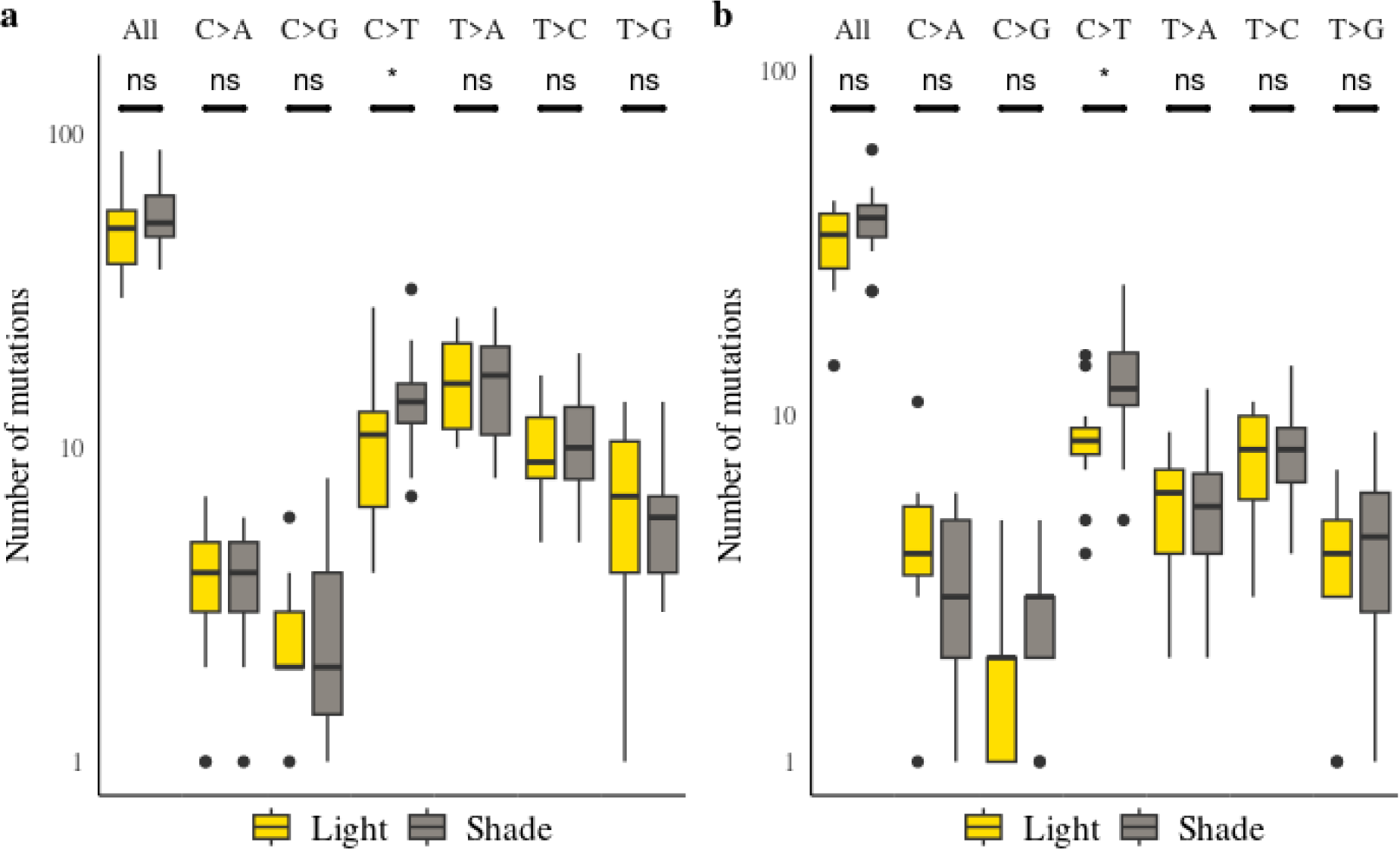
Distributions of somatic mutations with light for mutations passing the empirical variant score (EVS) filtering. The distributions of somatic mutations with light are similarly shown for the two tropical trees: the Dicorynia guianensis tree (a), and the Sextonia rubra tree (b). Different mutagens may cause specific mutation types, i.e., changing from base X to base Y (X>Y). The effect of light exposure on the accumulation of somatic mutations as a function of mutation type (X>Y) is represented in yellow and grey boxes. The yellow boxes represent the number of mutations accumulated in all leaves of light exposed branches and the grey boxes in all leaves of shaded branches. Boxplots show the median (centre line), upper and lower quartiles (box limits), 1.5x interquartile range (whiskers), and outliers (points). The “ns” labels indicate non-significant differences in Student’s T-tests (two-sided). Mutation types include all mutations and all types of transitions and transversions. The y-axis has been scaled logarithmically to facilitate reading of low values.

**Figure K2:**
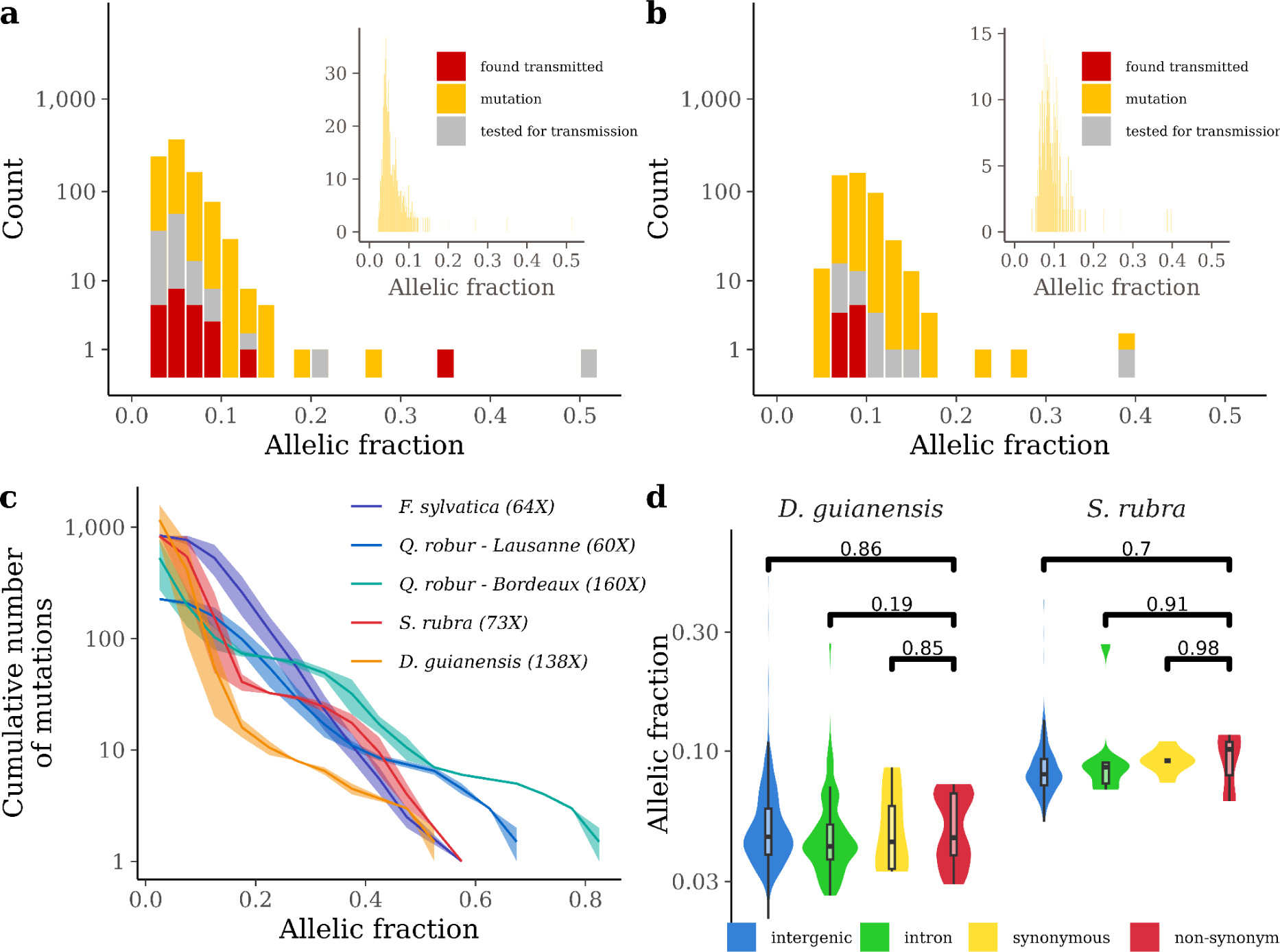
Allelic fractions of somatic mutations among trees and among genomic elements for mutations passing the empirical variant score (EVS) filtering. Histogram of allelic fractions of mutations detected in the crown of the two tropical trees: the Dicorynia guianensis tree (a), and the Sextonia rubra tree (b). The main histogram shows the allelic fractions of the somatic mutations using a bin of 0.02 and a log-transformed count with the mutations detected in the crown in yellow, the mutations tested for transmission in grey, and the mutations found transmitted to the embryos in red. The inner histogram shows the allelic fractions of the somatic mutations using a bin of 0.001 and a natural count. (c) Cumulative number of somatic mutations per branch with decreasing allelic fraction for five trees reanalysed with the same pipeline. The five trees include the two tropical trees studied, the Dicorynia guianensis tree in orange and the Sextonia rubra tree in red, and three temperate trees, two pedunculate oaks Quercus robur L. from Bordeaux in green and Lausanne in blue and a tortuous phenotype of common beech Fagus sylvatica L. in purple. All trees were analysed with the same pipeline (see methods) but were sequenced with a different depth indicated in brackets. The line represents the median value while the area represents the minimum and maximum values on the 2 to 10 branches per tree. (d) Comparisons of allelic fractions for non-synonymous mutations in red with synonymous mutations in yellow, intronic mutations in green and intergenic mutations in blue for the two tropical trees: the Dicorynia guianensis tree (left panel), and the Sextonia rubra tree (right panel). Boxplots show the median (centre line), upper and lower quartiles (box limits), 1.5x interquartile range (whiskers), and outliers (points). The p-value above the bars indicates the significance of the Student’s T-test (two-sided) for the pairs of groups.

**Table A1:** Characteristics of the 15 candidate species: **Family**, **genus**, and **species** names are tabulated; **An. rings**: The occurrence of annual rings either in the species, or in other species in the genus; **Rep. phen**.: if the reproductive phenology is known, we indicate if its is annual (A), biennial (B), or supra-annual (S-A); **Arch. Model:** the typical architecture model of the species; **GS (Mbp):** the size of the 1C haploid genome in mega base pairs (Mbp); **Ind**.: sampling location and individual identifier at the Paracou Station (P: Parcelle; C: carré; Ind: individual).

| Family | Genus | Species | ring | Rep. phen. | Arch. Model | GS (Mbp) | Ind. |
| --- | --- | --- | --- | --- | --- | --- | --- |
| Bignoniaceae | Handroanthus | serratifolius (Vahl) S.O. Grose | S |  | Koriba (Hot, 1978) | 572 | P_6, C_2, Ind_431 |
| Caryocaraceae | Caryocar | glabrum (Aubl.) | S | A | Scaronne (Vester & Cleef, 1998) | 899 | P_6, C_2, Ind_716 |
| Clusiaceae | Moronobea | coccinea (Aubl.) | S | A; B | Massart (Keller, 1994) | 1508 | P_6, C_2 Ind_776 |
| Clusiaceae | Platonia | insignis (Mart.) | S | B | Massart (Hot, 1978) | 1748 | P_14, C_3, Ind_112 |
| Clusiaceae | Symphonia | globulifera (L.) | S | A | Massart (Hot, 1978; Barthélémy, 1988) | 1836 | P_6, C_4, Ind_2941 |
| Dichapetalaceae | Tapura | capitulifera (Baill.) | G |  | Cook (Hot, 1978) | 948 | P_6, C_2, Ind_2919 |
| Fabaceae | Dicorynia | guianensis (Amshoff) | S | S-A | Troll (Drénou, 1988) | 691 | P_6, C_2, Ind_898 |
| Fabaceae | Vouacapoua | americana (Aubl.) | S | S-A | Troll (Hot, 1978) | 1028 | P_6, C_2, Ind_387 |
| Goupiaceae | Goupia | glabra (Aubl.) | S |  | Roux (Hot, 1978) | 482 | P_6, C_2, Ind_774 |
| Lauraceae | Sextonia | rubra (Mez) van der Werff | no |  | Aubréville (Laurans & Vincent 2016) | 1354 | P_6, C_3, Ind_2353 |
| Lecythidaceae | Eschweilera | coriacea (DC.) S.A. Mori | no |  | Troll (Obs. Caraglio & Nicolini) | 1040 | P_6, C_2, Ind_876 |
| Myristicaceae | Virola | michellii (Heckel) | G |  | Massart (Hot, 1978) | 1051 | P_6, C_2, Ind_2812 |
| Salicaceae | Laetia | procera (Poepp.) Eichler | no |  | Roux (Drénou, 1994) | 419 | P_6, C_2, Ind_790 |
| Sapotaceae | Pradosia | cochlearia (Lecomte) T.D.Penn. | no | A | Aubréville (Laurans & Vincent 2016) | 983 | P_6, C_2, Ind_766 |
| Vochysiaceae | Qualea | rosea (Aubl.) | no |  | Massart (Sanoja, 1992) | 1179 | P_6, C_2, Ind_607 |

**Table C1:**
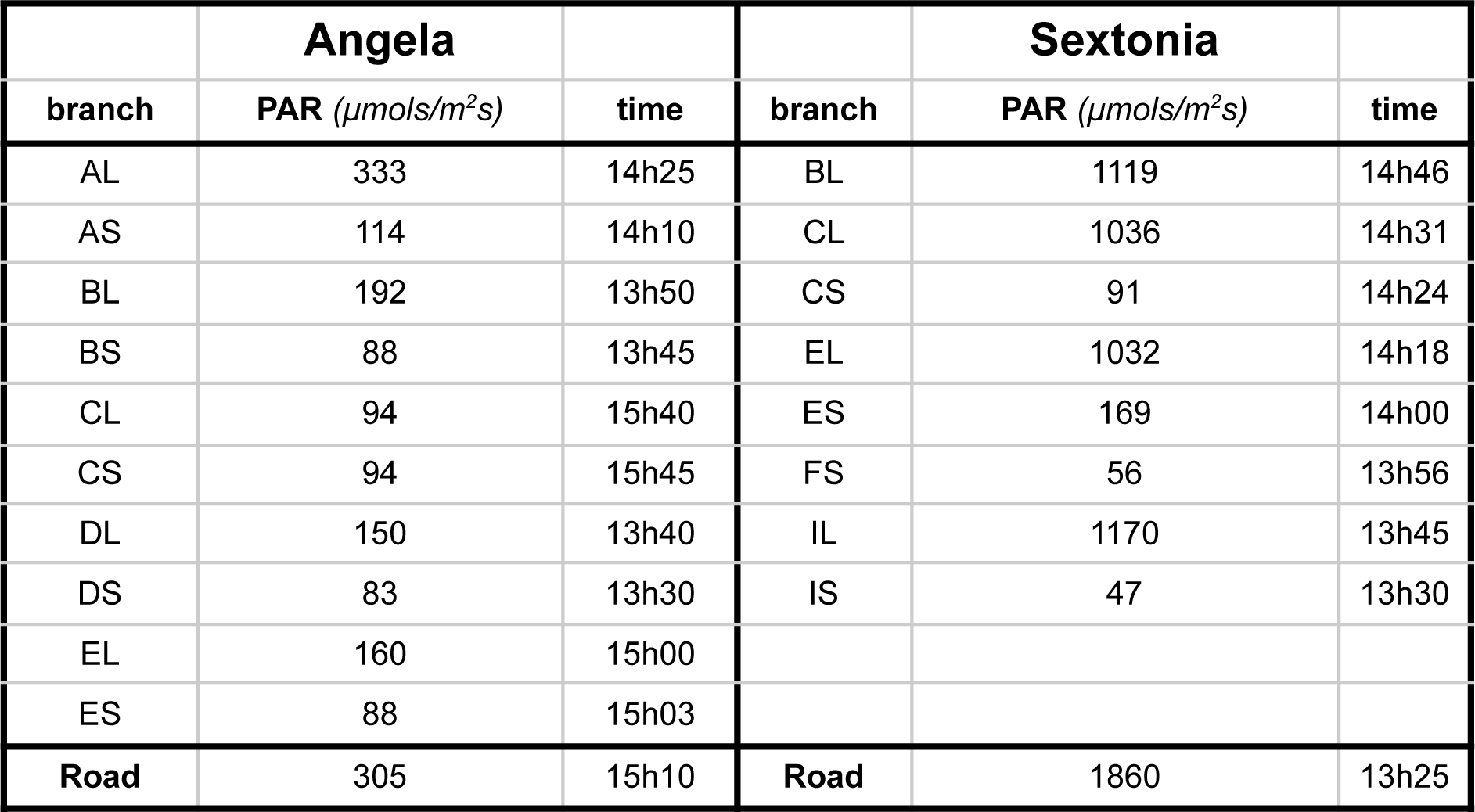
Direct incident light at sampling points in Angela and Sixto: **branch**: indicates the sampling point as mapped in Fig B3 and Fig B4; **PAR**: Photosynthetically active radiation measurement at the sampling point (μmols/m^2^s); **time**: the time the measurement was taken; **road**: indicates the PAR measurement at the nearest road (open space), maximum of 100m away from tree.

**Table C2:** Radiation balance at the ten sampling points in Angela. Table C2 summarises lidar and genomic identifiers for branches with corresponding light condition assessed by climbers and the transmittance estimated from the radiation balance. The standard deviation of transmittance in parenthesis is given for a distance of 0.5m from the identified sampling point.

| **ID lidar** | **ID genomic** | **Condition** | **Transmittance**<br>**(standard**<br>**deviation)** | **Height**<br>**above**<br>**ground** |
| --- | --- | --- | --- | --- |
| B31 | AL | light | 0.34 (0.09) | 44.31 |
| B31 | AS | shadow | 0.08 (0.02) | 44.76 |
| B32 | BL | light | 0.84 (0.22) | 46.44.7 |
| B32 | BS | shadow | 0.15 (0.04) | 38.08 |
| B4 | CL | light | 0.17 (0.04) | 37.18 |
| B4 | CS | shadow | 0.08 (0.02) | 33.6 |
| B2 | DL | light | 0.80 (0.08) | 43.25 |
| B2 | DS | shadow | 0.16 (0.04) | 38.95 |
| B1 | EL | light | 0.43 (0.01) | 31.45 |
| B1 | ES | shadow | 0.09 (0.04) | 28.97 |

**Table D1:**
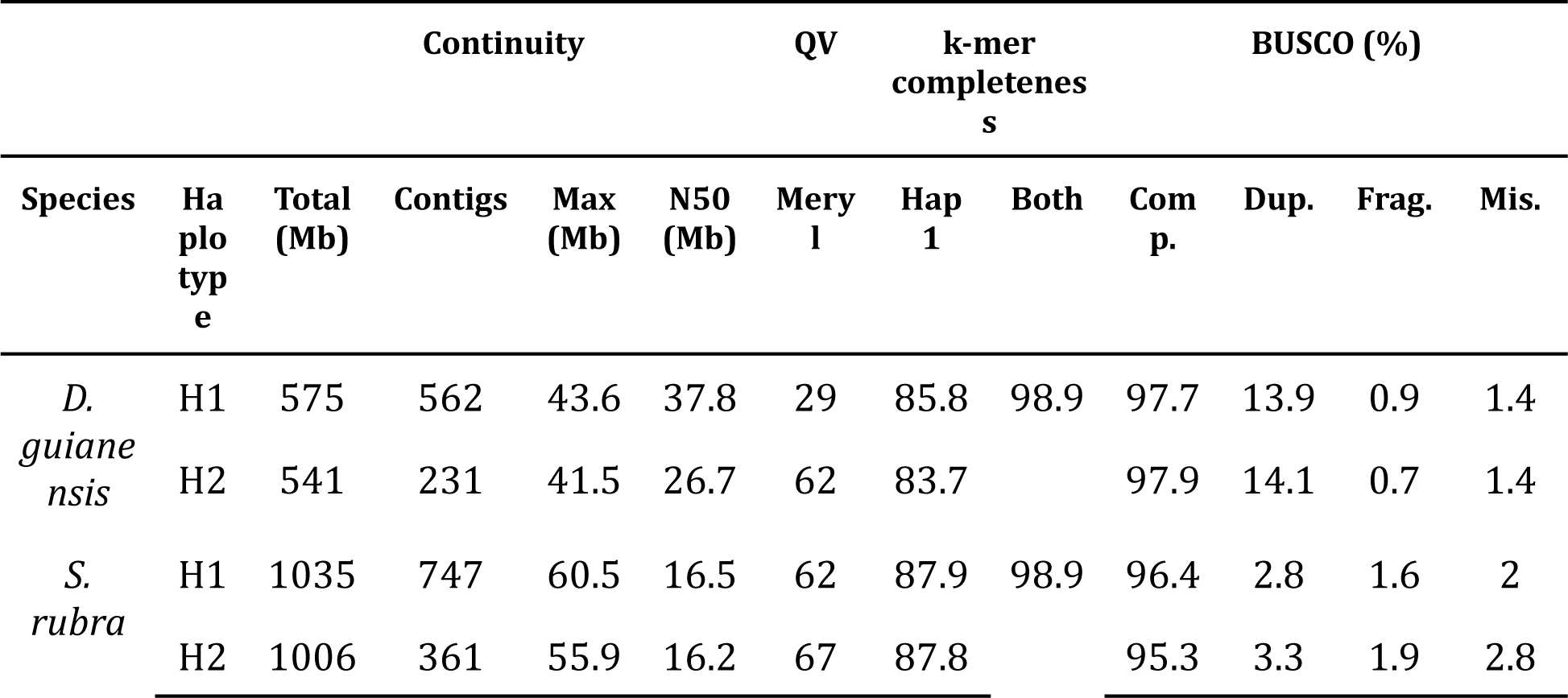
Quality check metrics for Dicorynia guianensis and Sextona rubra genomes. QV and completeness were measured with Merqury (Rhie et al., 2020). Assembly consensus quality values (QV) were estimated using k-mer analysis, which represents a log scaled probability of error for the consensus base calls. Genome completeness was assessed by BUSCO analyses (Seppey e al., 2019).

**Table F1:** Correlations of Angela mutations among sample point pairs. See figure F1 for labels.

|  | AL | AS | BL | BS | CL | CS | DL | DS | EL | ES |
| --- | --- | --- | --- | --- | --- | --- | --- | --- | --- | --- |
| AL | 1.00 | 0.01 | -0.02 | 0.01 | 0.00 | 0.02 | -0.05 | -0.03 | -0.01 | 0.00 |
| AS | 0.01 | 1.00 | -0.03 | 0.09 | -0.01 | -0.04 | 0.07 | -0.04 | -0.01 | -0.01 |
| BL | -0.02 | -0.03 | 1.00 | -0.03 | 0.01 | -0.02 | -0.02 | -0.02 | -0.03 | 0.04 |
| BS | 0.01 | 0.09 | -0.03 | 1.00 | 0.01 | -0.01 | 0.03 | -0.03 | 0.00 | 0.00 |
| CL | 0.00 | -0.01 | 0.01 | 0.01 | 1.00 | 0.01 | -0.05 | -0.03 | -0.01 | -0.05 |
| CS | 0.02 | -0.04 | -0.02 | -0.01 | 0.01 | 1.00 | -0.03 | -0.01 | 0.02 | 0.00 |
| DL | -0.05 | 0.07 | -0.02 | 0.03 | -0.05 | -0.03 | 1.00 | -0.07 | 0.00 | 0.02 |
| DS | -0.03 | -0.04 | -0.02 | -0.03 | -0.03 | -0.01 | -0.07 | 1.00 | -0.02 | 0.00 |
| EL | -0.01 | -0.01 | -0.03 | 0.00 | -0.01 | 0.02 | 0.00 | -0.02 | 1.00 | -0.02 |
| ES | 0.00 | -0.01 | 0.04 | 0.00 | -0.05 | 0.00 | 0.02 | 0.00 | -0.02 | 1.00 |

**Table F2:** Correlations of Sixto mutations among sample point pairs. See figure F1 for labels.

|  | B1L | C1S | C3L | E2L | E4S | F2S | I1L | I1S |
| --- | --- | --- | --- | --- | --- | --- | --- | --- |
| B1L | 1.00 | -0.09 | -0.05 | -0.08 | -0.02 | -0.07 | -0.05 | -0.05 |
| C1S | -0.09 | 1.00 | -0.06 | -0.05 | -0.02 | -0.05 | -0.06 | -0.03 |
| C3L | -0.05 | -0.06 | 1.00 | -0.03 | -0.01 | -0.03 | -0.06 | -0.01 |
| E2L | -0.08 | -0.05 | -0.03 | 1.00 | -0.04 | -0.07 | -0.05 | -0.03 |
| E4S | -0.02 | -0.02 | -0.01 | -0.04 | 1.00 | -0.02 | -0.06 | -0.04 |
| F2S | -0.07 | -0.05 | -0.03 | -0.07 | -0.02 | 1.00 | -0.07 | -0.06 |
| I1L | -0.05 | -0.06 | -0.06 | -0.05 | -0.06 | -0.07 | 1.00 | -0.02 |
| I2S | -0.05 | -0.03 | -0.01 | -0.03 | -0.04 | -0.06 | -0.02 | 1.00 |

